# Identification of cysteine metabolism regulator (CymR)-derived pentapeptides as nanomolar inhibitors of *Staphylococcus aureus* O-acetyl-ʟ-serine sulfhydrylase (CysK)

**DOI:** 10.1101/2024.09.19.614015

**Authors:** Jordan L. Pederick, Bethiney C. Vandborg, Amir George, Hannah Bovermann, Jeffrey M. Boyd, Joel S. Freundlich, John B. Bruning

## Abstract

The conditionally essential pathway of bacterial cysteine biosynthesis is gaining traction for the development of antibiotic adjuvants. Bacterial cysteine biosynthesis is generally facilitated by two enzymes possessing O-acetyl-ʟ-serine sulfhydrylase (OASS) activity, CysK and CysM. CysK enzymes can also form functional complexes with other proteins that regulate cysteine metabolism. In *Staphylococcus aureus* there exists a single OASS homologue, herein termed *Sa*CysK. Knockout of *Sa*CysK was found to increase sensitivity to oxidative stress, making it a relevant target for inhibitor development. *Sa*CysK forms two functional complexes via interaction with the preceding enzyme in the pathway serine acetyltransferase (CysE) or the transcriptional regulator of cysteine metabolism (CymR). These interactions occur through the insertion of a C-terminal peptide of CysE or CymR into the active site of *Sa*CysK, inhibiting OASS activity, and therefore represent an excellent starting point for developing *Sa*CysK inhibitors. Here we detail the characterization of CysE and CymR-derived C-terminal peptides as inhibitors of *Sa*CysK. First, interactions between CysE or CymR-derived C-terminal decapeptides and *Sa*CysK were assessed by X-ray crystallography. While both peptides occupied the active site of *Sa*CysK, the alternate sidechains of the CymR decapeptide formed more extensive interactions. Surface plasmon resonance binding assays and *Sa*CysK inhibition assays revealed that the CymR decapeptide bound to *Sa*CysK with nanomolar affinity (K_D_ = 25 nM) and inhibited *Sa*CysK activity (IC_50_ = 180 nM), making it a promising lead for the development of *Sa*CysK inhibitors. To understand the determinants of this high affinity interaction the structure-activity relationships of 16 rationally designed peptides were also investigated. This identified that the C-terminal pentapeptide of CymR alone facilitates the high affinity interaction with *Sa*CysK, and that subtle structural modification of the pentapeptide is possible without impacting potency. Ultimately, this work has identified CymR pentapeptides as a promising scaffold for the development of antibiotic adjuvants targeting *Sa*CysK.

**Author summary:** There is increasing interest in the investigation of non-essential pathways including bacterial cysteine metabolism for developing antibiotic adjuvants. Within this pathway the O-acetyl-ʟ-serine sulfhydrylase (OASS) enzymes CysK and CysM have been a focus. As such, the OASS enzyme of *Staphylococcus aureus*, *Sa*CysK, gained our interest. Previous efforts to inhibit CysK enzymes have mimicked the interaction between CysK and the C-terminus of serine acetyltransferase (CysE) which occurs inside the CysK active site and inhibits OASS activity. CysE peptides have only moderate potency, typically binding with micromolar affinity. In *S. aureus* another complex forms between *Sa*CysK and a transcriptional regulator CymR, but the ability of CymR peptides to inhibit CysK enzymes has not been investigated. We noticed there is variation between the C-terminus of CysE and CymR, suggesting that CymR peptides make distinct interactions with *Sa*CysK and may be superior inhibitors. Here we characterized CysE and CymR peptides as *Sa*CysK inhibitors. We found CymR peptides make more extensive molecular interactions with *Sa*CysK and bind with higher affinity, being the most potent peptide inhibitors of a CysK enzyme to date. A CymR pentapeptide is the minimal length required for this potency and provides a promising scaffold for developing antibiotic adjuvants targeting *Sa*CysK.

## Introduction

Antibiotic resistance continues to represent one of the greatest threats to modern medicine. The most recent systematic analysis performed by the Global Research on Antimicrobial Resistance project determined that 1.27 million deaths globally were caused by antibacterial resistance in 2019, and that antibiotic resistance contributed to a further 3.7 million deaths [1]. The high risk ESKAPE pathogens *(Enterococcus faecium, Staphylococcus aureus, Klebsiella pneumoniae, Acinetobacter baumannii, Pseudomonas aeruginosa,* and *Enterobacter species*) were the major group responsible. Furthermore, the increasing prevalence of multi-drug resistant bacteria has critically challenged our health care system. It is becoming clear that the traditional antibiotic development pipeline will be inadequate to combat acquisition of antibiotic resistance, as these efforts have largely focussed on the development of existing antibiotic classes with known mechanisms of resistance [2–4]. As such, it has been proposed that this issue can be addressed, in part, through focussing on the development of non-traditional or innovative agents that work by new mechanisms, and which can complement or enhance the effects of current antibiotics. Antibiotic adjuvants are one such example [5–7]. These agents possess limited or no antimicrobial effects alone but can illicit synergistic effects with existing antibiotics to sensitize the pathogen to certain antibiotics and even reduce or overcome antibiotic resistance. This outcome may be produced by a variety of mechanisms; therefore, the implementation of antibiotic adjuvants represents a versatile approach to mitigate resistance against current antibiotics.

The promise of adjuvant therapy in combatting bacterial pathogens has led to a recent interest in the investigation of non-essential pathways that hold potential as targets for antibacterial adjuvant development [5–7]. The signaling and regulatory network tied to bacterial cysteine biosynthesis is one example that has received substantial attention [8–10]. Although cysteine can be obtained from the mammalian infection environment, there is evidence that halting the ability of bacteria to produce their own cysteine can lead to decreased bacterial fitness and increased sensitivity to antibiotics [8,11–13]. This is consistent with the important roles of cysteine in protein synthesis, as a building block for important sulfur-containing molecules such as methionine, coenzyme A, Fe-S clusters and low molecular weight redox buffers such as glutathione, as well as the intrinsic antioxidant activity of cysteine that contributes to redox homeostasis [14–16]. In bacteria cysteine biosynthesis is facilitated by the activity of the pyridoxal phosphate (PLP)-dependent O-acetyl-ʟ-serine sulfhydrylase (OASS) enzymes, which catalyse the reaction of O-acetyl-ʟ-serine (OAS) and bisulfide (Fig 1A). OAS is produced by the activity of serine acetyltransferase (CysE) while bisulfide is produced through the reductive sulfur assimilation pathway or through degradation of cysteine [11,17–19].

**Fig 1.**
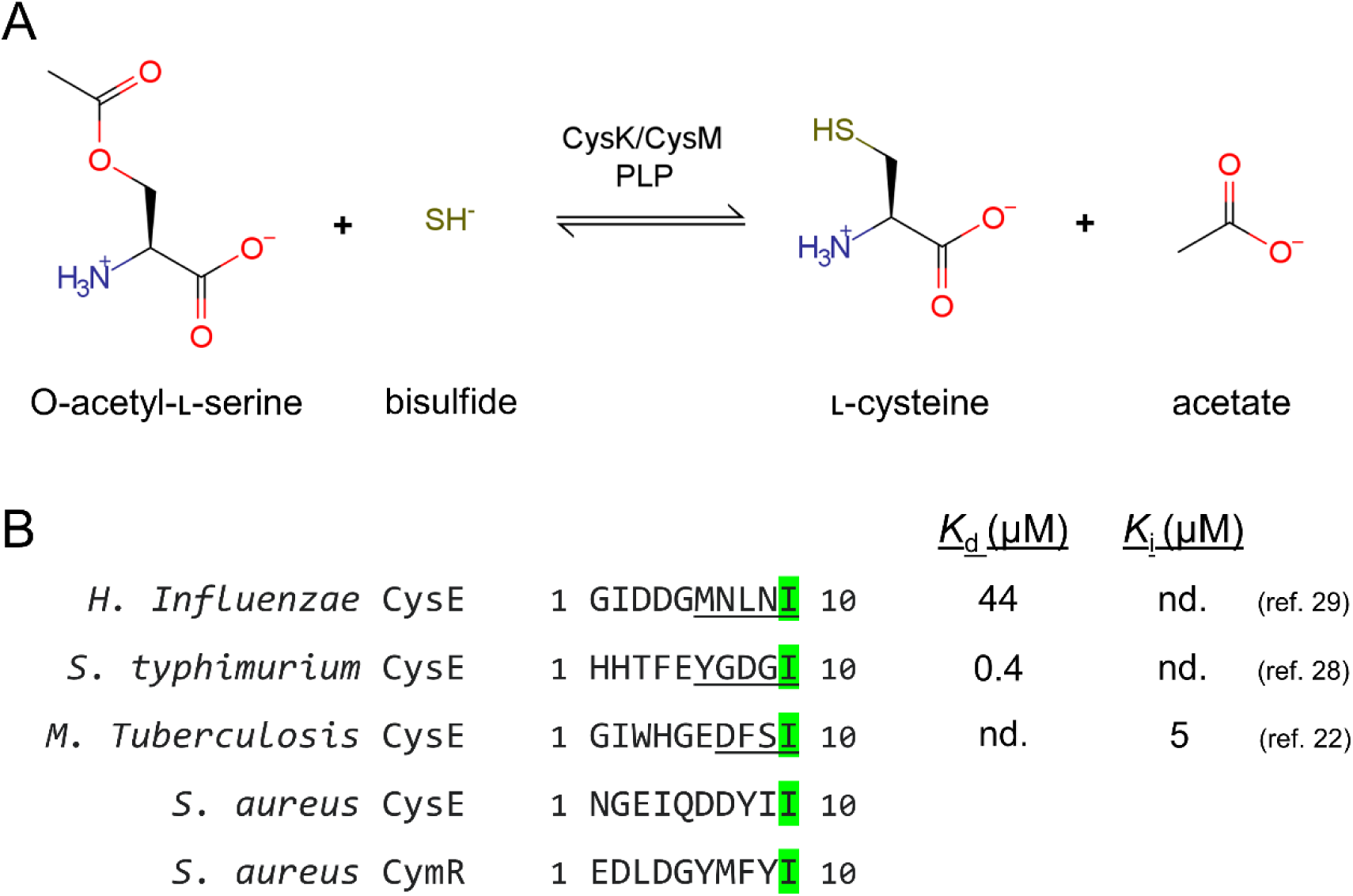
CysK enzymes possess multiple functions. **A)** Reaction catalyzed by the O-acetyl-ʟ-serine sulfhydrylase activity possessed by both CysK and CysM holo enzymes. **B)** The C-terminal regions of CysE and CymR enzymes that form functional complexes with CysK. The C-terminal isoleucine is conserved in all cases, indicated by the green shading. The binding affinities or inhibition constants for the previously characterized interactions of *H. influenzae*, *S. typhimurium* and *M. tuberculosis* CysE-derived peptides (indicated by underlined sequence) with the corresponding CysK enzyme are provided. The corresponding regions of *S. aureus* CysE and CymR are shown for comparison.

In most bacteria there exist two isoforms referred to as OASS-A (CysK) and OASS-B (CysM). The CysK and CysM terminology will be adopted herein. This assignment was originally based on the observation that CysM can use thiosulfate as an alternate sulfur donor while CysK cannot [20–22]. A more distinguishing property was subsequently identified with CysK being found to form functional complexes with specific protein partners that function to regulate cysteine metabolism (Fig 1B) [23]. The same complexes cannot form with CysM due to structural differences between the isoforms. The first of these characterized was the formation of a complex between CysK and CysE, termed the cysteine synthase complex (CSC). CSC formation was found to function as a wide-spread sensory and regulatory mechanism in bacterial cysteine biosynthesis, with structural characterisation of the interaction revealing that the C-terminus of CysE occupies the active site of CysK to control cysteine biosynthesis through competitive inhibition [24,25]. In the CSC CysE remains functional and produces OAS, resulting in the eventual dissociation of the CSC by the accumulation of OAS which competes with CysE for binding to CysK. Later, a second complex was identified to form between CysK and a transcriptional regulator of cysteine metabolism termed CymR. Intriguingly, CymR has only been observed in Gram-positive bacteria to date and represents a unique means of controlling cysteine metabolism [26–29]. It is suspected that the CysK:CymR complex forms similarly to the CSC through the C-terminus of CymR. Formation of the CysK-CymR complex results in stabilisation of CymR allowing it to bind to promoter regions of target genes involved in cysteine metabolism, repressing transcription of these genes. Under conditions of stress where cysteine is depleted and OAS accumulates this complex dissociates through the same competitive action of OAS, derepressing the CymR regulon. Due to these important roles in bacterial cysteine biosynthesis the inactivation of CysK and CysM has been pursued as a target for inhibitor design.

Efforts to date have mostly focussed on the development of inhibitors targeting CysK and CysM of Gram-negative pathogens. This has led to the discovery and characterization of both small molecule and peptide inhibitors [30]. Initially, the natural interaction between CysE and CysK was used as a basis to develop CysE-derived peptide inhibitors [25,31,32]. These efforts resulted in the identification of pentapeptide inhibitors of *Salmonella typhimurium* and *Haemophilus influenzae* CysK, and a tetrapeptide inhibitor of *Mycobacterium tuberculosis* CysK, with binding affinities mostly in the micromolar range. While a successful proof-of-concept, these inhibitors possess two main drawbacks. First, the affinity of the CysK:peptide interactions remained significantly lower compared to the natural CysK:CysE interaction, which approaches sub-nanomolar affinity [33]. Furthermore, for studies on *S*. *typhimurium* and *H. influenzae*, the interaction of the peptides with CysM was comparatively weak, with the binding affinities measured being >100 µM. Given the relatively low affinity for CysK and that inhibition of both CysK and CysM activity is required to block cysteine biosynthesis in *S*. *typhimurium* and *H. influenzae,* development of these peptides has not progressed further. Following this, attention turned to the development of small molecule inhibitors. Currently, the most prominent inhibitors are small-molecule cyclopropane derivatives that act as dual inhibitors of *S. typhimurium* CysK and CysM with the most potent compound reported to bind with affinities of 28 and 490 nM against the respective targets [34–36]. These cyclopropane derivatives were found to act as an adjuvant with the antibacterial colistin versus *S. typhimurium*, *E. coli*, and *K. pneumoniae* [35]. This effect was validated in *S. typhimurium*, with a strain lacking CysK and CysM activity displaying the same phenotype. Importantly, this provides direct evidence that CysK and CysM are valid targets for antibacterial adjuvant development.

While Gram-negative pathogens have embodied the primary focus of CysK and CysM inhibitor development there is evidence that inhibition of OASS activity in the Gram-positive pathogen *S. aureus* has adverse effects on bacterial fitness. Unlike *S. typhimurium*, *S. aureus* possesses one CysK/CysM homologue, corresponding to SA0471 in *S. aureus* N315. SA0471 was originally assigned as CysM based on the observation that a knockout mutation of SA0471 prevented the ability of *S. aureus* SH1000 to use thiosulfate as a sulfur source, despite possessing closer sequence identity to CysK enzymes [37]. It was later identified that SA0471 is the direct binding partner of CymR in *S. aureus* and is required for binding of CymR to the promoter regions of target genes, providing more conclusive evidence that SA0471 corresponds to CysK [29]. Therefore, SA0471 will herein be referred to as *Sa*CysK. The *Sa*CysK knockout mutant was also found to display poor growth in cysteine-limited conditions and produced less cysteine than wild-type *S. aureus* SH1000 [37]. This was accompanied by increased sensitivity to tellurite, hydrogen peroxide, acid and diamide, and a reduced ability to recover from starvation in amino acid or phosphate limiting conditions. These findings suggest that cysteine biosynthesis plays an important role in the stress response and survival mechanisms of *S. aureus*, and places *Sa*CysK as a potential target for the development of antibiotic adjuvants.

Given these previous findings, we aimed to characterize *Sa*CysK as a target for inhibitor development. The natural interactions of *Sa*CysK with CysE or CymR presented as a favourable starting point toward this goal. Firstly, as *Sa*CysK is the only target, it simplifies this approach relative to the previous studies in Gram-negative pathogens which focussed on dual targeting of CysK and CysM. Secondly, the C-terminal region of *S. aureus* CymR possesses a unique amino acid sequence distinct from any previously studied CysK:CysE interactions (Fig 1B). While the CysK:CymR interaction has been experimentally validated, the structural basis of this interaction had not been determined in *S. aureus* or any other bacterial species. Therefore, we anticipated that further investigation of the CysK:CymR interaction of *S. aureus* may result in the identification of a novel peptide binding mode and potentially lead to the discovery of potent peptide inhibitors of *Sa*CysK. In this work, we employed a combination of X-ray crystallography, surface plasmon resonance binding assays, and *Sa*CysK inhibition assays to understand how *S*. *aureus* CysE- and CymR-derived peptides bind to and inhibit *Sa*CysK. Collectively, this identified CymR-derived pentapeptides as potent inhibitors of *Sa*CysK and provides a strong platform for the development of peptide or peptidomimetic inhibitors of *Sa*CysK that may possess antibiotic adjuvant activity.

## Results and Discussion

### Preliminary characterization of *Sa*CysK function and structure

As neither the function nor structure of *Sa*CysK had been previously validated, an initial characterization of the enzyme’s function and structure was performed using a combination of enzyme activity assays and X-ray crystallography. For these experiments *Sa*CysK was produced recombinantly in *E. coli* and obtained at a high purity as an adduct with PLP (Fig 2A and Fig 2B). Using this material, the ability of *Sa*CysK to produce ʟ-cys from O-acetyl-ʟ-serine (OAS) and sodium sulfide was assessed using a discontinuous colorimetric assay based on the method of Gaitonde [38]. This confirmed that *Sa*CysK is a functional OASS enzyme, possessing a *k*_cat_ of 540 ± 30 s^-1^ and *K*_M app_ for the OAS substrate of 12 ± 2 mM (Fig 2C). These kinetic parameters of *Sa*CysK are fairly consistent with those for CysK and CysM enzymes of other bacterial species supporting a comparable enzymatic function and reaction mechanism for *Sa*CysK [21,25,39–41].

**Fig 2.**
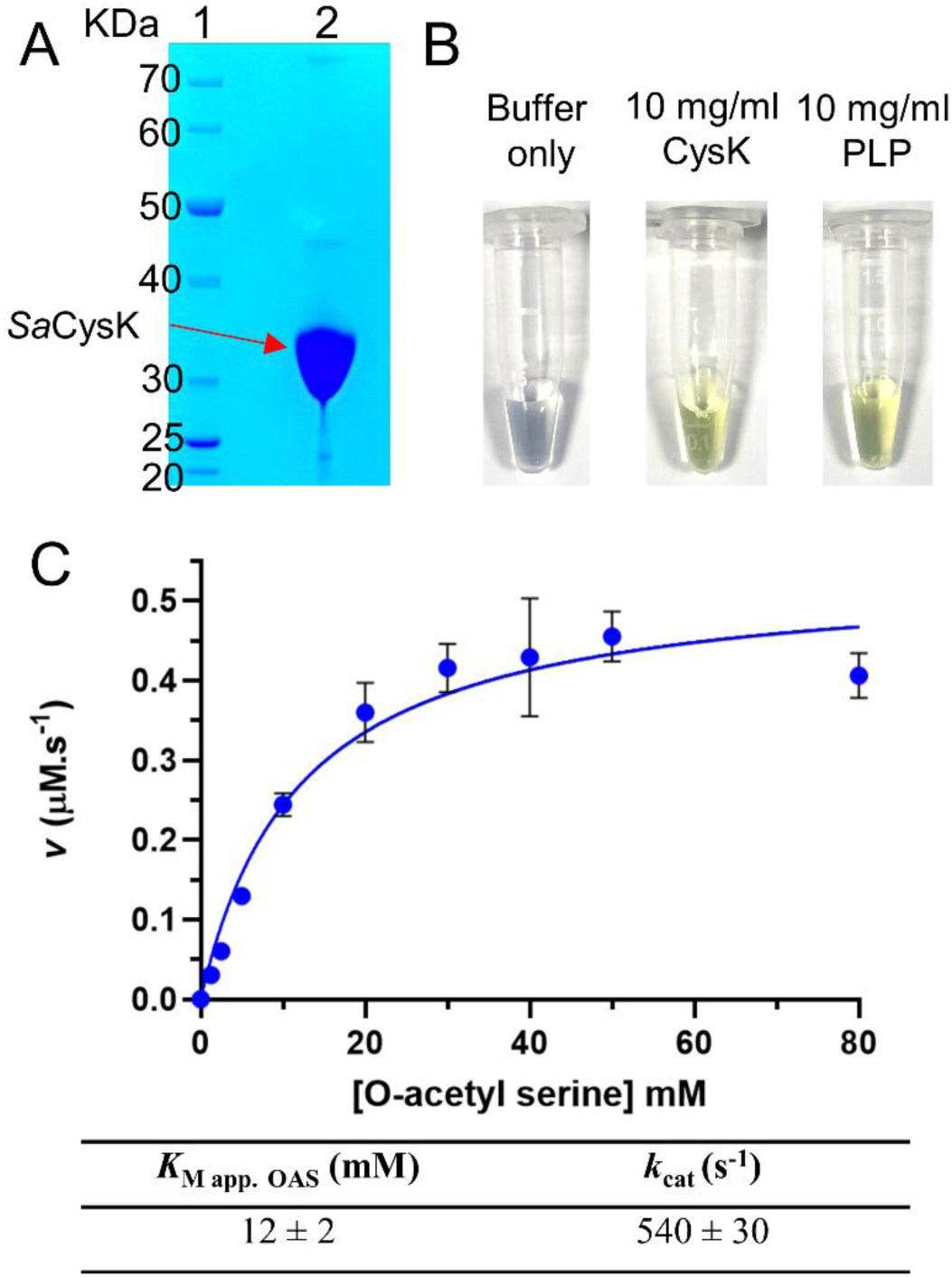
Isolation and functional characterization of *Sa*CysK. **A)** SDS-PAGE analysis of *Sa*CysK. Lane 1: NEB unstained protein ladder broad range (10 – 200 KDa), Lane 2: 5 µg purified *Sa*CysK. No contaminants were observed above or below the molecular weight range displayed on the gel. **B)** Appearance of purified *Sa*CysK. The intense yellow color indicated that *Sa*CysK was purified as a PLP-adduct. Buffer consisted of 20 mM Tris–HCl pH 8.0, 50 mM NaCl, and 1 mM DTT. **C)** Kinetic characterization of *Sa*CysK activity. Reactions were performed at 37 °C contained 50 mM Tris-HCl pH 7.5, 5 µM PLP, 1 mM sodium sulfide, 1 nM CysK and varying concentrations of OAS ranging between 1.25 and 80 mM. Data represents the mean ± standard deviation of three experiments.

The structure of the *Sa*CysK holo-enzyme was then elucidated by X-ray crystallography. Crystals of the *Sa*CysK holo-enzyme diffracted to high resolution, yielding a 1.9 Å dataset. *Sa*CysK forms the same homodimeric assembly as previously characterized CysK and CysM enzymes, with both active sites positioned on the same side of the homodimer (Fig 3A) [42]. The active site cavity of *Sa*CysK that is occupied by CysE and CymR is at the center of each monomer, being formed by Helix 4 and several loop regions – herein termed Loop 1, Loop 2, Loop 3, and Loop 4. Adjacent to this region is the PLP cofactor which forms a Schiff base linkage with the ε-amino group of K46. To confirm the correct assignment of the *S. aureus* cysteine synthase as *Sa*CysK these regions forming the active site cavity were compared with the previously characterized *S. typhimurium* CysK (*St*CysK) and CysM (*St*CysM) model isozymes, which possess 51 and 41% sequence identity to *Sa*CysK, respectively. While Helix 4 and Loops 1 – 3 were found to possess a high degree of conservation at the sequence and structural level for all three enzymes, Loop 4 is only conserved between *Sa*CysK and *St*CysK, but not for *St*CysM in which Loop 4 adopts an alternate structure (Fig 3B and Fig 3C). This is consistent with the previously identified role of Loop 4 residues in mediating the interaction of CysK enzymes with the C-terminus of CysE. Therefore, this functional and structural analysis has confirmed that the *S. aureus* cysteine synthase possesses the expected enzymatic activity and that assignment SA0471 is correctly assigned as *Sa*CysK.

**Fig 3.**
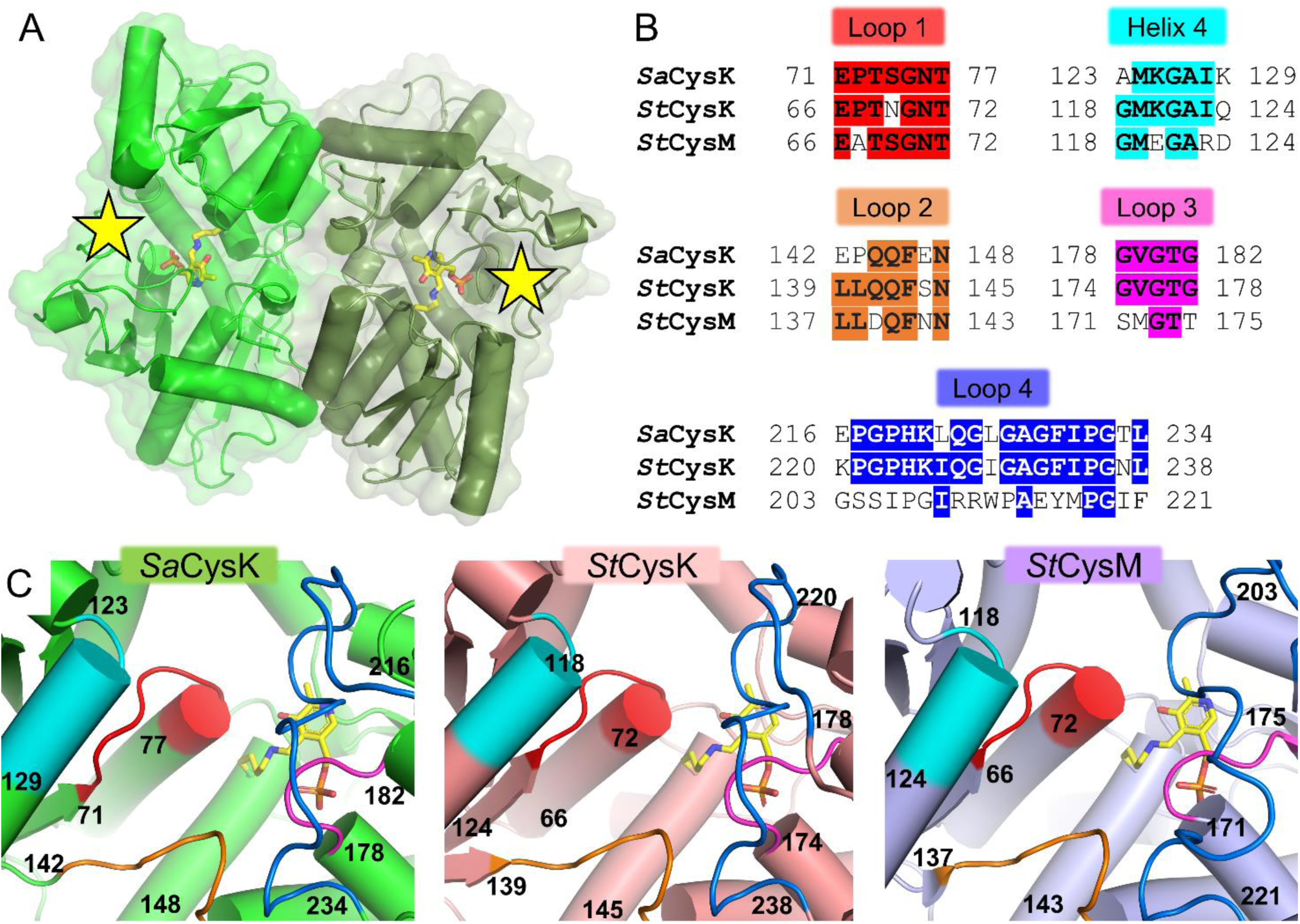
The overall fold and active site architecture of holo *Sa*CysK. **A)** The dimeric assembly of holo *Sa*CysK. The individual monomers of *Sa*CysK are shown in green and dark green, with the covalently bound PLP cofactor shown in yellow sticks. The active site cavity is indicated by the yellow stars. **B)** Sequence alignment of regions forming the active site in *Sa*CysK, *St*CysK and *St*CysM. Positions conserved between 2 or more sequences are indicated by colored shading. **C)** Structural superposition of the corresponding regions forming the active site in *Sa*CysK, *St*CysK (PDB: 1OAS) and *St*CysM (PDB: 2JC3). Loops 1 – 4 are shown as red, orange, pink and blue cartoons respectively, with Helix 4 shown in cyan.

### The C-terminus of CymR binds to *Sa*CysK with low nanomolar affinity through forming distinct molecular interactions

We next investigated the ability of *Sa*CysK to interact with the C-terminal region of the interacting proteins CysE and CymR using a surface plasmon resonance binding assay. For this purpose, a C-terminal decapeptide was used in each case, termed **CysE 10** (NH_2_–N_1_G_2_E_3_I_4_Q_5_D_6_D_7_Y_8_I_9_I_10_–COOH) and **CymR 10** (NH_2_–E_1_D_2_L_3_D_4_G_5_Y_6_M_7_F_8_Y_9_I_10_–COOH). In this experiment the histidine-tagged *Sa*CysK was employed as the ligand, being immobilized on a NiNTA loaded surface, with **CysE 10** or **CymR 10** used as the analytes. Binding responses for various concentrations of the **CysE 10** and **CymR 10** peptides were obtained and are displayed in Fig 4A. Consistent with previous studies employing similar short C-terminal peptides, both **CysE 10** and **CymR 10** were found to interact specifically with *Sa*CysK, indicated by the dose-dependent binding and saturation of the binding response at the highest concentrations assayed. To derive the binding affinity for each of the peptides this data was subjected to steady-state affinity analysis using the 1:1 binding model (Fig 4B). For **CysE 10** the determined binding affinity of 9 ± 1 µM was consistent with the affinities reported for previously characterized CysK:CysE peptide [25,31,32]. In contrast, the binding affinity of **CymR 10** was determined as 25 ± 5 nM, a remarkable ∼400-fold higher affinity interaction than measured for **CysE 10**. Notably, this places **CymR 10** as the highest affinity peptide interaction for a CysK enzyme reported in the literature to date.

**Fig 4.**
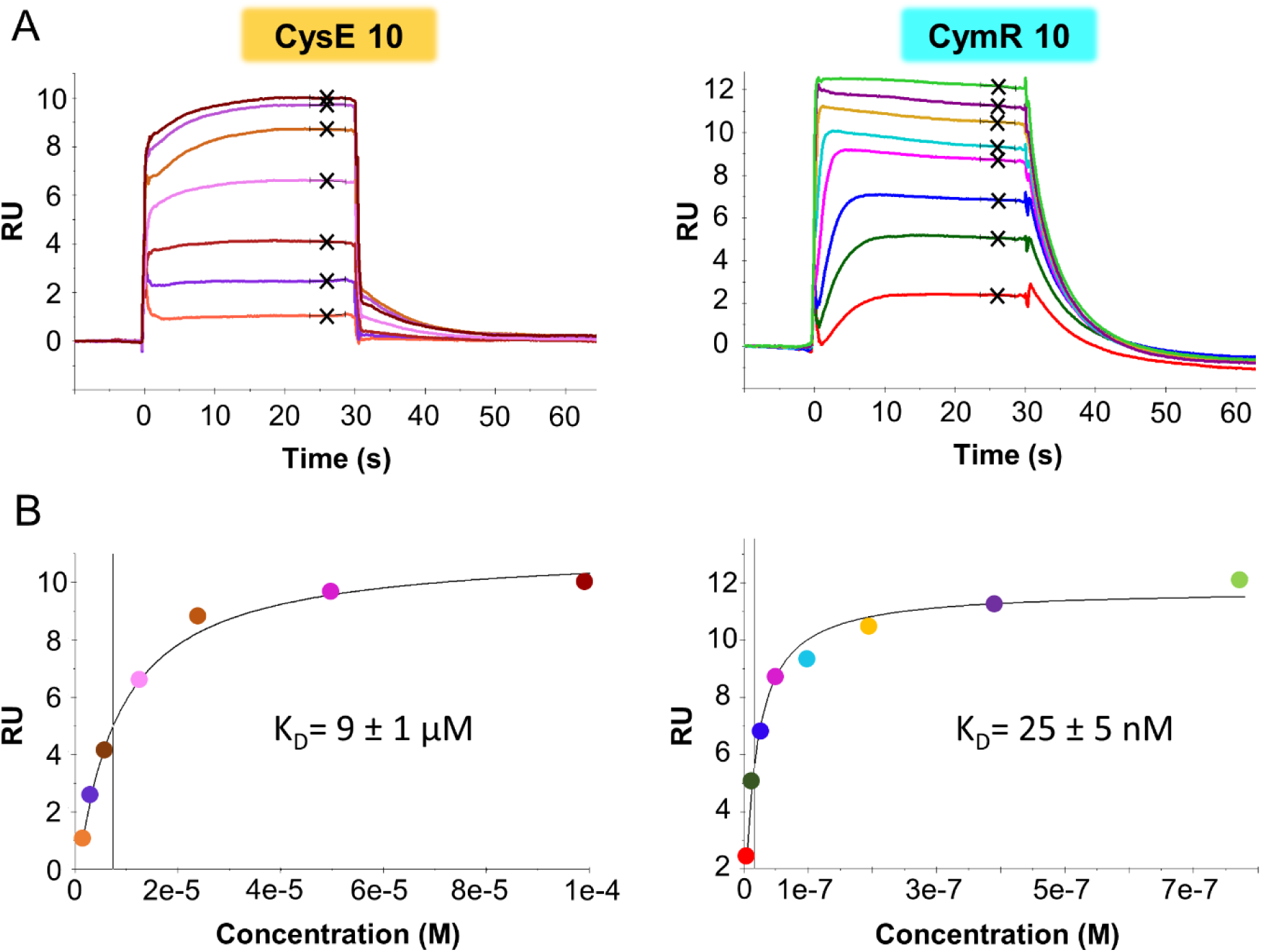
Characterizing the binding of CysE 10 and CymR 10 to *Sa*CysK by surface plasmon resonance (SPR). **A)** Representative sensorgrams detailing the specific, dose-dependent binding of **CysE 10** (left) and **CymR 10** (right) to *Sa*CysK. The steady-state binding responses used for calculation of the binding affinity (K_D_) are denoted by the black crosses. **B)** Representative steady-state binding responses for **CysE 10** (left) and **CymR 10** (right). The K_D_ was derived by steady-state binding analysis using the 1:1 binding model, with the vertical line representing the K_D_ value. The reported K_D_ values represent the mean ± standard deviation of at least two experiments.

Using X-ray crystallography, the interactions of each peptide with *Sa*CysK were then determined and compared to identify features that explain this discrepancy in binding affinity. Cocrystallization experiments resulted in the collection of two high-resolution datasets corresponding to the *Sa*CysK + **CysE 10** and *Sa*CysK + **CymR 10** complexes, solved at resolutions of 1.5 and 1.3 Å, respectively. Electron density maps for **CysE 10** and **CymR 10** are presented in Fig 5A. The same homodimeric structure of *Sa*CysK was observed in each structure, with no large structural perturbations observed relative to the holo form. As expected, a single binding event was observed within each monomer of *Sa*CysK, with the **CysE 10** and **CymR 10** peptides occupying the same active site region of *Sa*CysK (Fig 5B). In both cases electron density was only observed for a portion of the decapeptide, spanning positions D_7_ – I_10_ of **CysE 10** and G_5_ – I_10_ of **CymR 10**. The positioning and orientation of the peptide backbone is largely preserved for **CysE 10** and **CymR 10** which extends into the active site in a linear fashion. This is distinct from the previously reported CysK:CysE decapeptide complex of *H*. *influenzae* but is highly similar to the CysK:CysE tetrapeptide complex reported for *M. tuberculosis* (S1 Fig). The greatest difference in the binding mode between **CysE 10** and **CymR 10** is in the interactions involving the amino acid side chains, which differ substantially toward the N-terminal end of each peptide.

**Fig 5.**
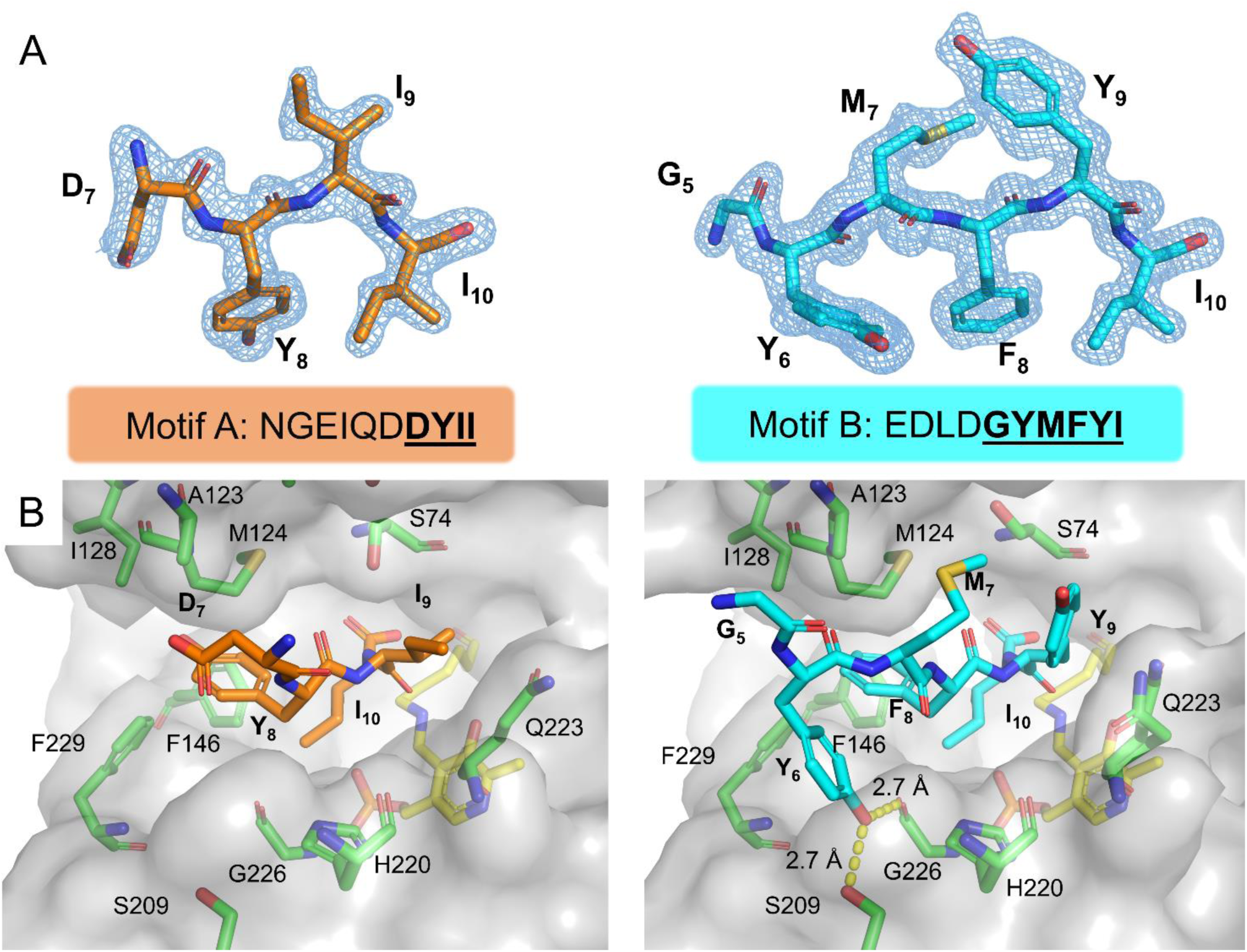
Overall binding mode of CysE 10 and CymR 10 within the active site of *Sa*CysK. **A)** Simulated annealing composite omit electron density maps (2 Fo-Fc, 1σ; blue mesh) for **CysE 10** (left; orange sticks) and **CymR 10** (right; cyan sticks). In each case, only a portion of the 10 amino acid peptide was ordered, with this region of the peptide indicated by the bold, underlined text. **B)** Binding of **CysE 10** (left; orange sticks) and **CymR 10** (right; cyan sticks) within the active site region of *Sa*CysK. The transparent surface of *Sa*CysK is shown in gray, with interacting sidechains shown as green sticks. The PLP adduct is shown as yellow sticks.

The shared C-terminal residue, I_10_, mediates extensive interactions with *Sa*CysK that are conserved for **CysE 10** and **CymR 10**. In total, four hydrogen bond interactions are formed by the C-terminal carboxylate group with the amido sidechain of Q145, the hydroxyl group of T73, and the backbone amides of N76 and T77 (Fig 6A). Similar interactions have been implicated in the recognition of the OAS substrate, supporting the role of **CysE 10** and **CymR 10** as direct competitors of OAS binding in *Sa*CysK [24]. The sidechain of I_10_ lies adjacent to the PLP adduct and packs tightly into a pocket lined by the sidechains of F146, T181, Y_8_/F_8_ of the respective peptide binder and the backbone of residues forming Loop 3 (G180 – T181) and Loop 4 (G224 – A227). Therefore, interactions involving I_10_ appear to play the same role in facilitating peptide recognition by *Sa*CysK but do not directly contribute to the large difference in the binding affinity of **CysE 10** and **CymR 10**.

**Fig 6.**
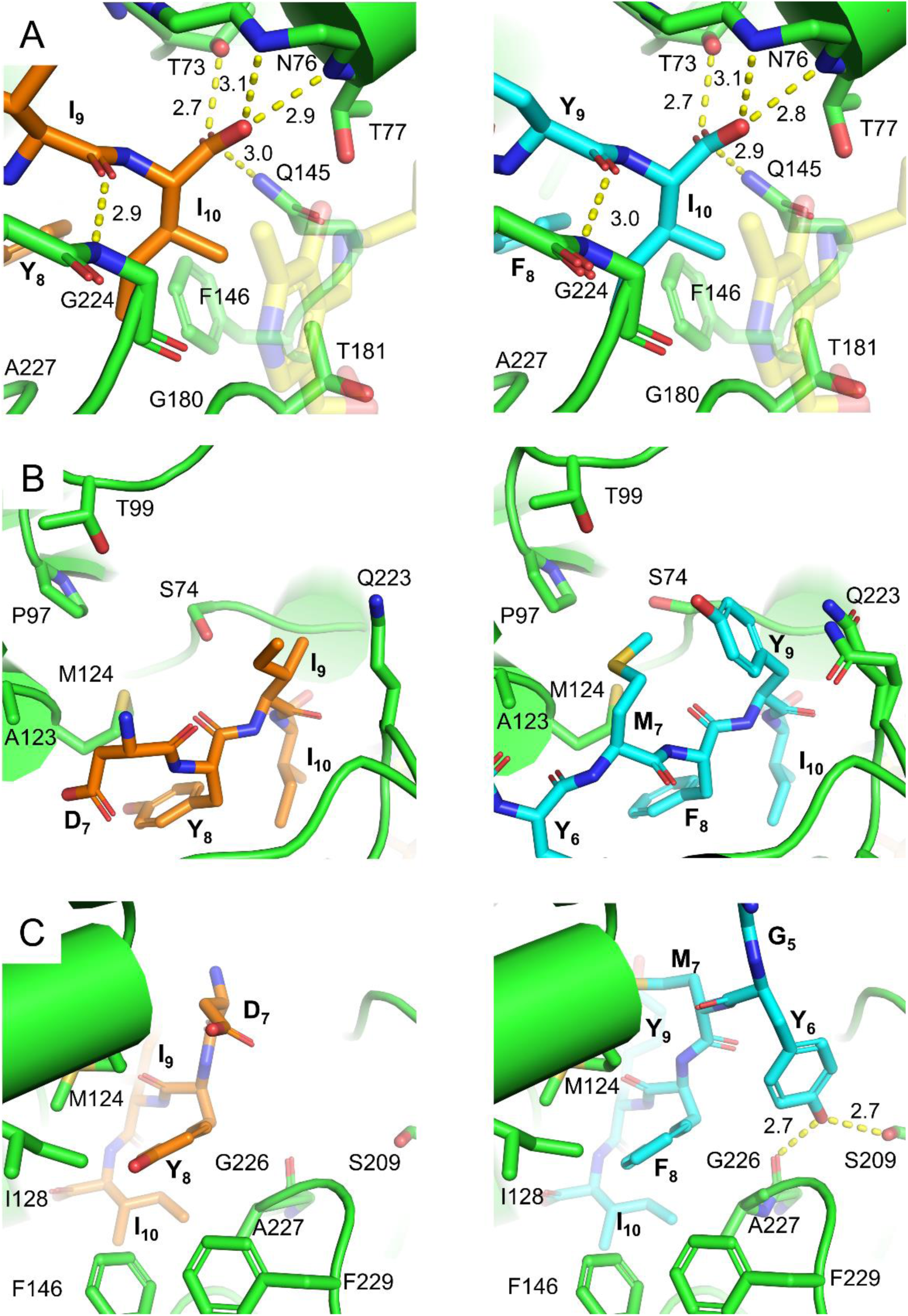
Structural comparison of CysE 10 and CymR 10 binding to *Sa*CysK. Comparison of binding at **A)** position 10, **B)** positions 7 and 9, and **C)** Positions 5, 6 and 8. **CysE 10** and **CymR 10** are shown as orange and cyan sticks, respectively. *Sa*CysK is represented by the green cartoon, with interacting residues shown as green sticks. The PLP adduct as shown as yellow, transparent sticks. Hydrogen bond interactions are indicated by yellow dashes, with distances reported in angstroms.

In both peptides position 9 is occupied by a hydrophobic residue, being I_9_ for **CysE 10** and Y_9_ for **CymR 10.** In each case the backbone carbonyl forms a hydrogen bond interaction with the backbone amide of Loop 4 residue G224 (Fig 6A). However, the sidechains of I_9_ and Y_9_ differ in their interactions. While both sidechains are positioned in a similar manner by interaction with Q223 of Loop 4, the bulky nature of Y_9_ in **CymR 10** results in the formation of a hydrophobic pocket together with S74, P97, T99, A123 and M124 of *Sa*CysK (Fig 6B). In **CymR 10** this pocket is occupied by M_7_, which behaves as a hydrophobic plug to further stabilize binding of **CymR 10**. In contrast, **CysE 10** possesses the acidic D_7_ at position 7 which projects in the opposite direction into space, participating in only weak van Der Waal interactions. The difference in residues at position 8 appears to play no part in this distinct difference, with Y_8_/F_8_ of **CysE 10** and **CymR 10** occupying a conserved hydrophobic channel lined by M124, I128, F146, A227, and F229 (Fig 6C).

In **CymR 10** electron density for the residues at position 5 and position 6 was also discernible, corresponding to residues G_5_ and Y_6_, suggesting these positions may also play an important role in the interaction with *Sa*CysK. Only van Der Waal interactions were evident between G_5_ and S121 – A123 of Helix 4 (Fig 6C). However, the hydroxyl group of Y_6_ forms two hydrogen bonds in a unique manner, simultaneously acting as a hydrogen bond donor with the backbone carbonyl of A227, and as a hydrogen bond acceptor with the hydroxyl sidechain of S209. This structural comparison reveals that the nanomolar affinity interaction observed for **CymR 10** can likely be attributed to a concerted effect involving the interplay between positions 7 and 9, and the additional hydrogen bond interactions formed at position 6, as neither of these interactions exist for **CysE 10**. In all, this identifies **CymR 10** as a promising alternative scaffold relative to CysE-derived peptides for the design of peptide inhibitors targeting *Sa*CysK.

### Structure-activity relationship (SAR) analysis of CymR 10 yields potent pentapeptide inhibitors of *Sa*CysK

Toward this endeavor, structure-activity relationship (SAR) analysis of the **CymR 10**:*Sa*CysK interaction was performed to define the requisite features of the **CymR 10** scaffold that confer high-affinity binding. We specifically aimed to define i) the minimal **CymR 10-**derived peptide that retains high-affinity binding, ii) the contributions of each residue within this peptide to the interaction with *Sa*CysK, and iii) whether this peptide is amenable to further development and optimization. In total, 16 rationally designed **CymR 10-**derived peptides were commercially sourced. These were initially characterized for binding to *Sa*CysK by SPR and then assessed for inhibitory activity against *Sa*CysK *in vitro* (Table 1). Finally, the interactions for the most potent of these designed peptides with *Sa*CysK were determined by X-ray crystallography.

**Table 1.**
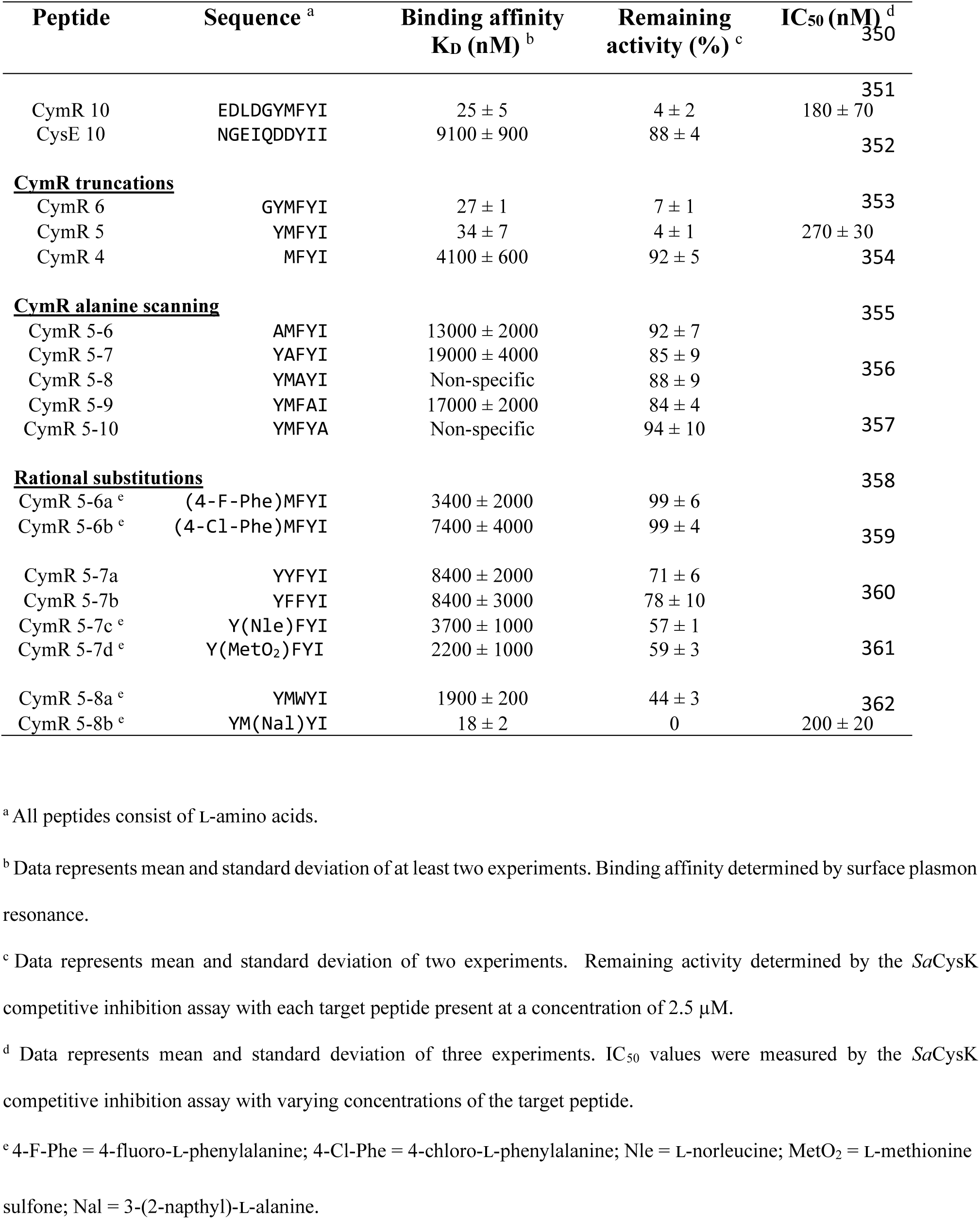
Binding affinities and inhibitory activities CymR 10-derived peptides with *Sa*CysK.

#### Binding affinities

First, a set of three **CymR 10** N-terminal truncated peptides were designed. Of these, **CymR 6** comprises the interacting region of **CymR 10** which displays strong electron density in the cocrystal structure (positions 5 – 10), while **CymR 5** and **CymR 4** correspond to positions 6 – 10 and 7 – 10, respectively. Analysis by SPR revealed that **CymR 6** and **CymR 5** bind with affinities of 27 ± 1 nM and 34 ± 7 nM, respectively, comparable to **CymR 10**. This indicates that G_5_ is dispensable for peptide binding and agrees with the cocrystal structure of **CymR 10** that showed very few interactions of G_5_ with *Sa*CysK. In stark contrast was **CymR 4,** which displayed a 160-fold lower affinity 4100 ± 600 nM, being comparable to the affinity of **CysE 10**. This apparent reduction in affinity is therefore attributed to the absence of Y_6_, which was observed to mediate two hydrogen bond interactions in **CymR 10**. Therefore, the pentapeptide **CymR 5** was identified as the minimal peptide that retains the high-affinity interaction with *Sa*CysK and so was used as the basis for designing subsequent peptides.

Having identified the minimal **CymR 5** scaffold, an alanine scan of **CymR 5** was performed to assess the individual contributions of amino acid sidechains. This yielded five peptides termed **CymR 5-6** to **CymR 5-10**, each containing a single alanine substitution at the labeled position (Table 1). In all cases binding was found to be greatly attenuated relative to **CymR 5**. **CymR 5-6**, **CymR 5-7**, and **CymR 5-9** still displayed specific binding to *Sa*CysK, possessing similar binding affinities in the range of 10 – 20 µM, being comparable to that of **CysE 10**. Interestingly, the lack of a graded decrease in affinity for these peptides indicates there is an interdependency between the formation of the hydrophobic plug by M_7_ and Y_9,_ and the formation of the hydrogen bond interactions mediated by Y_6_. In comparison, **CymR 5-8** and **CymR 5-10** showed no specific binding to *Sa*CysK at the highest assayed concentration of 100 µM. For **CymR 5-10** this was expected as I_10_ is strictly conserved in all interacting proteins of *Sa*CysK identified to date. The sidechain of Y_8_ also forms extensive interactions with the hydrophobic channel of *Sa*CysK. Therefore, it is unsurprising that loss of this sidechain in **CymR 5-8** greatly attenuates binding. This combined analysis indicates that all positions in the **CymR 5** scaffold contribute significantly to the interaction with *Sa*CysK.

Further interrogation of the **CymR 5** scaffold was also completed, with eight rationally designed peptides incorporating a single substitution of a proteinogenic or non-proteinogenic amino acid at position 6, 7 or 8 assayed by SPR. The aim of this experiment was to identify if these positions in the **CymR 5** scaffold are tolerant to structural modification, which may subsequently identify a starting point for the design of improved CymR-derived peptide inhibitors. Positions 9 and 10 were not interrogated as Y_9_ plays a key role in binding of M_7_, and the latter, I_10_, is known to be highly important for peptide recognition. Two peptides, **CymR 5-6a** and **CymR 5-6b**, contained substitutions of Y_6_ to 4-fluoro-phenylalanine and 4-chloro-phenylalanine, respectively. While both peptides displayed ∼2 to 3-fold higher binding affinity compared to the Y_6_ → A_6_ substitution in **CymR 5-6**, they were unable to replace the role of Y_6_ effectively despite the structural similarity. This reiterates the importance of the Y_6_ OH which behaves as a hydrogen bond donor and acceptor; the halogens F and Cl may only act as hydrogen bond acceptors and so cannot facilitate the same interaction. Substitution of M_7_ yielded similar results. **CymR 5-7a** and **CymR 5-7b** contained the bulkier hydrophobic residues tyrosine and phenylalanine in place of M_7_, respectively, and possessed the same binding affinity of 8.4 µM. Interestingly, substitution of M_7_ to the structurally similar norleucine in **CymR 5-7c** or methionine sulfone in **CymR 5-7d** was comparable, having respective binding affinities of 3.7 ± 1 µM and 2.2 ± 1 µM. This demonstrates that in the M_7_ – Y_9_ pair, M_7_ is optimal in acting as the hydrophobic plug, with sidechains of larger or lesser volume unable to do so effectively. In contrast to positions 6 and 7, position 8 was found to tolerate a substitution and retain high-affinity binding. While substitution of F_8_ to the bulkier tryptophan in **CymR 5-8a** resulted in an ∼60-fold lower binding affinity compared to **CymR 5**, substitution of F_8_ to the structurally similar 3-(2-napthyl)-ʟ-alanine (Nal) in **CymR 5-8b** produced a binding affinity of 18 ± 2 nM. This represents a ∼2-fold improvement over the **CymR 5** scaffold and indicates that in contrast to the findings for positions 6 and 7, position 8 of **CymR 5** has a level of tolerance to structural modification and may serve as a starting point for further optimization of the scaffold.

#### Inhibitory activities

Next, the ability of the 16 rationally designed peptides to inhibit the enzymatic activity of *Sa*CysK was assessed *in vitro*. First, each peptide was assayed at a single intermediate concentration of 2.5 µM with the outcome of this initial screen consistent with the results of the SPR analysis (Table 1). This confirmed that CymR-derived peptides can inhibit *Sa*CysK activity, with the high-affinity binders **CymR 10**, **CymR 6**, **CymR 5**, and **CymR 5-8b** all causing a > 90% reduction in the OASS activity of *Sa*CysK. **CymR 5-7c**, **CymR 5-7d**, and **CymR 5-8a** also produced ∼ 50 % inhibition of *Sa*CysK, consistent with the low micromolar binding affinities of these peptides, while the remaining peptides produced little to no inhibitory effect. For the most potent of these peptides, **CymR 10**, **CymR 5** and **CymR 5-8b** IC_50_ values were also determined (Table 1 and S2 Fig). This revealed that truncation of **CymR 10** has a minimal effect on inhibitory activity, with **CymR 10** and **CymR 5** possessing IC_50_ values of 180 ± 70 and 270 ± 30 nM, respectively. The modified peptide **CymR 5-8b** displayed comparable potency to **CymR 10** with an IC_50_ value of 200 ± 20 nM. This confirms that CymR-derived peptides can inhibit *Sa*CysK activity and that the **CymR 5** pentapeptide scaffold retains potent inhibitory activity.

Finally, the growth inhibitory ability of the **CymR 5-8b** was assessed versus cultured *S. aureus*. This peptide exhibited no growth inhibition of the USA300 MRSA *S. aureus* strain FPR3757 grown in rich (LB) media at concentrations as high as 200 µM (S3 Fig). This lends well to the prospect of **CymR 5-**derived peptides as antibacterial adjuvants, since low antibacterial activity of an adjuvant is preferable. To assess whether **CymR 5-8b** was able to inhibit *Sa*CysK *in bacterio*, the growth inhibitory activity of the peptide was also assessed versus the USA300 *S. aureus* strain LAC grown in defined media supplemented with either cysteine or thiosulfate as the sulfur source (S3 Fig). As CysK activity is required for growth of *S. aureus* when utilizing thiosulfate as a sulfur source [37], we hypothesized that it would be sensitized to **CymR 5 8-b** if it was effectively inhibiting *Sa*CysK. Although some minor growth inhibition was noted at the higher concentrations tested, this was comparable for both sulfur sources. The lack of thiosulfate-dependent growth inhibition observed under these conditions suggested that **CymR 5-8b** was either unable to enter the cell or was rapidly degraded upon entry. These possibilities were explored by measuring the accumulation of **CymR 5-8b** within the *in vitro* cultured strain LAC grown in defined media using an intrabacterial drug accumulation/metabolism (IBDM) platform with a lower detection limit of ∼ 1 nM [43]. Neither **CymR 5-8b** nor a limited set of metabolites were detected suggesting that it does not accumulate intracellularly and explains the lack of growth inhibition noted (S1 Supporting Information). Therefore, the cellular uptake and metabolic stability of CymR-derived peptides must be improved prior to assessing the adjuvant activity of these *Sa*CysK inhibitors.

#### Structural characterization

Finally, the cocrystal structures of *Sa*CysK complexed with the most potent pentapeptides **CymR 5** and **CymR 5-8** were determined by X-ray crystallography to allow comparison with the binding mode identified for **CymR 10**. The overall structure of *Sa*CysK was conserved in each case, and the binding modes of **CymR 5** and **CymR 5-8b** closely resembled that of **CymR 10** (Figs 7A–7C. However, electron density for the Y_6_ sidechain was surprisingly absent in both pentapeptides. Close inspection of the crystal lattice surrounding the active site region identified alternative crystal contacts in both structures that directly clash with the binding mode of Y_6_ seen for **CymR 10** (S4 Fig). Combined with the SPR analysis that verified the importance of Y_6_ in the pentapeptide scaffold, this strongly suggests that the absence of Y_6_ is a crystallographic artifact and that Y_6_ has the same role in both pentapeptides. For **CymR 5-8b** the bulkier Nal sidechain retains the interactions of Y_8_ but fills the entirety of the hydrophobic channel, forming tighter hydrophobic interactions with the sidechains of M124, I128 and F146 (Fig 7C). Beyond the hydrophobic channel also exists an opening to an additional small cavity lined by the sidechains of P72, E71, E142, Q144 and E147, suggesting that modifications at position 8 that extend further into this space would be feasible (Fig 7D). Together, this structural comparison indicates that the shorter pentapeptides **CymR 5** and **CymR 5-8b** retain the same binding mode and interactions as **CymR 10** and identifies position 8 as a promising starting point for the development of more potent *Sa*CysK inhibitors based on the **CymR 5** scaffold.

**Fig 7.**
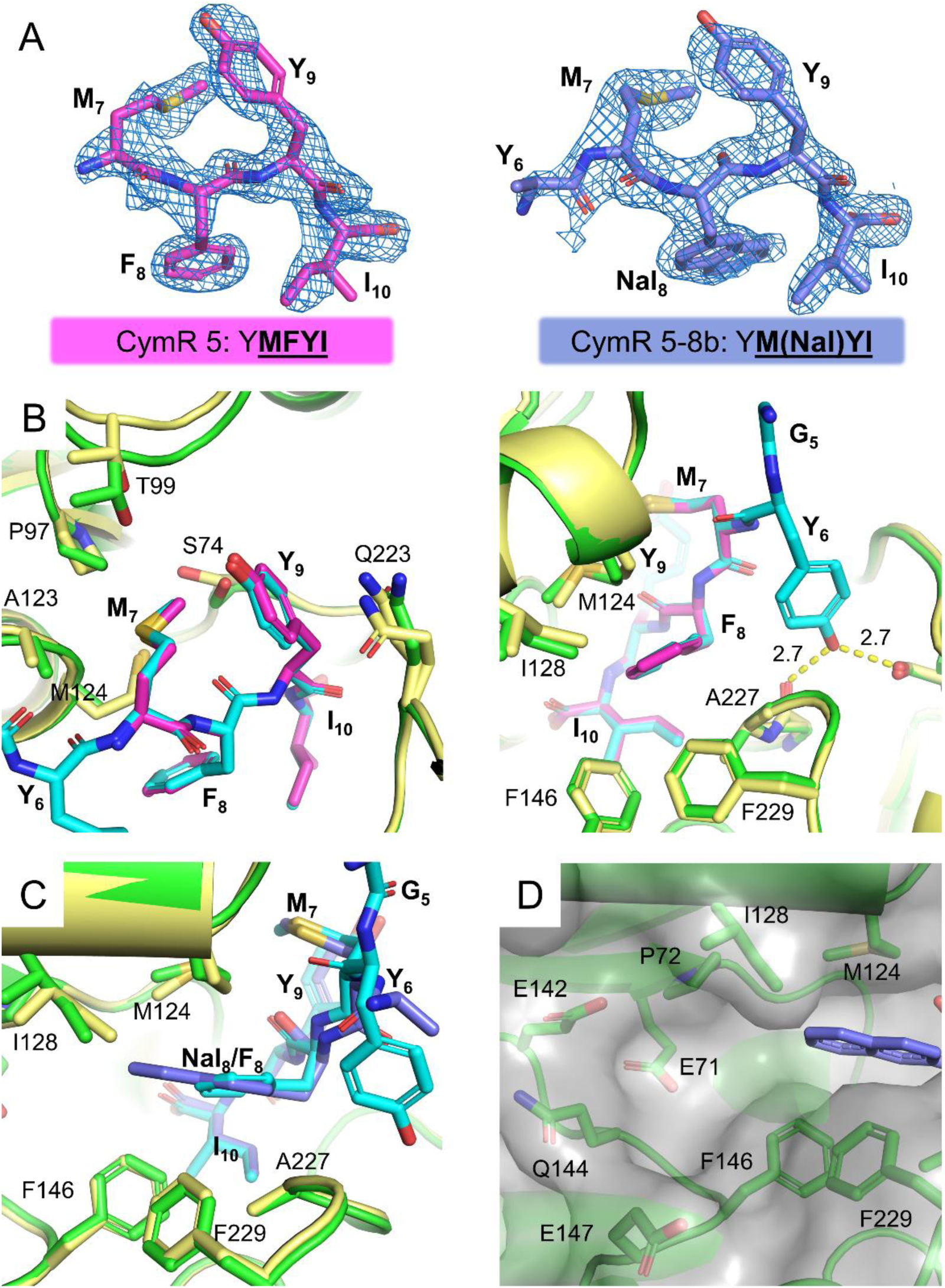
Structural characterization of CymR 5 and CymR 5-8b binding to *Sa*CysK. **A)** Simulated annealing composite omit electron density maps (2Fo-Fc, 1σ; blue mesh) for **CymR 5** (left; pink sticks) and **CymR 5-8b** (right; purple sticks). **B)** Structural comparison of **CymR 5** (pink sticks) and **CymR 10** (cyan sticks) binding to the active site region of *Sa*CysK. **C)** Structural comparison of **CymR 5-8b** (purple sticks) and **CymR 10** (cyan sticks) binding at position 8. **D)** Location of the small cavity lying beyond the hydrophobic channel occupied by the sidechain of Nal_8_. The *Sa*CysK:**CymR 5** and *Sa*CysK:**CymR 5-8b** complexes are represented by green cartoon, with the *Sa*CysK:**CymR 10** complex shown as yellow cartoon. The hydrogen bond interactions formed by Y_6_ of **CymR 10** are indicated by yellow dashes, with distances labelled in angstroms.

### The CymR 5 scaffold is a promising lead for the development of peptide inhibitors targeting CysK

The functional and structural characterisation of the interactions between *Sa*CysK and CysE or CymR-derived peptides provides new insight toward the development of peptide based CysK inhibitors. Prior to this work only CysE-derived peptides had been investigated in this capacity. These previous studies concluded that within inhibitory pentapeptides the three C-terminal amino acids determine the affinity of the interaction with CysK [31,32]. Here we interrogated the structural basis of the CysK:CymR interaction for the first time. This identified that the **CymR 5** pentapeptide participates in a novel interaction with *Sa*CysK where all five positions work in concert to impart a > 10-fold higher affinity interaction than the most potent CysE-derived peptide reported. While already a significant advancement in potency relative to CysE-derived peptide inhibitors, we expect that the **CymR 5** scaffold can be optimized to yield further improvements. **CymR 5** therefore provides an alternative avenue toward the optimisation and development of peptide inhibitors targeting *Sa*CysK. Furthermore, as the structure and amino acid composition of the active site region is highly conserved between bacterial CysK enzymes, **CymR 5** may also prove suitable for developing inhibitors that target other CysK enzymes of interest.

Although CymR-derived peptides are effective inhibitors of isolated *Sa*CysK, the lack of cell accumulation observed for **CymR 5-8b** represents the largest hurdle to achieving effective inhibition of *Sa*CysK *in bacterio*. Therefore, we believe that chemical modification of **CymR 5** should be prioritized to improve this property. From the SAR analysis performed in this study it was determined that chemical modification of **CymR 5** is tolerated in a limited capacity and most frequently leads to a substantially weaker interaction with *Sa*CysK. Therefore, to guide this process we have identified several positions that appear amenable to chemical modification, as detailed in Fig 8. First are positions 7 and 9 which form the hydrophobic plug. While substitutions at position 7 resulted in decreased affinity the effect of substitutions at position 9, particularly how this effects formation of the hydrophobic plug, was not explored. Investigation of simultaneous substitution at positions 7 and 9 using a wider range of natural and non-proteinogenic amino acids may uncover alternative combinations that facilitate formation of the hydrophobic plug. If successful, this would allow for greater chemical diversity within the **CymR 5** scaffold while retaining or improving potency. Second is position 8 which places the F_8_ sidechain within the hydrophobic pocket. Replacing F_8_ with a reactive handle would allow extension through the hydrophobic cleft and into the cavity highlighted in Fig 7D, with possible benefits including increased potency, specificity and increased stability through shielding of the peptide backbone from proteolysis. Last is the N-terminus of **CymR 5**. The observation the **CymR 10** and **CymR 5** bind with similar affinity indicates that chemical modifications at the N-terminal primary amide can likely be tolerated. This provides the most direct route to improving the cellular uptake of CymR-derived peptides through altering the chemical properties of the scaffold or through conjugation of cell penetrating peptides, which may also contribute to increased metabolic stability. If successful, this approach will allow for the development of improved peptide inhibitors of *Sa*CysK and allow evaluation of these inhibitors as antibiotic adjuvants.

**Fig 8.**
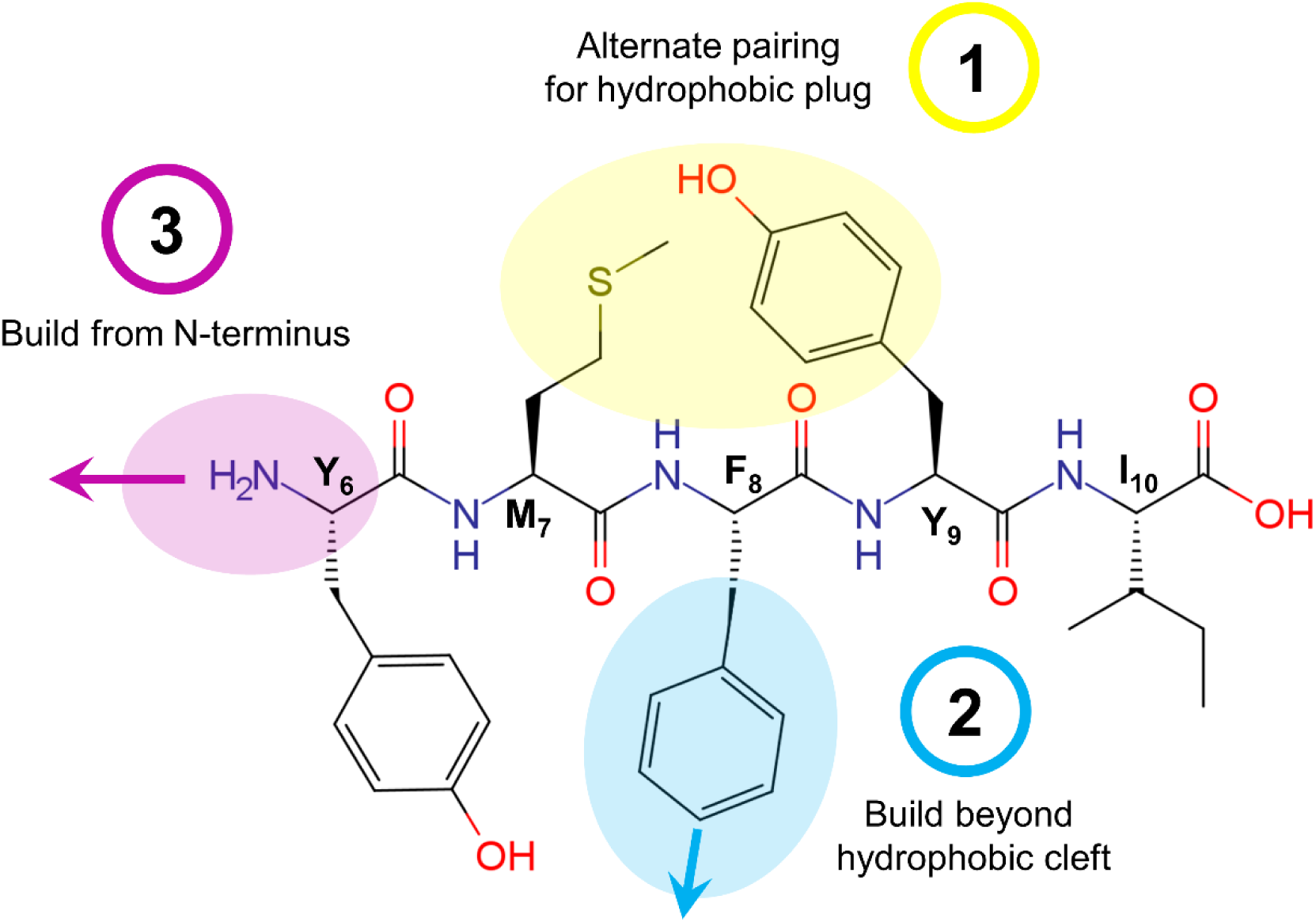
Proposed avenues for the optimisation of the CymR 5 scaffold. Chemical modifications at position 6, 8 and dual modifications at positions 7 and 9 represent feasible routes to maintain potency and improve cell permeability and metabolic stability.

## Materials and Methods

### Materials and reagent preparation

Oligonucleotides for cloning of *cysK* were purchased from Sigma. All peptides included in this study were purchased from Genscript with the purities listed in S1 Table. HPLC and MS traces for the key target peptides **CysE 10**, **CymR 10**, **CymR 5** and **CymR 5-8b** are provided in (S5 Fig). For all experiments peptide stock solutions (10 mM) were prepared in MilliQ water and neutralized to pH 7.5 by dropwise addition of 2M sodium hydroxide. OAS hydrochloride (A6262), sodium sulfide nonahydrate (208043) and PLP hydrate (P9255) were purchased from Sigma. Due to the unstable nature of sodium sulfide and OAS in aqueous solution, stock solutions of these substrates were prepared immediately prior to use. Sodium sulfide was prepared as a 20 mM solution in MilliQ water. OAS was prepared as a 200 mM stock solution in 300 mM sodium hydroxide, giving a final pH of ∼8.0. PLP was prepared as a 10 mM stock solution in MQ water and stored as single use aliquots at -20 °C. The ninhydrin reagent was prepared fresh each day at room temperature and contained 250 mg ninhydrin, 6 ml of glacial acetic acid and 4 ml of 32% hydrochloric acid.

### Cloning of S. aureus cysK

The gene encoding *Sa*CysK (saouhsc_00488; encodes equivalent protein to SA0471 in *S. aureus* N315) was amplified from *S. aureus* NCTC 8325 genomic DNA by PCR, with oligonucleotides incorporating an N-terminal hexahistidine tag and restriction endonuclease recognition sites, as detailed below. Digested fragments were ligated into pET-16b using *NcoI* and *BamHI* restriction endonuclease sites and transformed into *E. coli* BL21(DE3) for recombinant protein expression. All resulting DNA products were confirmed by Sanger sequencing.

Forward: 5’ AAA**CCATGG**GC*CATCACCATCACCATCAC*ATGAAAATTTCTACTAAAGGGAGAT 3’

The restriction endonuclease sites are shown in bold underlined text, and the hexahistidine tag indicated by underlined italicised text.

Reverse: 5’ AAA**GGATCC**TTAAATATAAAACATGTATCCGTCTAAAT 3’

### Expression and purification of *Sa*CysK

Two flasks containing 1 L LB + 100 μg/ml ampicillin were inoculated with 10 ml of a saturated overnight culture and grown at 37 °C. Upon reaching an A_600_ of 0.5 PLP was added to a final concentration of 100 µM and expression was induced by the addition of 1 mM IPTG at 16 °C overnight. Cell pellets were collected by centrifugation at 5000*g* (5 min, 4 °C), resuspended in 25 ml of Buffer A (50 mM Tris–HCl pH 8.0, 300 mM NaCl, 10 mM imidazole and 5 mM β-mercaptoethanol) and lysed by mechanical disruption. The lysate was clarified by centrifugation at 40,000*g* (10 min, 4 °C). Clarified lysate was applied to a 5 ml HisTrap HP column (GE Life Sciences) connected to an NGC Medium-Pressure Liquid Chromatography System (Bio-Rad) equilibrated in Buffer B (50 mM Tris–HCl pH 8.0, 300 mM NaCl, 10 mM imidazole, 5 mM β-mercaptoethanol), followed by washing with five column volumes of 10% Buffer C (50 mM Tris–HCl pH 8.0, 300 mM NaCl, 250 mM imidazole, 5 mM β-mercaptoethanol). *Sa*CysK was eluted in a single step with five column volumes of 100% Buffer B. Eluted fractions containing *Sa*CysK were combined (10 ml total) and applied to a HiPrep 26/60 Sephacryl S-200 HR gel filtration column (GE Life Sciences) equilibrated with Buffer D (20 mM Tris–HCl pH 8.0, 50 mM NaCl, and 1 mM DTT). Fractions containing *Sa*CysK were pooled and concentrated using an Amicon Ultra centrifugal filter (10,000 molecular weight cutoff, Sigma-Aldrich) to 25 mg/ml and stored at −80 °C. For enzyme activity measurements, *Sa*CysK was diluted to 1 mg/ml in Buffer D and stored at −80 °C in single use aliquots. *Sa*CysK concentration was determined by measuring absorbance of the PLP cofactor using the experimentally determined molar extinction coefficient of ε_412nm_= 9090 M^-1^·cm^-1^, as determined by the alkali denaturation method [44].

### Crystallization, data collection and refinement

Crystallization experiments to obtain structures of holo *Sa*CysK and the *Sa*CysK-**CysE 10** and *Sa*CysK-**CymR 10** complexes were performed using the sitting drop vapor diffusion method in 96-well Intelliplates (Art Robbins). The *Sa*CysK-CysE10 and *Sa*CysK-CymR-10 complexes were formed by combining 117 µl 25 mg/ml *Sa*CysK and 13 µl of 1 mM CysE10 or CymR10, respectively, and incubated on ice for 30 min. Conditions promoting crystal formation were probed using the PEG/Ion HT, Index HT, Crystal Screen HT (Hampton Research), and NR-LBD HT (Molecular Dimensions) sparse matrix screens at 16 °C. Reservoirs contained 80 μl of each condition with drops formed by adding 1 μl of 25 mg/ml *Sa*CysK or the *Sa*CysK-peptide complex to 1 μl of reservoir solution. Crystals of holo *Sa*CysK and the *Sa*CysK-**CysE 10** complex were obtained in Index HT condition B8 (200 mM ammonium acetate, 100 mM HEPES-NaOH pH 7.5 and 25 % PEG 3350). Crystals of the *Sa*CysK-**CymR 10** complex were obtained in NR-LBD HT condition E5 (100 mM NaCl, 100 mM PIPES pH 7.0 and 22 % PEG 2000 MME). In all cases crystals formed as rectangular prisms, appearing within a week.

Crystallization experiments to obtain structures of the *Sa*CysK-**CymR 5** and *Sa*CysK-**CymR 5**-**8b** complexes were performed using a microseeding approach by the hanging drop vapor diffusion method in 24 well plates containing 200 mM ammonium acetate, 100 mM HEPES-NaOH pH 7.5 and 20 – 25 % PEG 3350 as the reservoir solution. The complexes were formed by combining 27 µl of 25 mg/ml *Sa*CysK and 3 µl of 10 mM **CymR 5** or **CymR 5-8b** peptide followed by incubation on ice for 30 min. Wells contained 500 µl of reservoir solution, with drops formed by adding 1 µl of the *Sa*CysK-peptide complex and 0.4 µl of a 1:100 *Sa*CysK seed stock to 1 µl of reservoir solution. Plates were stored at 22 °C, with crystals consistently forming as thin plates within 3 days. In the absence of seed stock crystal formation was not observed. To prepare the neat *Sa*CysK seed stock a single crystal of holo *Sa*CysK (obtained by the sitting drop method in Index HT condition B8) was finely crushed with a steel needle. The crystal fragments were collected by pipetting and transferred to 50 µl of reservoir solution (200 mM ammonium acetate, 100 mM HEPES-NaOH pH 7.5 and 25 % PEG 3350), followed by vortexing at maximum speed for 10 min. The neat seed stock was diluted to 100-fold in reservoir solution and stored at 4 °C for up to one month. New seed stocks were prepared from crystals of holo *Sa*CysK grown using the same microseeding approach detailed above.

For data collection crystals were transferred to NVH immersion oil for cryoprotection and flash-frozen in liquid nitrogen. Datasets were collected at the MX1 and MX2 beamlines of the Australian Synchrotron [45,46]. Indexing, integration and scaling were completed using XDS, with Aimless (CCP4) used for merging of scaled datasets [47,48]. The holo structure of *Sa*CysK (PDB: 8SRT) was solved by molecular replacement using Phaser, with the holo structure of *Mycobacterium tuberculosis* CysK (PDB: 2Q3B) used as the search model [49]. The phase problem for peptide-bound structures of *Sa*CysK were subsequently solved using Phaser and the holo *Sa*CysK as the search model. The resulting models were subjected to multiple rounds of rebuilding in Coot and refinement in Phenix until the refinements converged [50,51]. Data collection, processing and refinement statistics are presented in S2 Table.

### Surface plasmon resonance binding assays

SPR binding assays were performed using a Biacore S200 instrument (GE Healthcare) using a NIHC 1500 M sensor chip (Xantec Bioanalytics) at 25°C. For these experiments *Sa*CysK was employed as the receptor, being immobilized via capture of the N-terminal hexahistidine tag by the Ni^2+^-NTA surface of the sensor chip. The sensor chip surface was first regenerated by a 60 s injection of 500 mM imidazole pH 8.0, a 60 s injection of Running Buffer and a final 60 s injection of 500 µM NiCl_2_, with all steps performed at a flow rate of 10 µl/min. The immobilization step was performed using Running Buffer (10 mM HEPES pH 7.5, 150 mM NaCl, 0.05% Tween 20 and 12 µM PLP). First, a 10 µl aliquot of 1 mg/ml *Sa*CysK in Buffer D was diluted in Running Buffer to a final concentration of 500 nM. This sample was injected across the target flow cell (flow cell 2) at a flow rate of 10 µl/min until ∼ 3000 – 4500 RU of *Sa*CysK was captured. The surface of flow cell 1 was kept blank for reference subtraction. Prior to binding experiments, the sensor chip was allowed to equilibrate in Running Buffer for ∼30 min to achieve a stable baseline. On completion of an experiment the regeneration procedure was repeated to clean the chip surface for reuse in subsequent binding experiments.

Binding experiments were performed in Running Buffer, with each peptide analyte assayed as 2-fold serial dilutions consisting of 8 concentrations, with a buffer only injection included between each serial dilution. Binding curves were obtained by injecting varying concentrations of peptide across the sensor chip at a flow rate of 30 µl/min for 30 s followed by a 60 s dissociation phase. All sensorgrams were double referenced. K_D_ values were determined by steady-state analysis using the 1:1 binding model in Biacore S200 Evaluation Software (GE Healthcare). The mean K_D_ and standard deviation were determined from at least two experiments. Representative sensorgrams and steady-state binding responses for all peptides analysed in this study are presented in S6 Fig.

### Measurement of *Sa*CysK activity

The activity of *Sa*CysK was measured through monitoring the formation of ʟ-cys using a discontinuous colorimetric assay based on the method of Gaitonde [38]. The preparation of the OAS, sodium sulfide and ninhydrin reagent stock solutions mentioned herein is outlined in the “Materials and reagent preparation” section. All experiments were performed at 37 °C in 1.5 ml polypropylene tubes. Prior to starting the reaction *Sa*CysK and the reaction mixture were equilibrated at 37 °C for 5 min. Reactions were initiated by adding 50 µl of 2×*Sa*CysK to 50 µl of 2×reaction buffer, giving a final reaction volume of 100 µl. The reactions were allowed to proceed for exactly 5 min, with the activity of *Sa*CysK quenched by addition of 100 µl glacial acetic acid.

For color development, 100 µl of ninhydrin reagent was added to each sample and the sample mixed by vortexing. Each sample was then placed into a metal block equilibrated at 110 °C and heated for 4 min, with a second metal block equilibrated at 110 °C held firmly on the samples to prevent evaporation and limit condensation. A change in color from pale yellow to pink is observed at this step, with the intensity of the pink color proportional to the cysteine content of the sample. Following color development, each sample tube was immediately transferred to a third metal block equilibrated at 22 °C, then vortexed and centrifuged to ensure complete mixing of the sample. To stabilise the color 100 µl of each sample was then transferred to 300 µl of ice-cold absolute ethanol and mixed by pipetting. The resulting 400 µl samples were equilibrated at 22 °C for 5 min and then 300 µl of each sample transferred to a single well of a 96-well flat bottom plate (Corning 3370). Absorbance was measured at 560 nm (Pherastar FX microplate reader; BMG Labtech), with the absorbance of a no enzyme sample used for blank subtraction. The concentration of ʟ-cys was determined using the slope of a ʟ-cys standard curve generated by the above method, with good linearity observed for samples containing an initial concentration of ʟ-cys between 5 and 160 µM (S7 Fig).

For the determination of kinetic parameters (*K*_M app_ OAS and *k*_cat_) reactions contained 50 mM Tris-HCl pH 7.5, 5 µM PLP, 1 mM sodium sulfide, 1 nM CysK and varying concentrations of OAS ranging between 1.25 and 80 mM. Data were fit with the Michaelis–Menten Equation (Equation 1) by nonlinear regression in GraphPad Prism 9. The experiment was performed three times, with a single replicate measured for each concentration assayed per experiment.

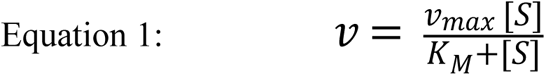

### *Sa*CysK competitive inhibition assay

To assess the inhibitory activity of CysE/CymR-derived peptides against *Sa*CysK the same colorimetric assay was used. For initial screening of *Sa*CysK inhibition by the CysE/CymR-derived peptides, reactions contained 50 mM Tris-HCl pH 7.5, 5 µM PLP, 1 mM sodium sulfide, 10 mM OAS, 4 nM *Sa*CysK and 2.5 µM peptide. The remaining *Sa*CysK activity was determined relative to a sample containing no CysE/CymR-derived peptide. The experiment was performed twice, with a single replicate measured for each concentration assayed per experiment. For determination of IC_50_ values reactions similarly contained 50 mM Tris-HCl pH 7.5, 5 µM PLP, 1 mM sodium sulfide, 10 mM OAS and 4 nM *Sa*CysK, but with the concentration of the CymR-derived peptide varied. Peptides were assayed at 8 different concentrations, and the remaining *Sa*CysK activity was determined relative to a sample containing no CysE/CymR-derived peptide. IC_50_ values were calculated using a log(inhibitor) vs response – variable slope model in GraphPad Prism 9 (GraphPad Software, Inc.). The experiment was performed in triplicate, with a single replicate measured for each concentration assayed per experiment.

### Determination of minimum inhibitory concentrations (MICs) and growth inhibition against *S. aureus*

Initially, a broth microdilution assay was performed to determine the minimum inhibitory concentration (MIC) of a compound tested against the methicillin-resistant USA300 *S. aureus* strain FPR3757 (ATCC BAA-1556). An overnight culture of bacteria was subcultured 1:100 in fresh LB media and grown at 37 °C and 180 rpm until A_600_ of 0.2 – 0.3. The bacterial culture was then diluted 1:1000 in LB media and added to a 96-well microtiter plate (100 µl per well) containing different concentrations of serially diluted compound. A DMSO solution of **CymR 5-8b** was added in duplicate and serially diluted across the plate (starting concentration of 200 µM). DMSO-treated wells and JSF-3151 (starting concentration 100 µM) were used as the negative and positive controls, respectively [43]. The plate was grown at 37 °C without shaking overnight. After 18 h, MIC was determined as the lowest concentration of drug, resulting in complete growth inhibition by visual inspection.

To determine if **CymR 5-8b** inhibited *S. aureus* growth under sulfur-limited conditions we used a previously described chemically defined medium that did not contain a sulfur source which can be utilised by *S. aureus* [52]. This medium consists of deionized water containing 3.2 mM KH₂PO₄, 10.8 mM Na₂HPO₄, 22.3 mM NaCl, 48.6 mM NH₄Cl, 28.9 mM glucose and 6 mM MgSO₄. Also included is 3.5 mM of aspartate, leucine, alanine, glycine, and valine, 2.6 mM proline, 2 mM arginine, 1 mM of histidine, tyrosine, isoleucine, phenylalanine, serine, and threonine, 0.35mM glutamate and 0.02mM of lysine and tryptophan. Where indicated, 50µM of either thiosulfate or cysteine were also added as a sulfur source. One mL of cells from an overnight culture of the USA300 MRSA strain LAC grown in tryptic soy broth was pelleted and washed with phosphate-buffered saline (PBS) before suspending in 1 mL of PBS [53]. These cells diluted to an optical density (A_600_) of 0.01 in 200 µL of defined media containing 0-1000 nM CymR 5-8b in a 96-well assay plate. Cells were grown with shaking at 200 rpm at 37°C for 18 hours. After which time, the optical densities of the cultures were quantified.

### Intrabacterial drug accumulation/metabolism (IBDM) assay

The IBDM assay was performed with the methicillin-resistant USA300_*S. aureus* MRSA strain LAC. All solutions added to cell pellets were pre-chilled over ice and all centrifugations were performed at 4 °C to minimize further metabolic processes post-sample collection. An overnight culture of bacteria was subcultured 1:100 in fresh chemically defined media and grown at 37 °C and 180 rpm. A concentration of 50 µM compound was added to five 1 mL bacterial cultures at optical density A_600_ of 0.3 – 0.4 (maximum A_600_ in this medium is ∼0.8). A DMSO treated culture was used as the negative control. After 60 min of incubation, samples were centrifuged at 10,000 rpm for 5 min. The cells were then washed with ice cold 0.85% NaCl solution. The resulting cell pellet was quenched with an ice-cold mixture of 2:2:1 CH_3_OH:CH_3_CN:H_2_O (MAW). The resulting solution was then lysed by four cycles of freeze-thaw (5 min freeze/30 s thaw) in a dry ice/acetone bath. The cell lysate was subsequently centrifuged at 13,000 rpm for 5 min. 700 µL of supernatant was then transferred to a 0.22-micron filter tube and centrifuged at 13,000 rpm for 10 min to filter any remaining cellular debris. Samples were stored in a -80 °C freezer until LC/MS analysis.

### LC/MS analysis of biological lysates

Accumulation was measured using an Agilent 1290 Infinity II liquid chromatography system coupled with an Agilent 6125 single quadrupole mass spectrometer. Liquid chromatography separation was achieved using an Agilent Poroshell 120 C-18 column with 1.9 µm particle size, 2.1 mm internal diameter, and 50 mm length. The solvent system consisted of CH_3_CN and H_2_O supplemented with 0.1% formic acid. The gradient involved four main phases: 1) an isocratic phase from 0 – 0.3 min at 5% CH_3_CN, followed by a 2) 0.3 – 0.6 min increasing CH_3_CN to 30% then 3) 0.6 – 3.3 min increasing CH_3_CN from 30% to 95%, and 4) an isocratic phase from 3.3 – 3.6 min at 95% CH_3_CN. This was followed by a post-run for 1 min at 95% H_2_O. The flow rate was set to 0.5 mL/min, and column temperature was set to 40 °C. For mass spectrometry, all analyses were performed using an API-ES ion source in positive polarity. The following spray chamber parameters were used: nebulizer pressure of 40 psi, capillary voltage of 3000 V, drying gas temperature of 350 °C, and drying gas flow of 12.0 L/min. Two main methods were used to quantify intrabacterial accumulation. Firstly, a scan method ranging from 100 to 1000 m/z was used to investigate the retention time of the compound being tested and its ionization. Then a selective ion monitoring (SIM) method was employed to maximize sensitivity. SIM method was set up to search for two target masses: the peptide tested and the internal standard (verapamil).

### LC-TOF analysis for metabolites

To search for metabolites of a compound, an Agilent 1260 Infinity II liquid chromatography coupled with an Agilent 6230 time-of-flight mass spectrometry (TOF) was used to analyze the IBDM samples. LC and spray chamber parameters were identical to LC-MS conditions listed above with the exception of a longer run time (10 min) to accommodate for the lower flow rate (0.2 mL/min) used in the 1260 Infinity II LC system. The Agilent Technologies MassHunter Quant software version 10.0 combined with Agilent Personal Compound Database and Library (PCDL version 8.0) were used to search for a list of peptide fragments in the compound treated sample and the DMSO treated control.

### Data analysis and visualisation

Structural superposition and generation of figures for visualisation of enzyme structure was performed in PyMOL [54]. Multiple sequence alignments were performed using Clustal Ω [55]. Marvin was used for drawing, displaying chemical structures, Marvin 22.22, Chemaxon (https://www.chemaxon.com).

## Acknowledgements

BCV was supported by an Australian Government Research Training Program (RTP) scholarship. This research was undertaken in part using the MX1 and MX2 beamlines at the Australian Synchrotron, part of ANSTO, and made use of the Australian Cancer Research Foundation (ACRF) detector. JSF acknowledges funding from NIH grant U19AI109713. The Boyd lab is supported by NIAID 1R01AI139100-01, NSF 1750624, and USDA MRF project NE−2248.

## Author contributions

JLP, BCV, AG and HB; data curation, formal investigation, visualization and methodology JLP; conceptualisation and writing original draft. JLP, BCV, AG, HB, JMB, JSF and JBB; writing, review and editing. JMB, JSF and JBB; supervision and resources.

## Data availability statement

The structural data that support these findings were deposited in the wwPDB with PDB IDs 8SRT, 8SRU, 8SRV, 8SRW and 8T2C. Remaining data that support the findings of this study are available from the corresponding author upon reasonable request.

