## Supplementary figures and tables for "Identification of cysteine metabolism regulator (CymR)-derived pentapeptides as nanomolar inhibitors of *Staphylococcus aureus* O-acetyl-ʟ-serine sulfhydrylase (CysK)"

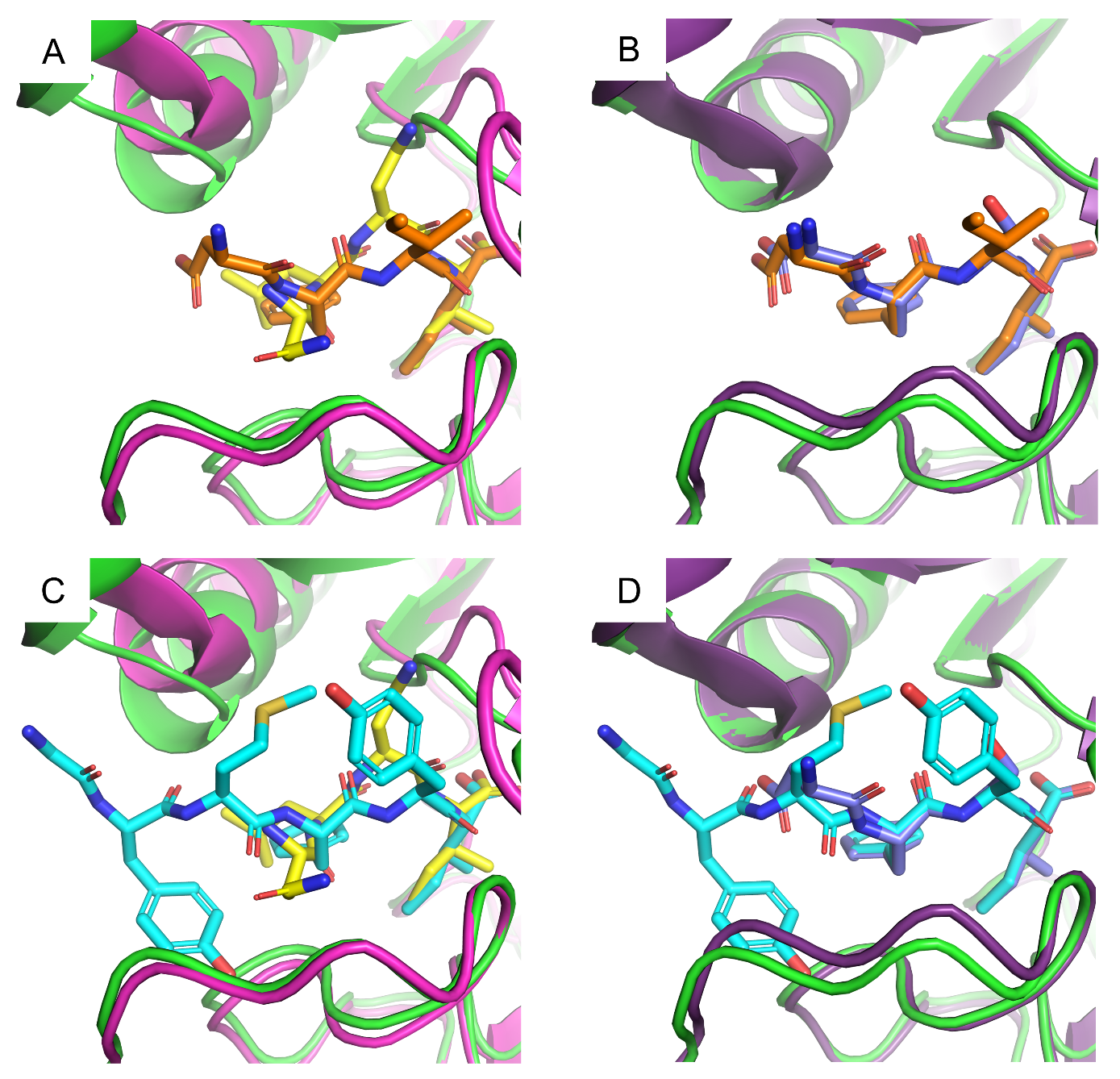

**S1 Fig. Structural comparison of *Sa*CysK:CysE 10 and *Sa*CysK:CymR 10 complexes with CysK:CysE complexes of other *H. influenzae* and *M. tuberculosis*.**

Comparison of binding modes for **A) CysE 10** and the *H. influenzae* CysE decapeptide, **B)** **CysE 10** and the *M. tuberculosis* CysE tetrapeptide, **C)** **CymR 10** and the *H. influenzae* CysE decapeptide, and **D) CymR 10** and the *M. tuberculosis* CysE tetrapeptide. **CysE 10**, **CymR 10**, the *H. influenzae* CysE decapeptide and the *M. tuberculosis* CysE tetrapeptide are shown as orange, cyan, yellow and purple sticks, respectively. *Sa*CysK is shown as green cartoon, *H. influenzae* CysK (PDB: 1Y7L) is shown as pink cartoon and *M. tuberculosis* CysK (PDB: 2Q3C) is shown as dark purple cartoon.

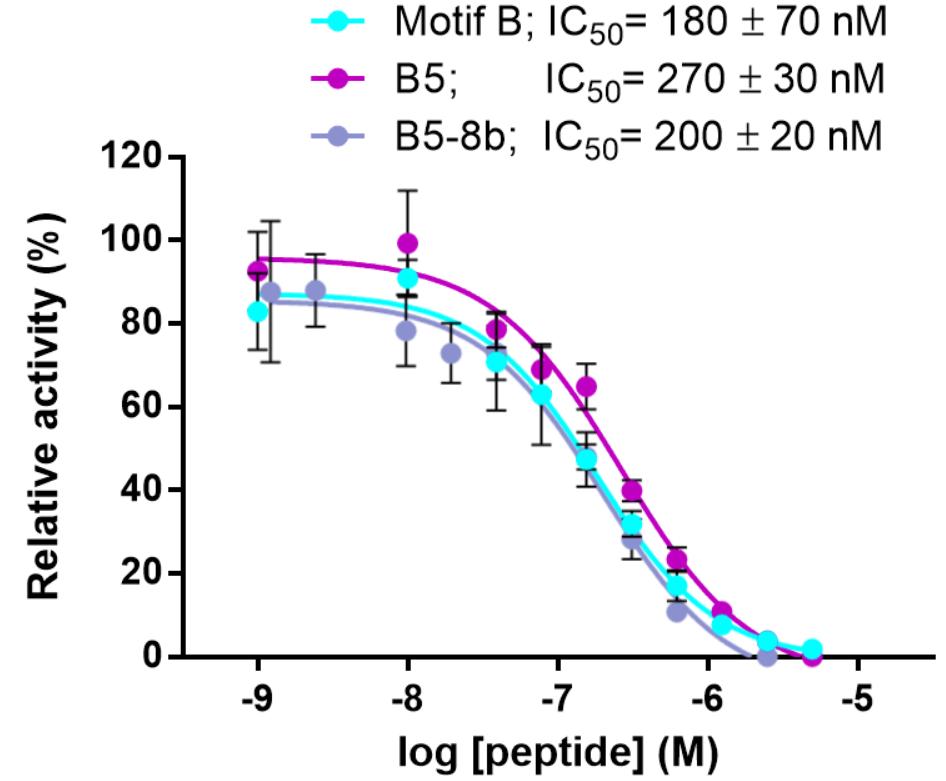

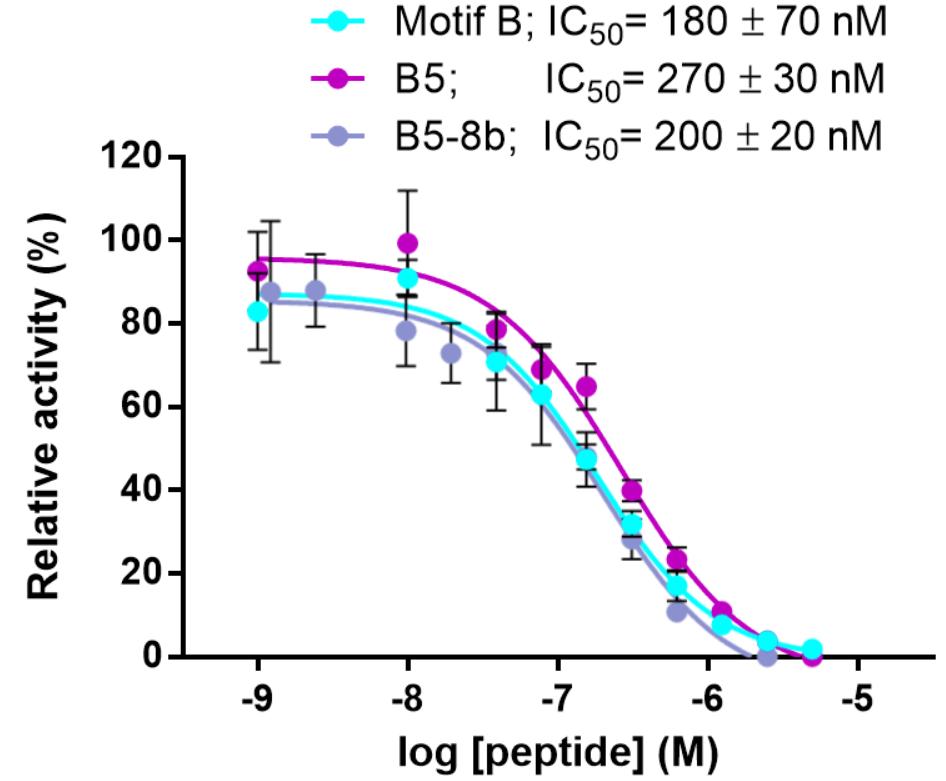

CymR 10; IC_50_ = 180 ± 70 nM

CymR 5; IC_50_ = 270 ± 30 nM

CymR 5-8b; IC_50_ = 200 ± 20 nM

**S2 Fig. Determination of IC_50_ values for *Sa*CysK inhibition by CymR 10, CymR 5 and CymR 5-8b.**

Reactions contained 50 mM Tris-HCl pH 7.5, 5 µM PLP, 1 mM sodium sulfide, 10 mM OAS, 4 nM CysK with the concentration of the **CymR 10**, **CymR 5** or **CymR 5-8b** peptide varied. Data represents the mean ± standard deviation of 3 experiments, each with a single technical replicate for all concentrations assayed.

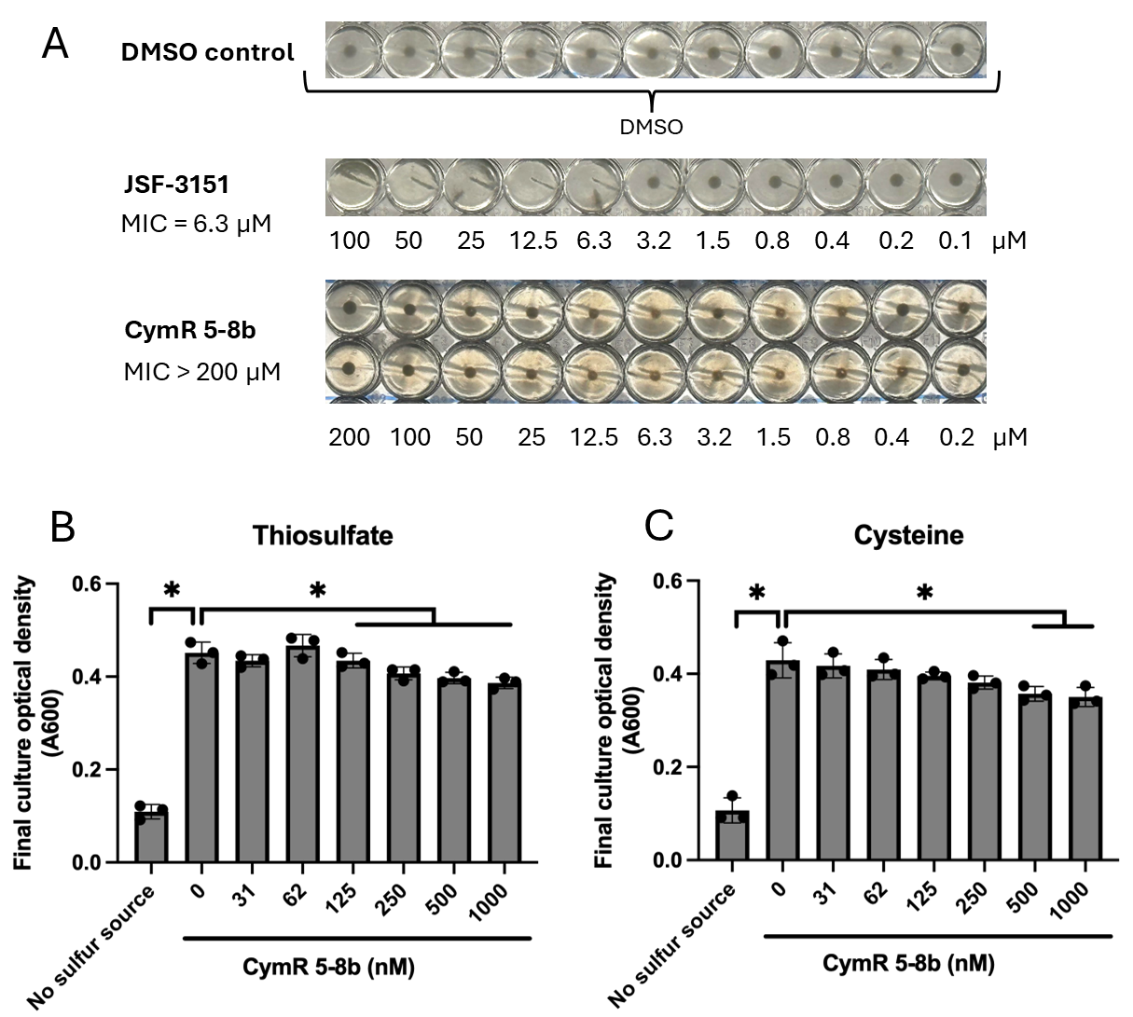

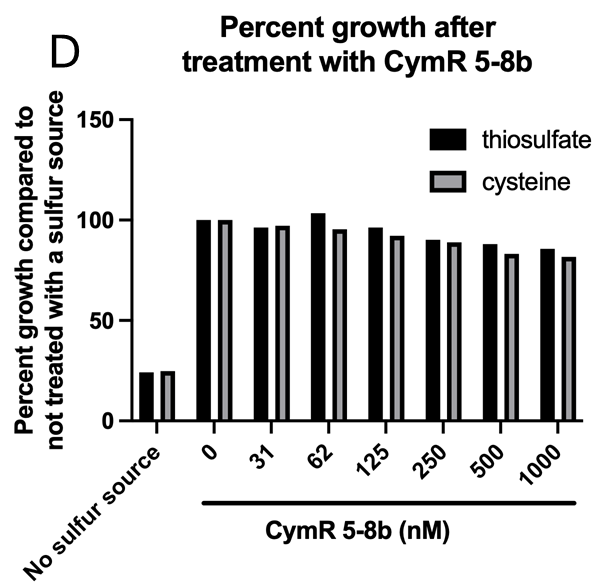

**S3 Fig. Effect of CymR 5-8b on *S. aureus* growth *in vitro*.**

**A)** Estimation of minimum inhibitory concentration for **CymR 5-8b** against *S. aureus* strain FPR3757 (ATCC BAA-1556) grown in LB. An overnight culture of bacteria was subcultured 1:100 in fresh LB media and grown at 37 °C and 180 rpm until A_600_ of 0.2 – 0.3. The bacterial culture was then diluted 1:1000 in LB media and added to a 96-well microtiter plate (100 µl per well) containing different concentrations of 2-fold serially diluted compound. DMSO-treated wells and JSF-3151) were used as the negative and positive controls, respectively. Cells were grown at 37°C shaking for 18 hours. MIC was determined as the lowest concentration of drug, resulting in complete growth inhibition by visual inspection. **B)** Effect of **CymR 5-8b** on the growth of *S. aureus* strain LAC with or without thiosulfate as a sulfur source. **C)** Effect of **CymR 5-8b** on *S. aureus* LAC grown with or without cysteine. **D)** The percent growth was determined using the data presented in panels B and C. For panels B and C, *S. aureus* was inoculated to a starting OD_600_ of 0.01 in a chemically defined media containing no sulfur source, 50 µM thiosulfate, or 50 µM cysteine. Cells were grown in 200 µL cultures in a 96-well assay plate at 37°C shaking at 200rpm for 18 hours. The data displayed represent the average final optical density (A_600_) of biological triplicates with standard deviations shown. Student’s t-tests were performed on the data, and * indicates p < 0.05.

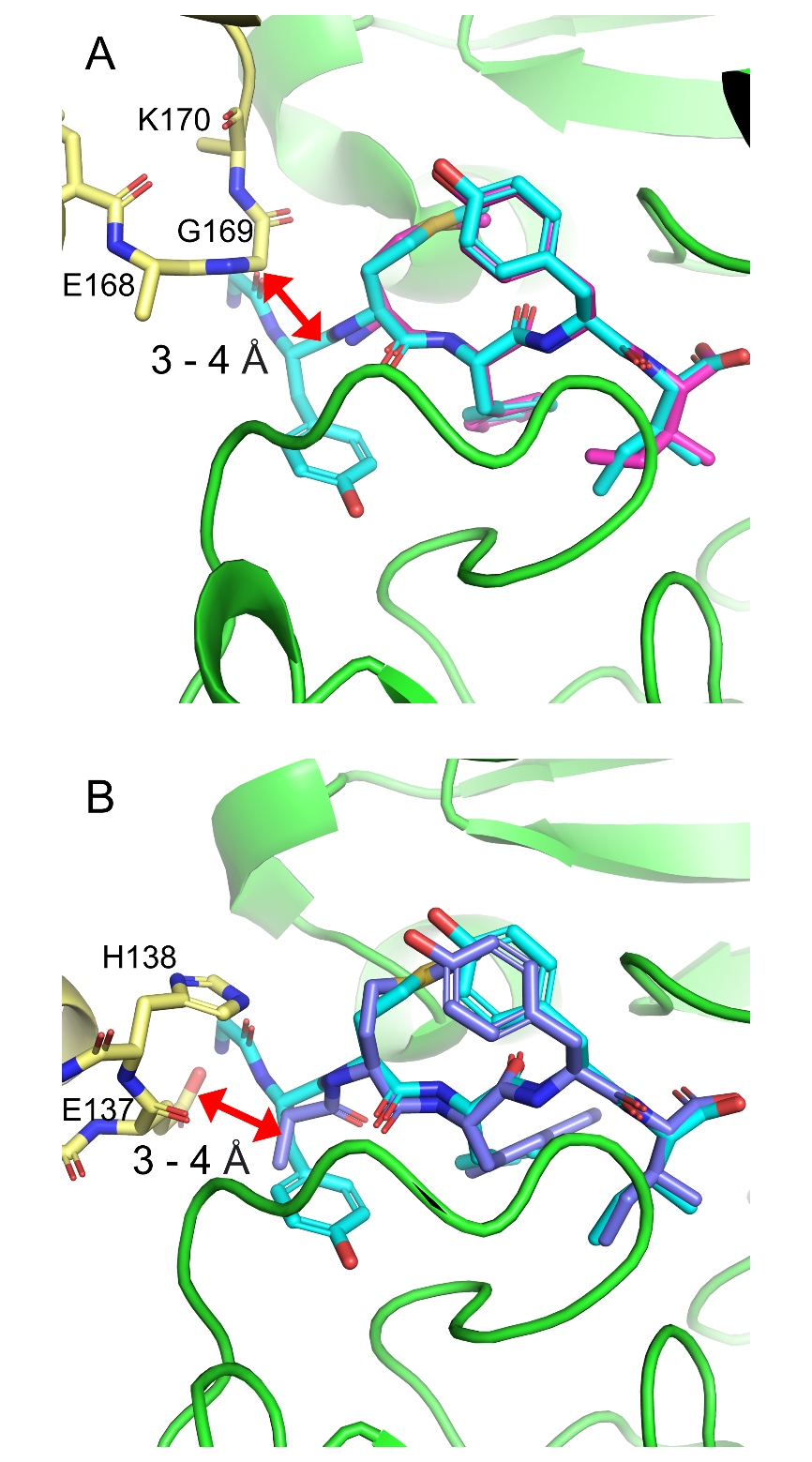

**S4 Fig. Crystal contacts surrounding Y_6_ for** **CymR 5** and **CymR 5-b cocrystal structures.**

**A)** Crystal contacts observed in Chain A of the **CymR 5** cocrystal structure, and **B)** Crystal contacts observed in Chains C and D of the **CymR 5-8b** cocrystal structure. In both cases the crystal contacts clash with the position Y_6_ was observed to occupy in the **CymR 10** cocrystal structure. The *Sa*CysK monomer binding **CymR 5** or **CymR 5-8b** is shown as green cartoon, with the *Sa*CysK monomer forming the crystal contact shown as yellow cartoon. **CymR 5**, **CymR 5-8b** and **CymR 10** are shown as pink, purple and cyan sticks, respectively. The site of the clash is indicated by the red arrows, with the approximate distance from the crystal contact reported in angstroms.

**S1 Table. Peptide characterization data.**

| **Peptide** | **Sequence** | **Molecular formula** | **Theoretical MW**  (g/mol) | **Observed MW** ^a^  (g/mol) | **Purity** ^b^  (%) |
| --- | --- | --- | --- | --- | --- |
| CymR 10 | EDLDGYMFYI | C_59_H_80_N_10_O_19_S_1_ | 1265.4 | 1265.2 | 95 |
| CysE 10 | NGEIQDDYII | C_51_H_78_N_12_O_20_ | 1179.2 | 1178.7 | 99 |
| CymR 6 | GYMFYI | C_40_H_52_N_6_O_9_S_1_ | 793.0 | 792.4 | 93 |
| CymR 5 | YMFYI | C_38_H_49_N_5_O_8_S_1_ | 735.9 | 735.3 | 94 |
| CymR 4 | MFYI | C_29_H_40_N_4_O_6_S_1_ | 572.7 | 572.3 | 94 |
| CymR 5-6 | AMFYI | C_32_H_45_N_5_O_7_S_1_ | 643.8 | 643.5 | 93 |
| CymR 5-7 | YAFYI | C_36_H_45_N_5_O_8_ | 675.8 | 675.4 | 98 |
| CymR 5-8 | YMAYI | C_32_H_45_N_5_O_8_S_1_ | 659.8 | 659.4 | 98 |
| CymR 5-9 | YMFAI | C_32_H_45_N_5_O_7_S_1_ | 643.8 | 643.4 | 88 |
| CymR 5-10 | YMFYA | C_35_H_43_N_5_O_8_S_1_ | 693.8 | 693.4 | 98 |
| CymR 5-6a | (4-F-Phe)MFYI | C_38_H_48_F_1_N_5_O_7_S_1_ | 737.9 | 737.5 | 98 |
| CymR 5-6b | (4-Cl-Phe)MFYI | C_38_H_48_Cl_1_N_5_O_7_S_1_ | 754.3 | 753.4 | 97 |
| CymR 5-7a | YYFYI | C_42_H_49_N_5_O_9_ | 767.9 | 767.5 | 99 |
| CymR 5-7b | YFFYI | C_42_H_49_N_5_O_8_ | 751.9 | 753 | 97 |
| CymR 5-7c | Y(Nle)FYI | C_39_H_51_N_5_O_8_ | 717.9 | 717.5 | 98 |
| CymR 5-7d | Y(MetO_2_)FYI | C_38_H_49_N_5_O_10_S_1_ | 767.9 | 767.4 | 95 |
| CymR 5-8a | YMWYI | C_40_H_50_N_6_O_8_S_1_ | 774.9 | 774.5 | 98 |
| CymR 5-8b | YM(Nal)YI | C_42_H_51_N_5_O_8_S_1_ | 786.0 | 785.3 | 96 |

^a^ Determined by supplier using ESI-MS analysis

^b^ Determined by supplier using HPLC analysis

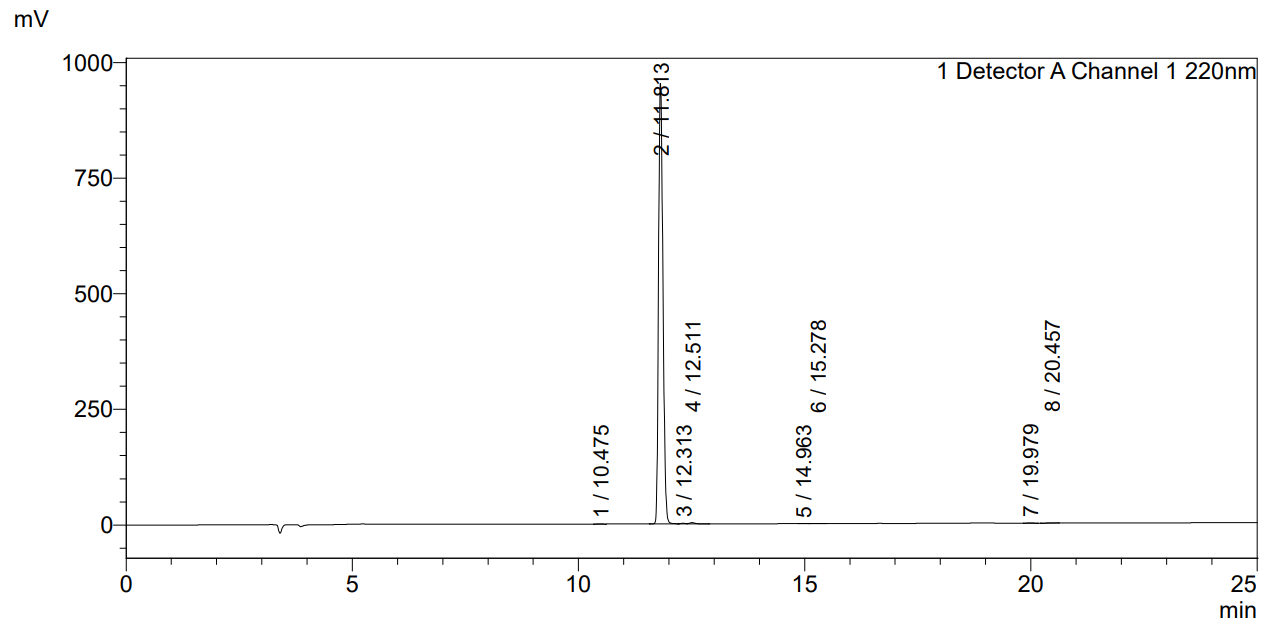

**HPLC analysis**

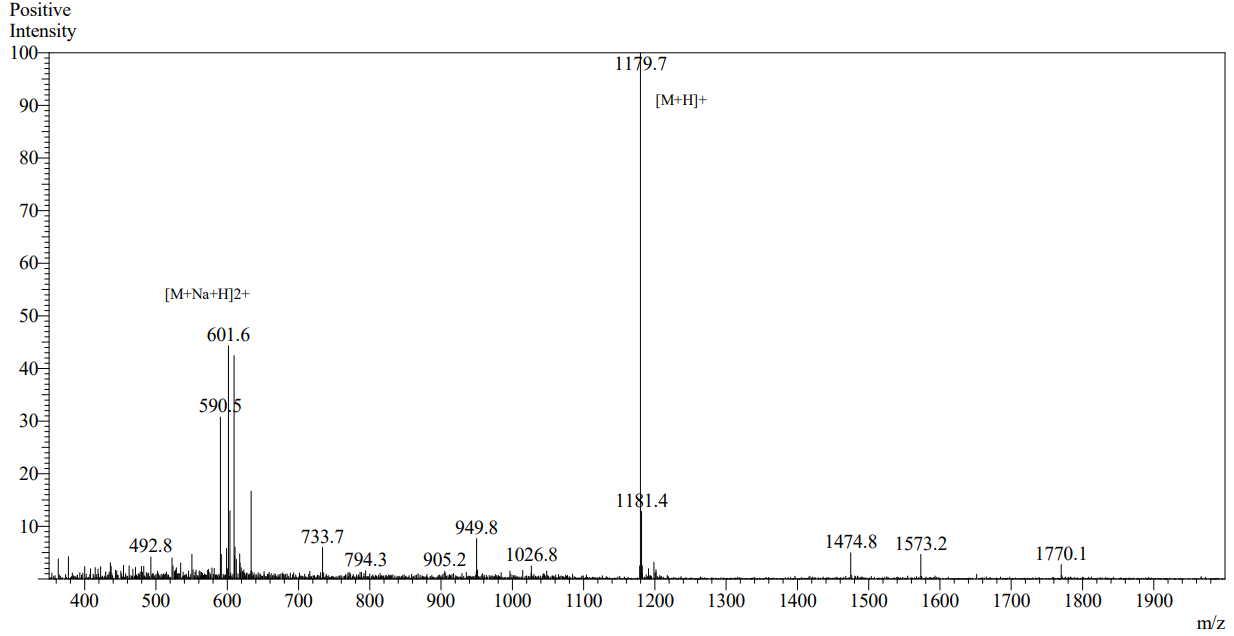

**MS analysis**

**CysE 10**

**S5 Fig. MS and HPLC analysis for key peptides investigated.**

The peak corresponding to the respective peptide is indicated by the red arrow.

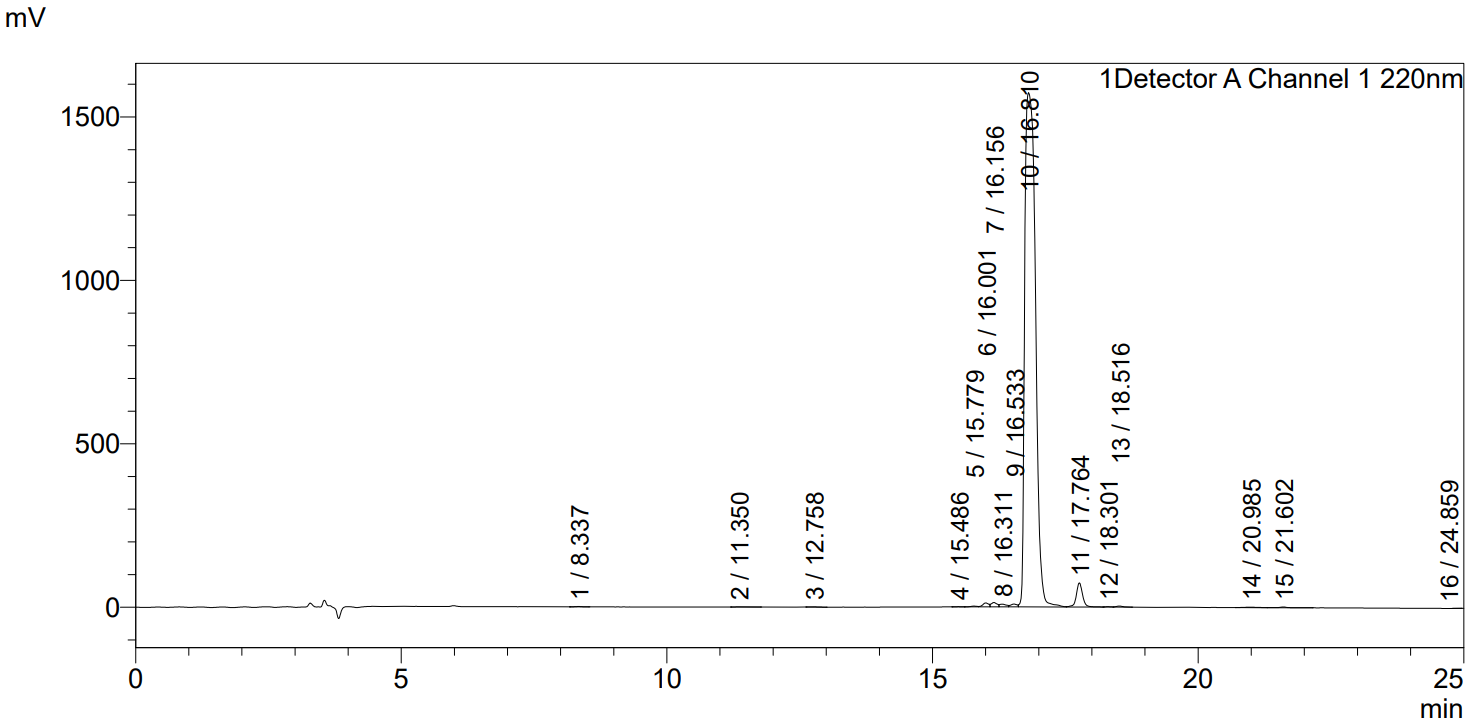

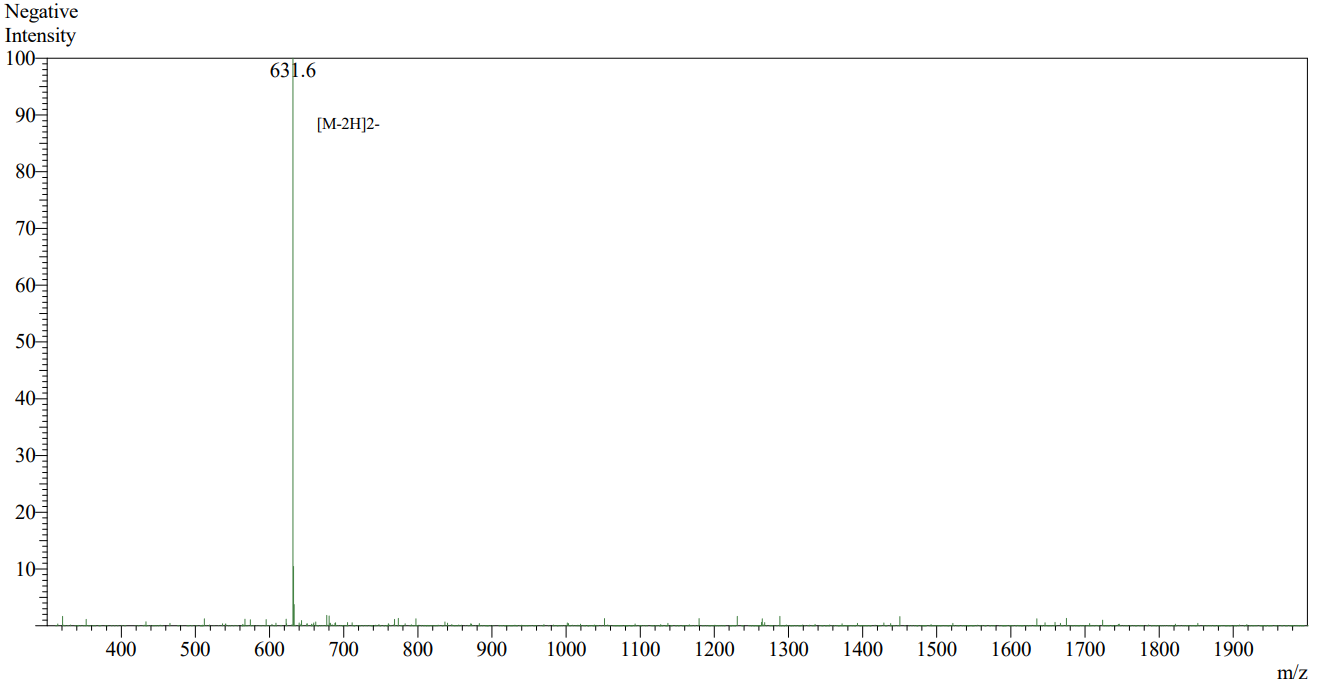

**MS analysis**

**HPLC analysis**

**CymR 10**

**S5 Fig. MS and HPLC analysis for key peptides investigated.** (continued)

The peak corresponding to the respective peptide is indicated by the red arrow.

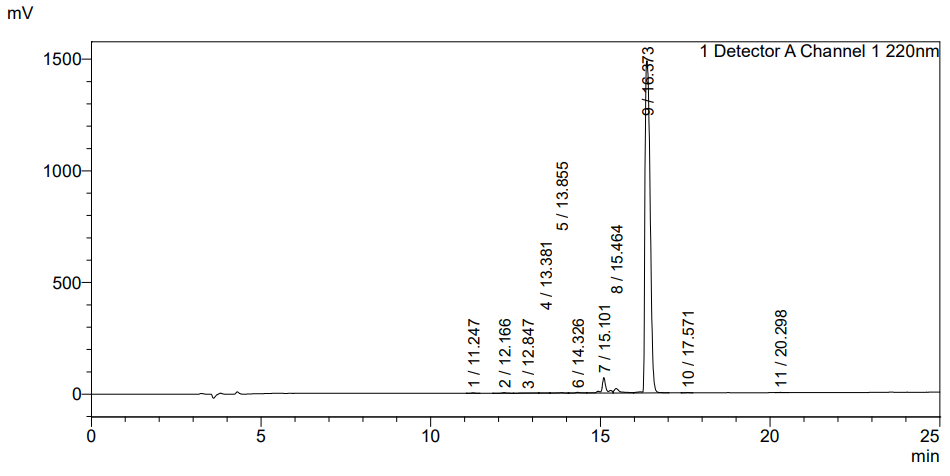

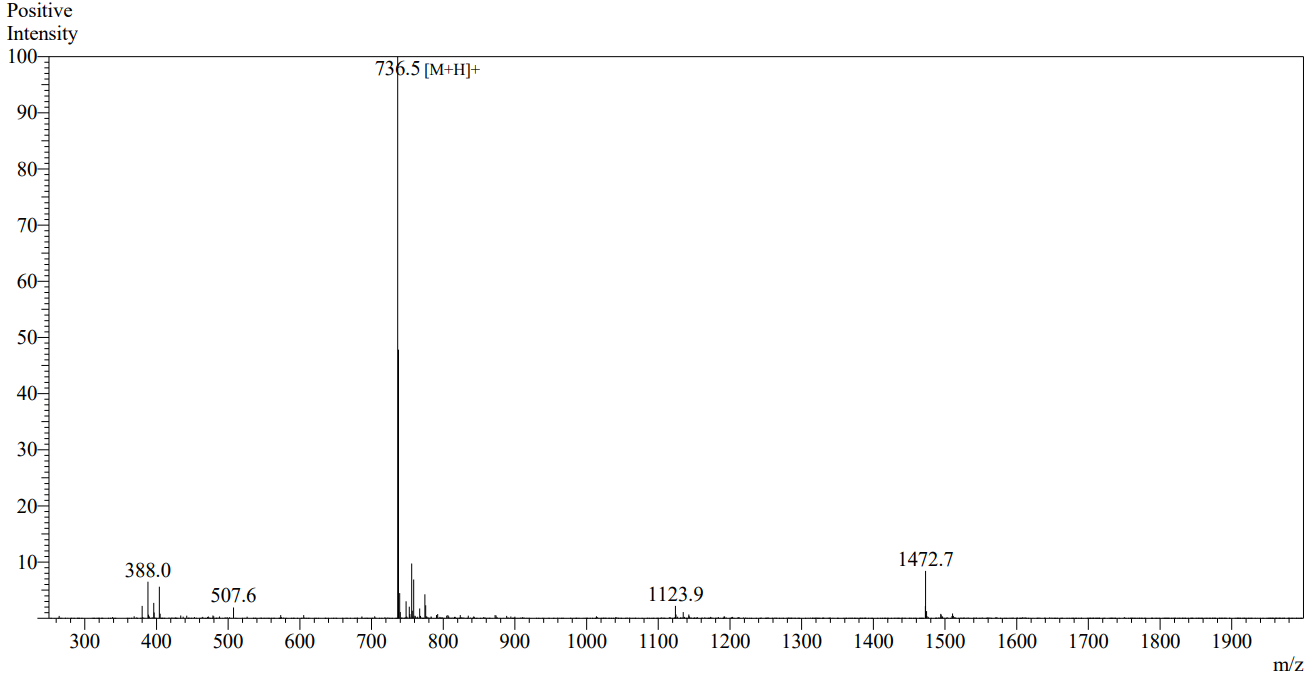

**HPLC analysis**

**MS analysis**

**CymR 5**

**S5 Fig. MS and HPLC analysis for key peptides investigated.** (continued)

The peak corresponding to the respective peptide is indicated by the red arrow.

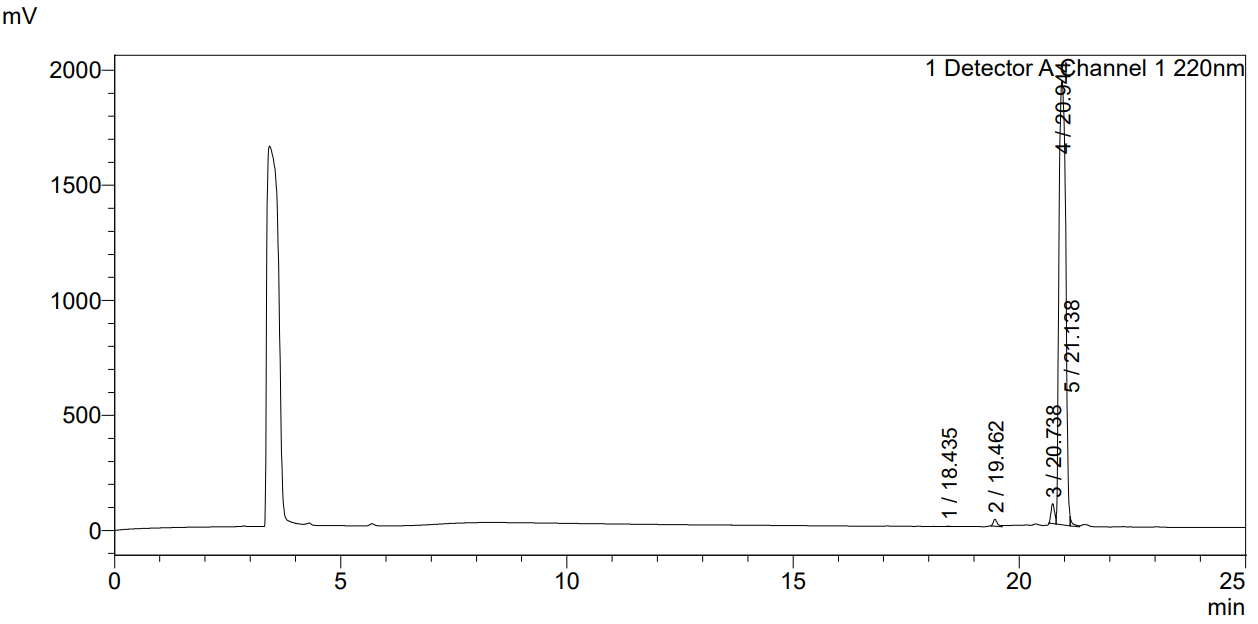

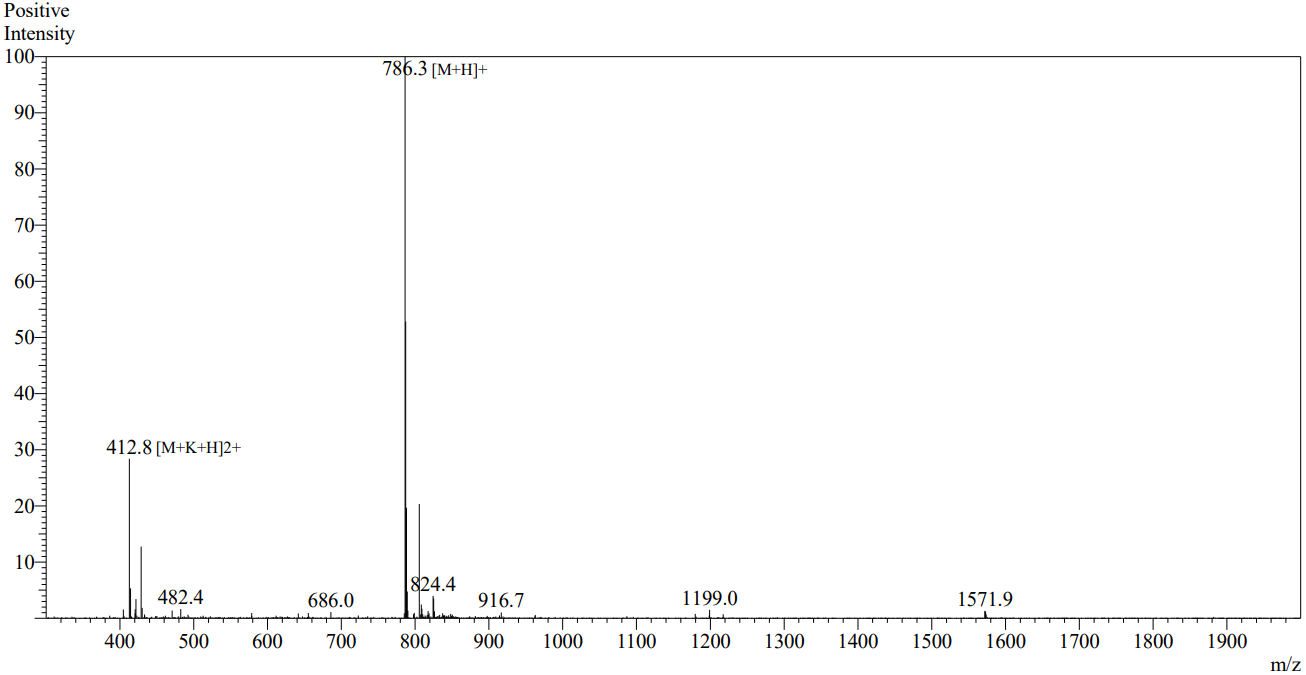

**CymR 5-8b**

**MS analysis**

**HPLC analysis**

**S5 Fig. MS and HPLC analysis for key peptides investigated.** (continued)

The peak corresponding to the respective peptide is indicated by the red arrow.

**S2 Table. Crystallographic data collection, processing and refinement statistics.**

| **Statistic** | **CysK**  **holo**  PDB: 8SRT | **CysK**  **CysE 10**  PDB: 8SRU | **CysK**  **CymR 10**  PDB: 8SRV | **CysK**  **CymR 5**  PDB: 8T2C | **CysK**  **CymR 5-8b**  PDB: 8SRW |
| --- | --- | --- | --- | --- | --- |
| **Wavelength (Å)** | 0.9537 | 0.9537 | 0.9537 | 0.9537 | 0.9537 |
| **Resolution range (Å)** | 35.95 - 1.9 (1.97 - 1.9)^a^ | 38.71 - 1.5 (1.55 - 1.5) | 35.42 - 1.32  (1.37 - 1.32) | 39.19 - 1.8 (1.86 - 1.8) | 39.50 - 2.15 (2.23 - 2.15) |
| **Space group** | P 1 2_1_ 1 | P 1 2_1_ 1 | P 1 2_1_ 1 | P 1 2_1_ 1 | P 1 2_1_ 1 |
| **Unit cell** |  |  |  |  |  |
| a, b, c (Å) | 55.0 96.0 58.5 | 64.7 96.7 94.3 | 56.4 96.2 59.0 | 50.6 94.5 56.1 | 65.9 119.7 84.7 |
| α, β, γ (^o^) | 90 112.3 90 | 90 93.0 90 | 90 111.9 90 | 90 108.5 90 | 90 98.71 90 |
| **Total reflections** | 301264 (29055) | 1256260 (125007) | 918783  (79848) | 319177 (32899) | 494012 (50723) |
| **Unique reflections** | 44270 (4412) | 184494 (18294) | 136382 (13521) | 46230  (4588) | 70279 (7032) |
| **Multiplicity** | 6.8 (6.6) | 6.8 (6.8) | 6.7 (5.9) | 6.9 (7.2) | 7.0 (7.2) |
| **Completeness (%)** | 99.79 (99.84) | 99.77 (99.46) | 99.78 (99.40) | 99.73 (99.89) | 99.54 (99.86) |
| **Mean I/sigma(I)** | 13.01 (1.87) | 12.36 (1.59) | 14.58 (1.16) | 9.83 (1.24) | 8.86 (1.07) |
| **Wilson B-factor** | 38.22 | 24.00 | 19.16 | 31.04 | 33.28 |
| **R-merge** | 0.063 (0.939) | 0.062 (1.168) | 0.056(1.139) | 0.094 (1.573) | 0.213 (1.792) |
| **R-meas** | 0.0684 (1.02) | 0.067 (1.264) | 0.060(1.252) | 0.102 (1.694) | 0.230 (1.931) |
| **R-pim** | 0.026 (0.394) | 0.026 (0.480) | 0.023(0.507) | 0.039(0.624) | 0.086 (0.715) |
| **CC1/2** | 0.999 (0.766) | 0.999 (0.637) | 0.999 (0.567) | 0.999 (0.686) | 0.994 (0.446) |
| **Reflections used** | 44259 (4410) | 184435(18292) | 136347(13508) | 46186 (4584) | 70237 (7028) |
| **in refinement** |  |  |  |  |  |
| **Reflections used** | 2214 (244) | 9657 (1060) | 6982 (666) | 1997 (198) | 3517 (350) |
| **for R-free** |  |  |  |  |  |
| **R-work** | 0.188 (0.264) | 0.185 (0.266) | 0.165(0.277) | 0.223 (0.488) | 0.194 (0.283) |
| **R-free** | 0.220 (0.305) | 0.219 (0.296) | 0.188 (0.297) | 0.245 (0.504) | 0.236 (0.315) |
| **Number of** | 4722 | 10462 | 5495 | 4610 | 9909 |
| **non-hydrogen atoms** |  |  |  |  |  |
| **macromolecules** | 4489 | 9333 | 4806 | 4462 | 9349 |
| **solvent** | 233 | 1093 | 689 | 148 | 560 |
| **Protein residues** | 618 | 1252 | 633 | 619 | 1248 |
| **RMS(bonds)** | 0.004 | 0.016 | 0.008 | 0.003 | 0.002 |
| **RMS(angles)** | 0.84 | 1.43 | 1.05 | 0.58 | 0.53 |
| **Ramachandran** | 97.86 | 97.22 | 97.58 | 97.36 | 98.26 |
| **favored (%)** |  |  |  |  |  |
| **Ramachandran** | 2.14 | 2.78 | 2.42 | 2.64 | 1.74 |
| **allowed (%)** |  |  |  |  |  |
| **Ramachandran** | 0.00 | 0.00 | 0.00 | 0.00 | 0.00 |
| **outliers (%)** |  |  |  |  |  |
| **Rotamer outliers (%)** | 0.00 | 0.00 | 0.00 | 0.46 | 1.36 |
| **Clashscore** | 5.17 | 3.64 | 2.18 | 3.85 | 3.85 |
| **Average B-factor** | 43.73 | 29.48 | 24.99 | 46.38 | 36.07 |
| **macromolecules** | 43.59 | 28.53 | 23.50 | 46.31 | 35.86 |
| **solvent** | 46.57 | 37.54 | 35.36 | 48.39 | 39.48 |

^a^ Closed brackets contain values for the highest resolution shell.

**CymR 6**

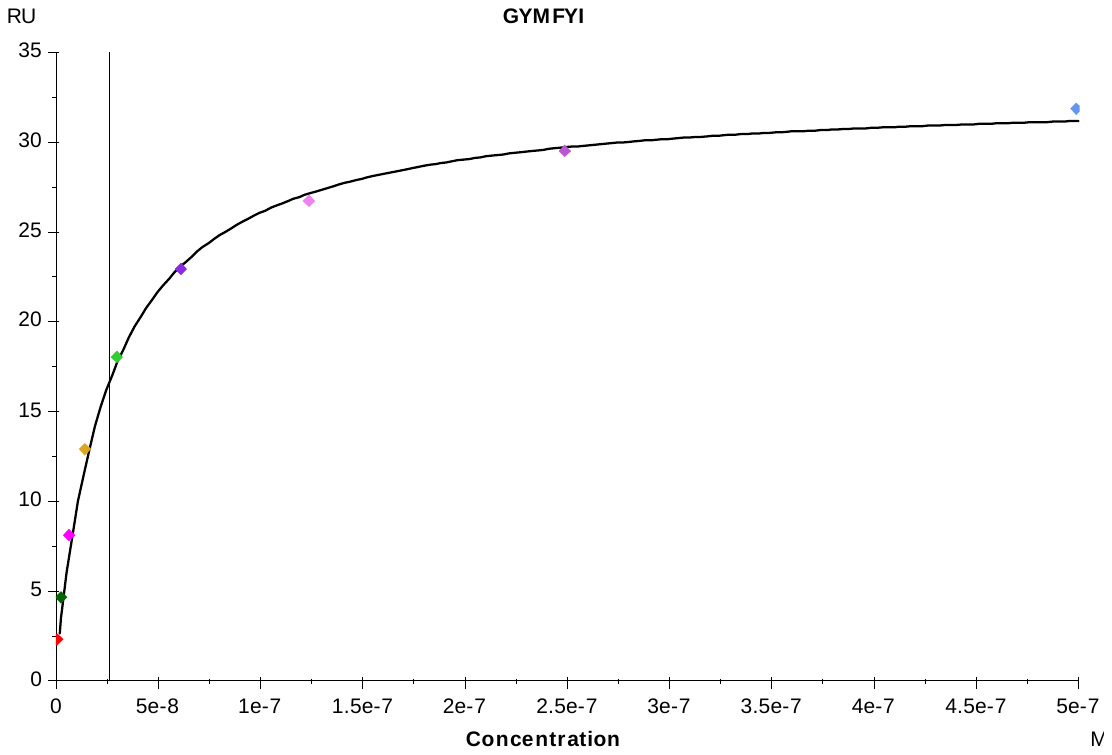

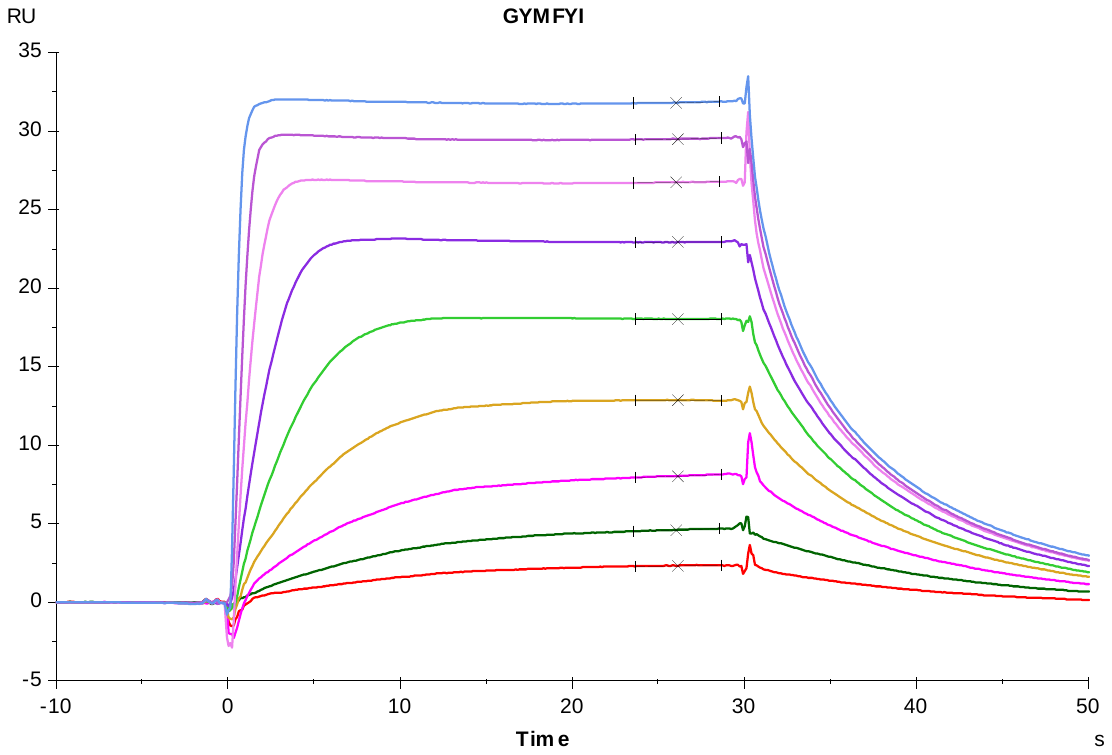

**S6 Fig. Representative sensorgrams (Top) and steady-state binding responses (Bottom) for CymR derived peptides.**

The vertical line indicates estimated equilibrium dissociation constant (K_D_).

**CymR 5**

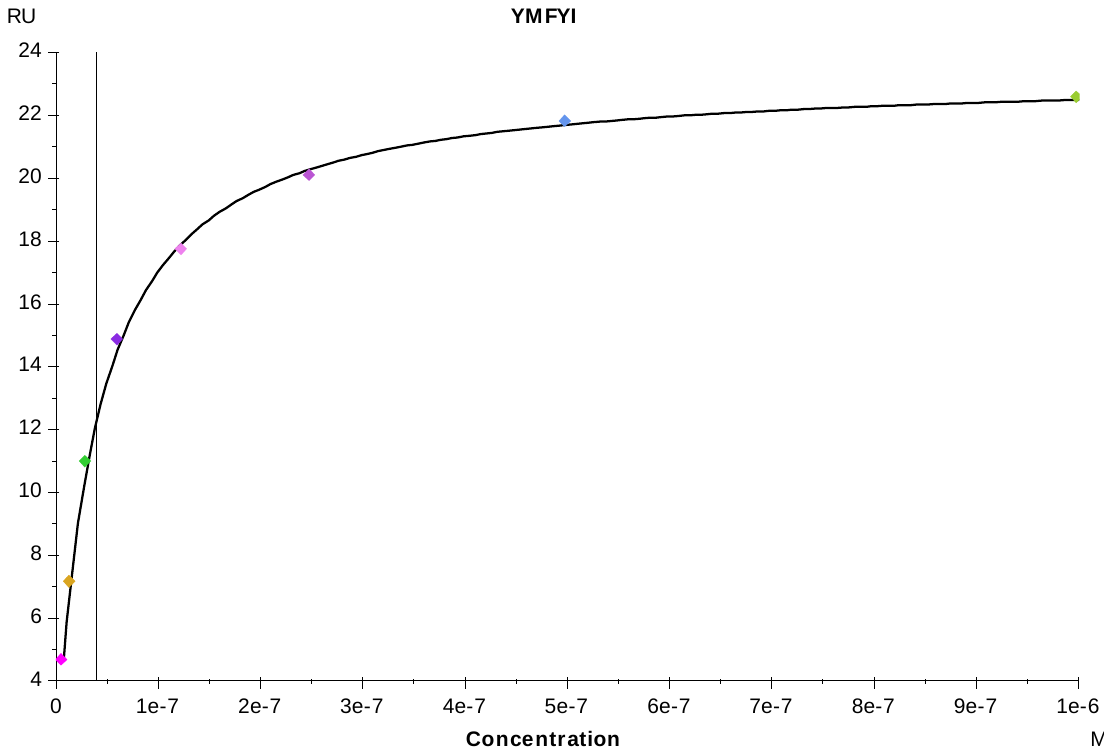

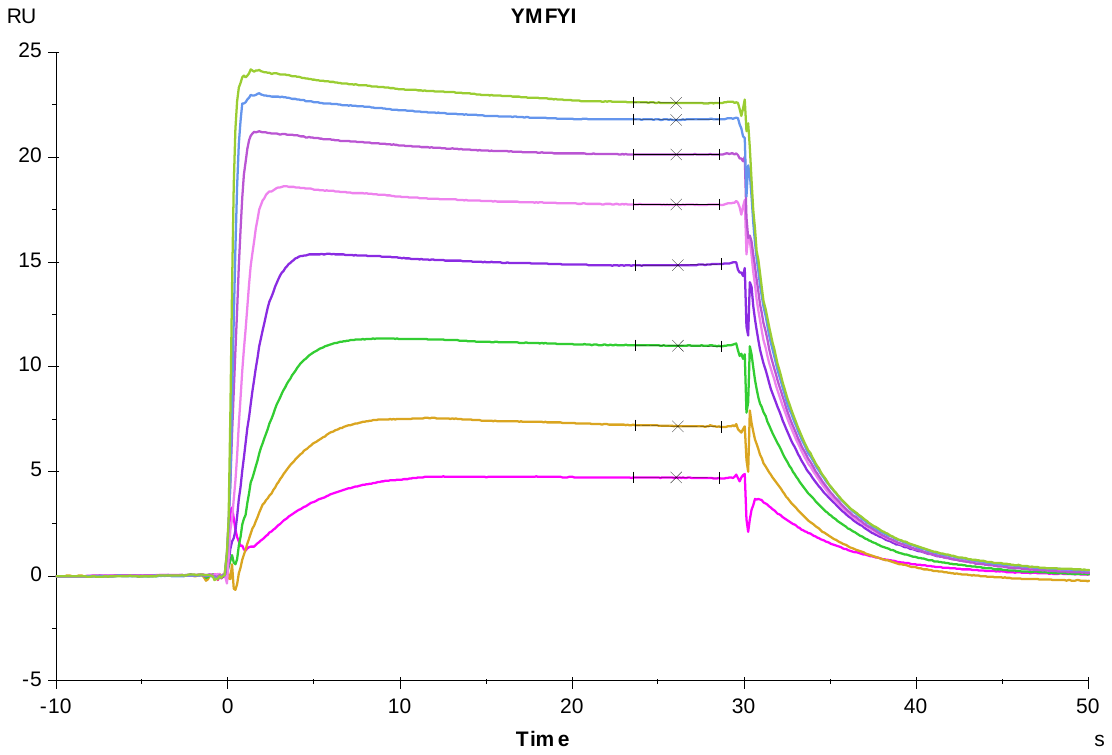

**S6 Fig. Representative sensorgrams (Top) and steady-state binding responses (Bottom) for CymR derived peptides.** (continued)

**CymR 4**

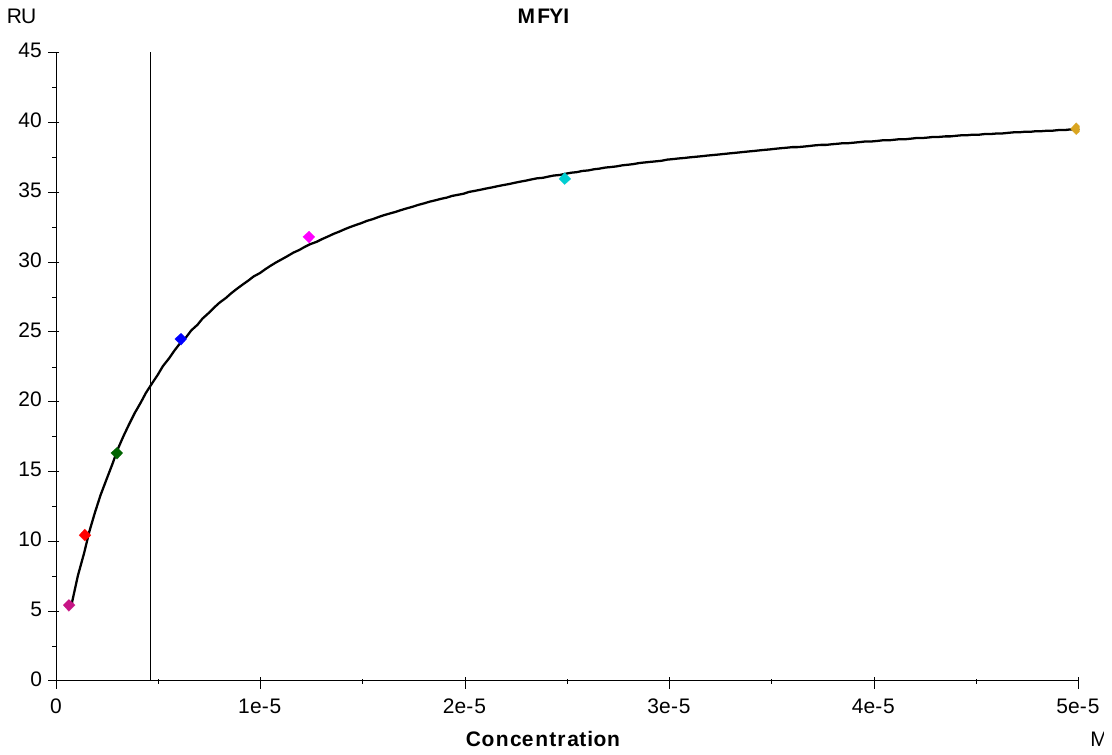

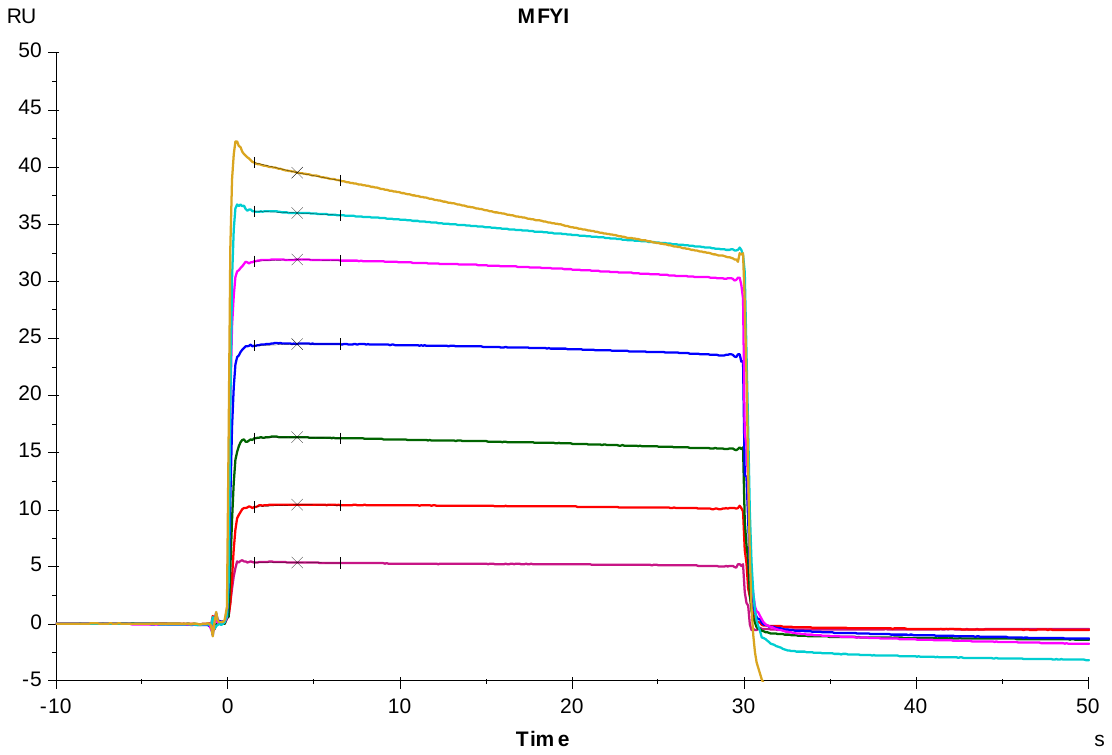

**S6 Fig. Representative sensorgrams (Top) and steady-state binding responses (Bottom) for CymR derived peptides.** (continued)

**CymR 5-6**

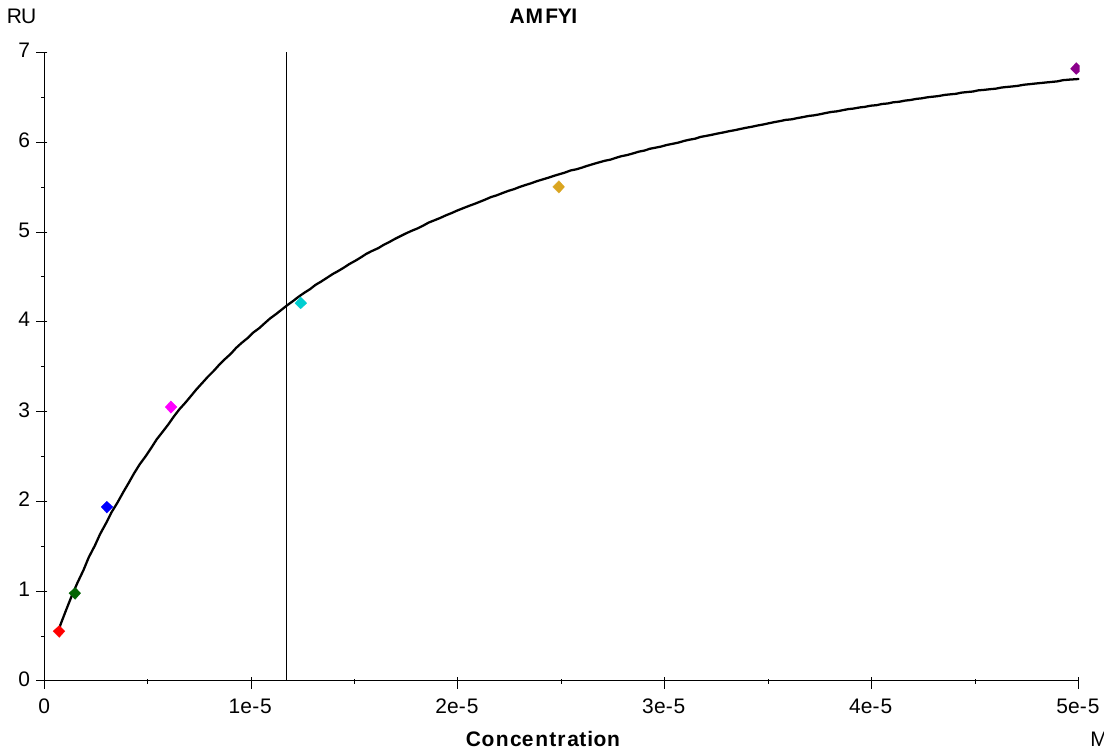

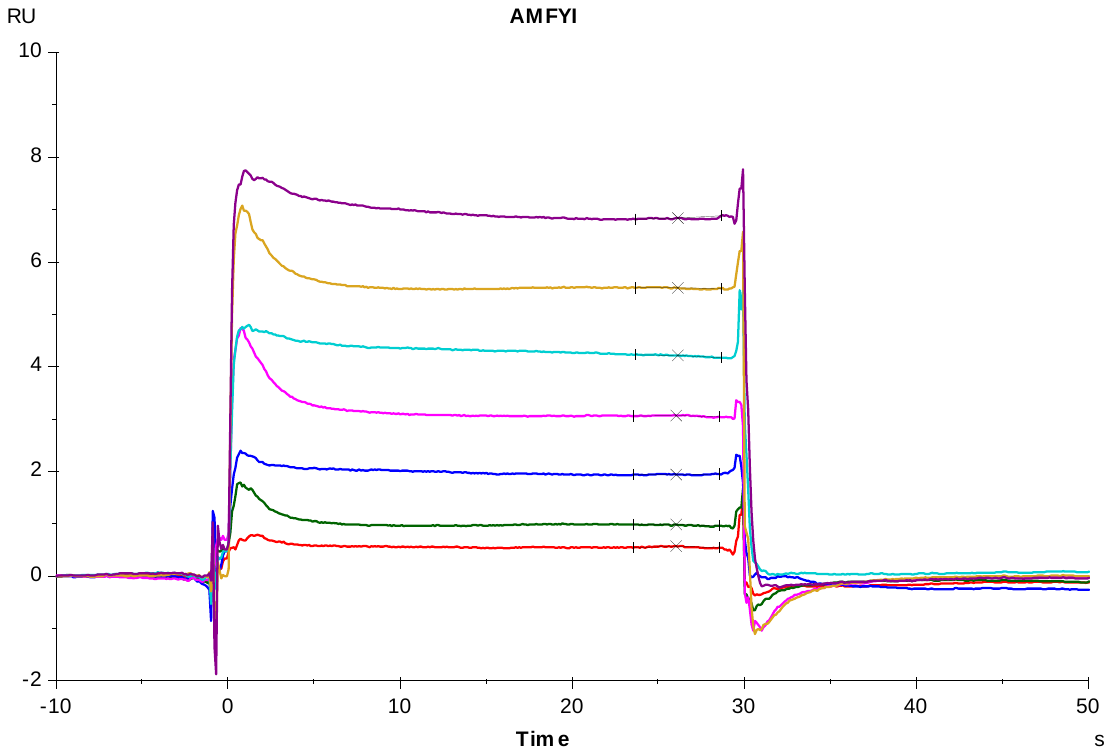

**S6 Fig. Representative sensorgrams (Top) and steady-state binding responses (Bottom) for CymR derived peptides.** (continued)

**CymR 5-7**

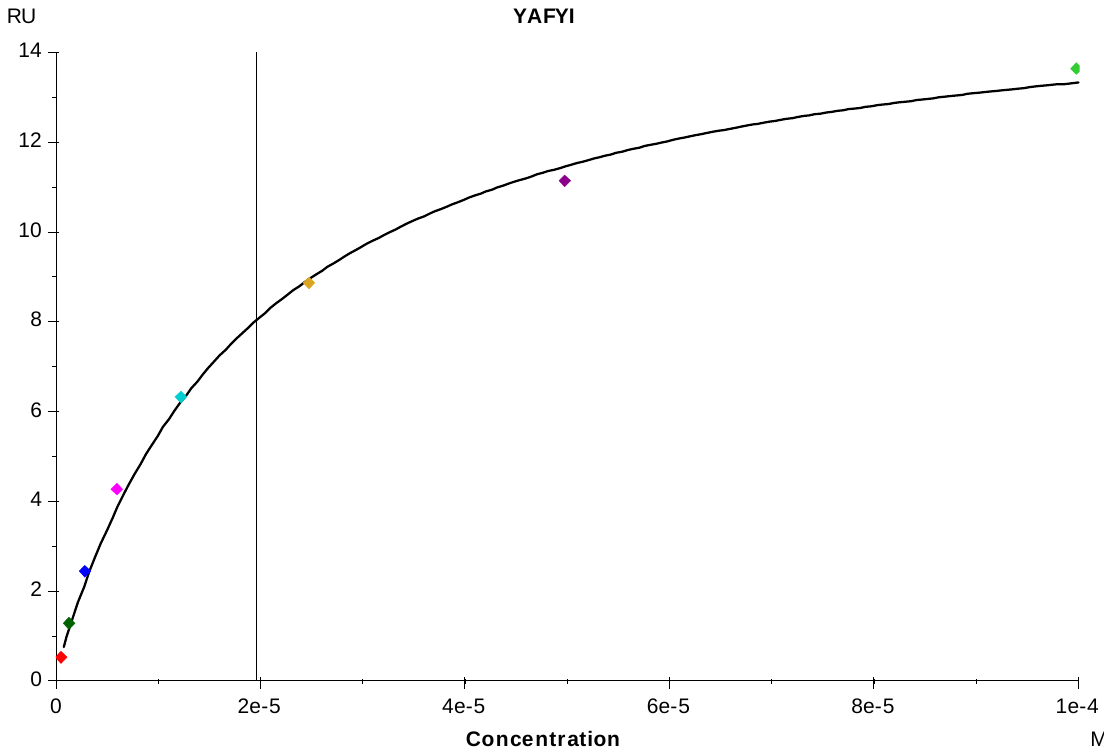

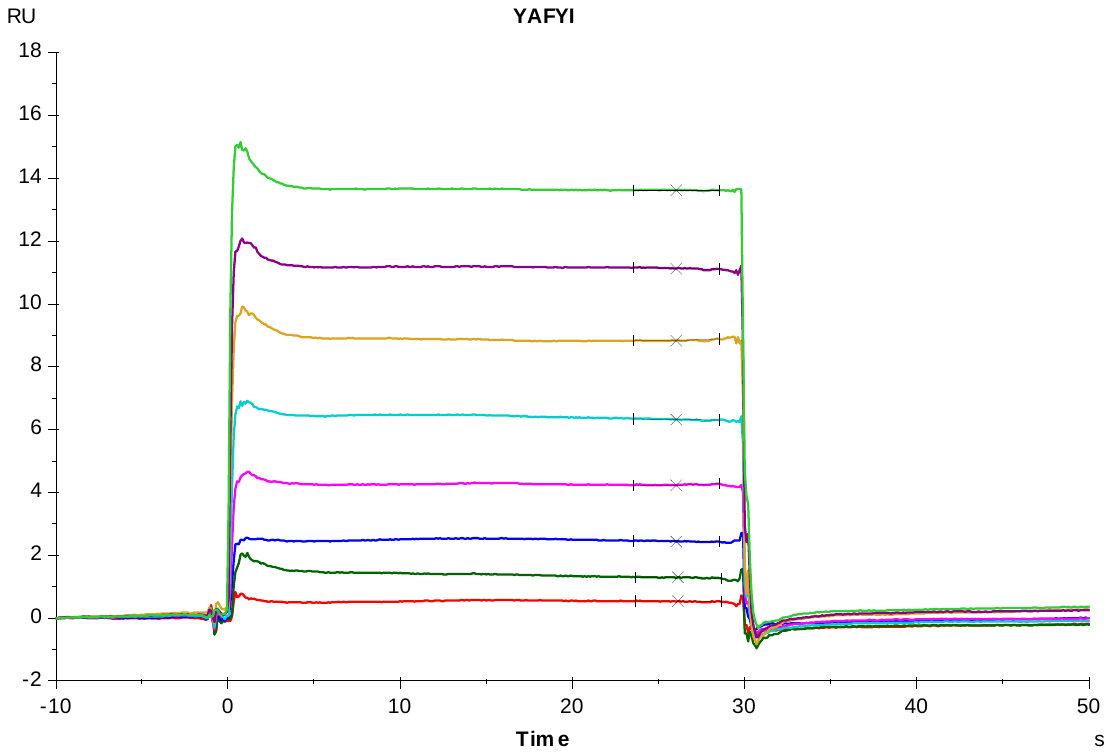

**S6 Fig. Representative sensorgrams (Top) and steady-state binding responses (Bottom) for CymR derived peptides.** (continued)

**CymR 5-8**

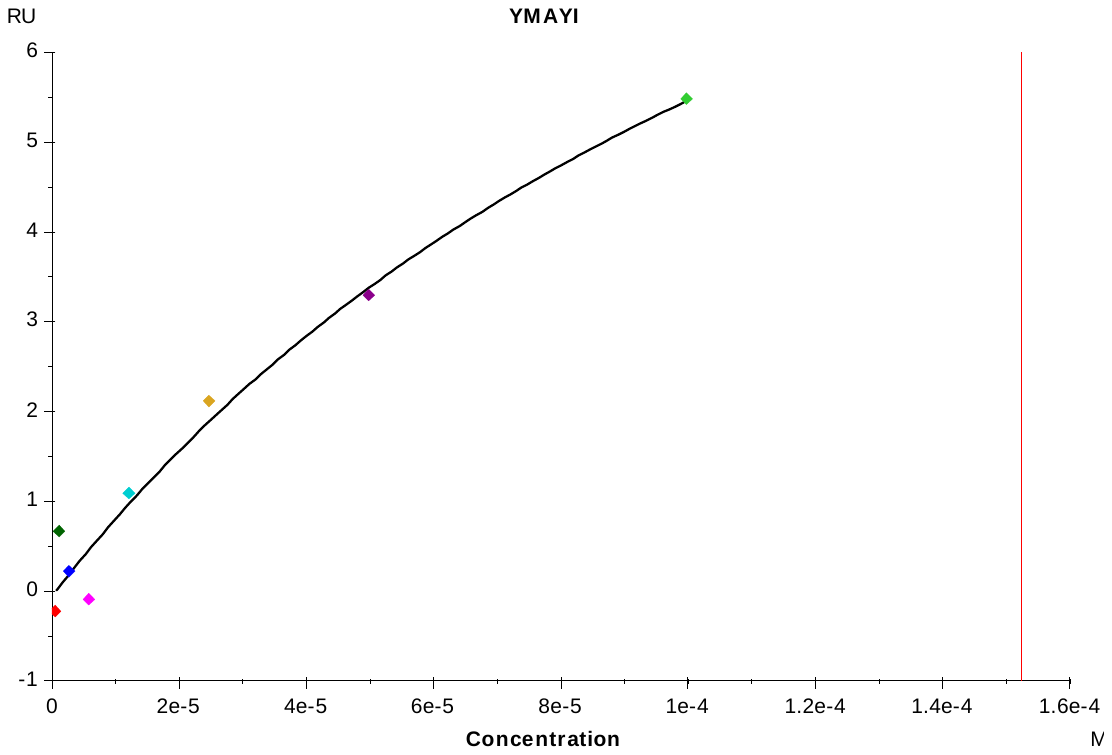

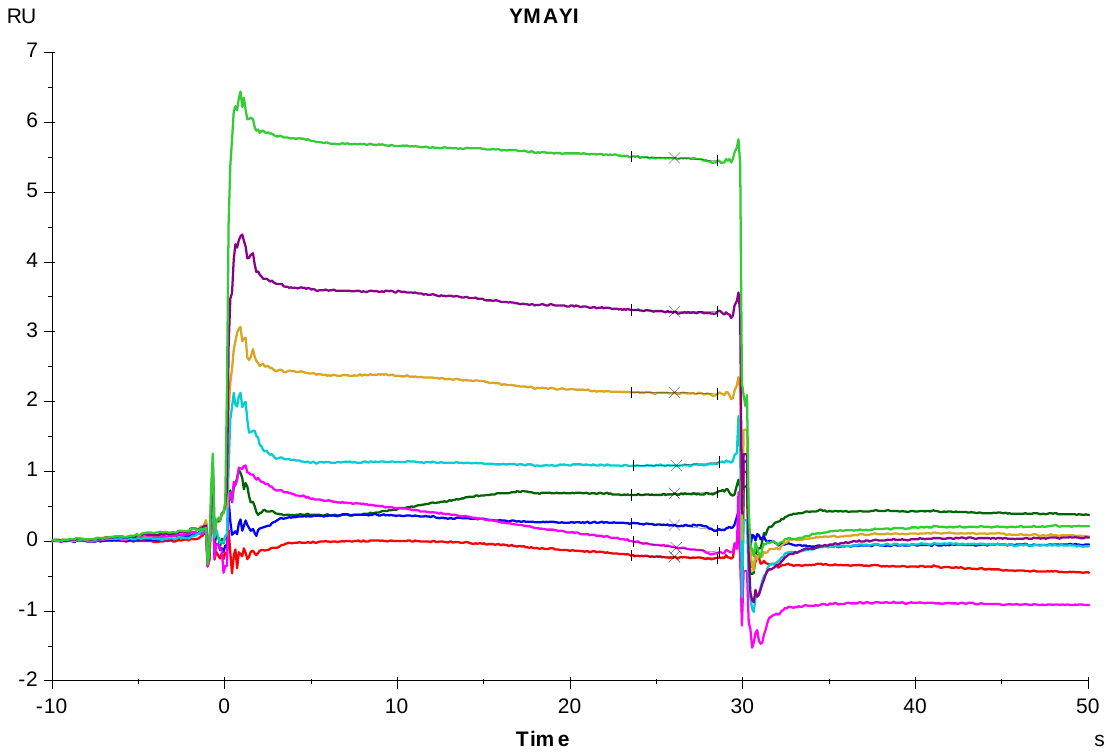

**S6 Fig. Representative sensorgrams (Top) and steady-state binding responses (Bottom) for CymR derived peptides.** (continued)

**CymR 5-9**

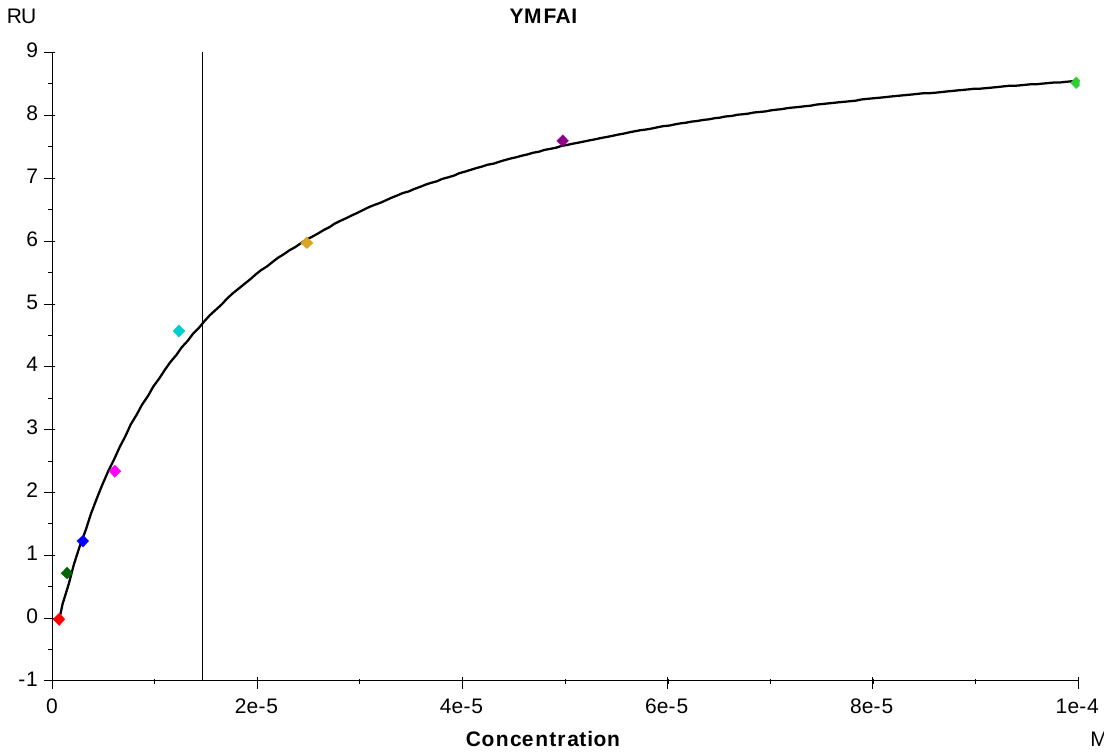

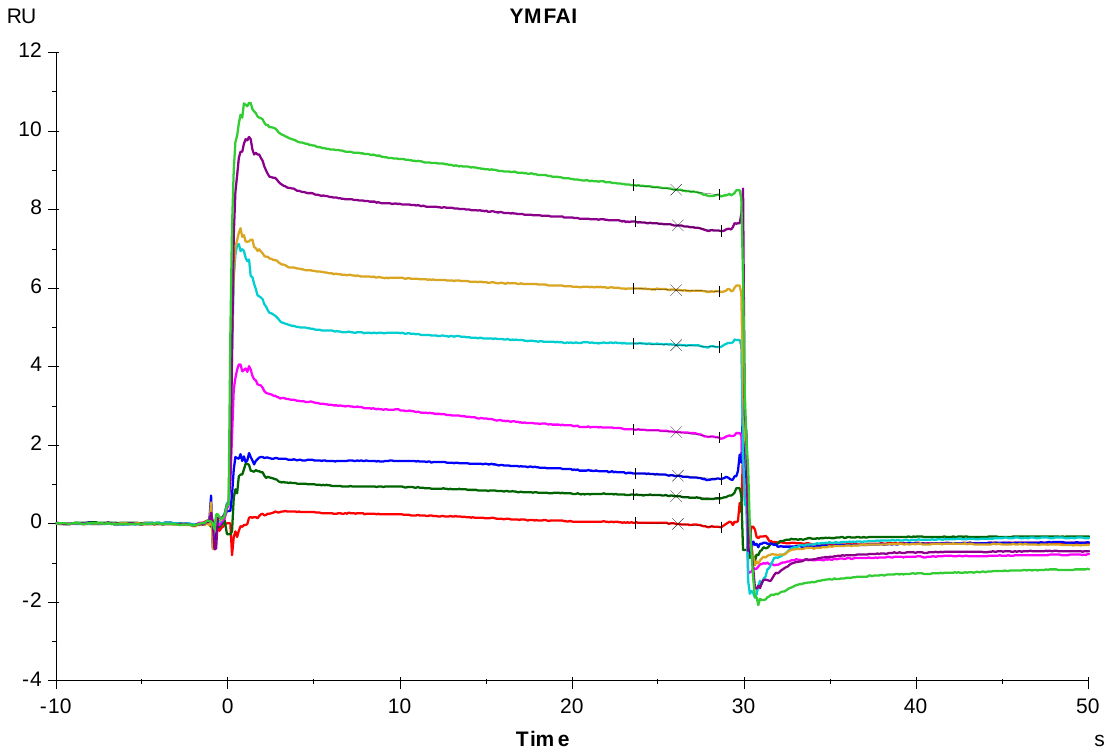

**S6 Fig. Representative sensorgrams (Top) and steady-state binding responses (Bottom) for CymR derived peptides.** (continued)

**CymR 5-10**

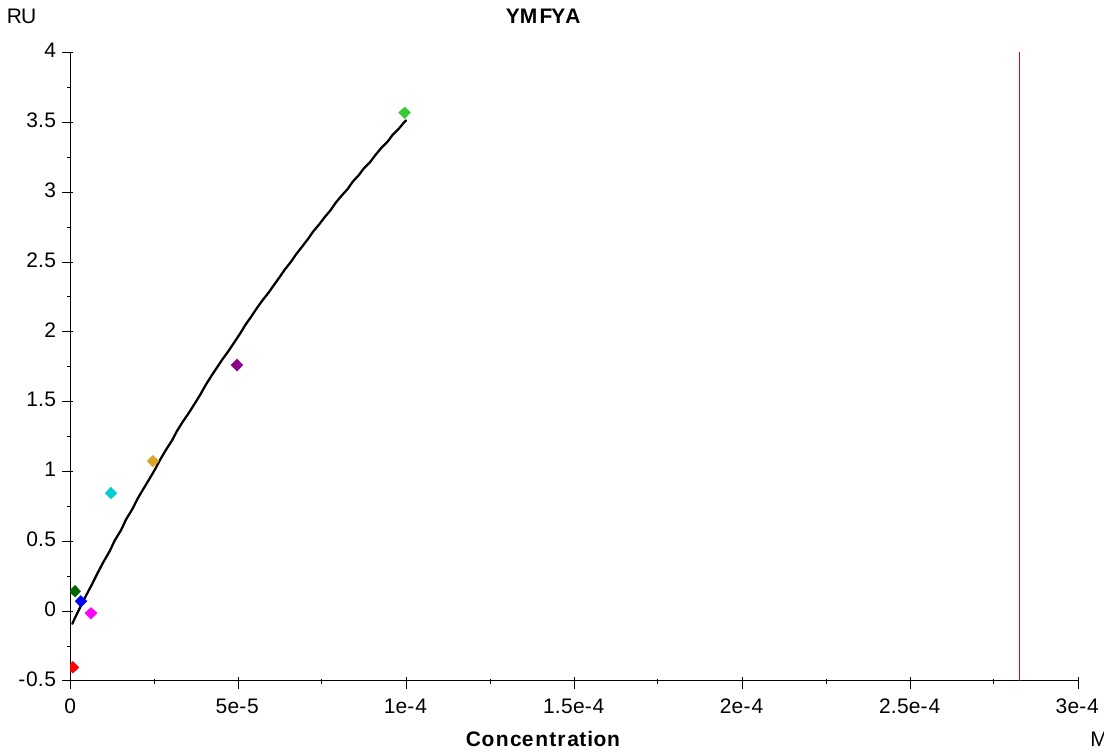

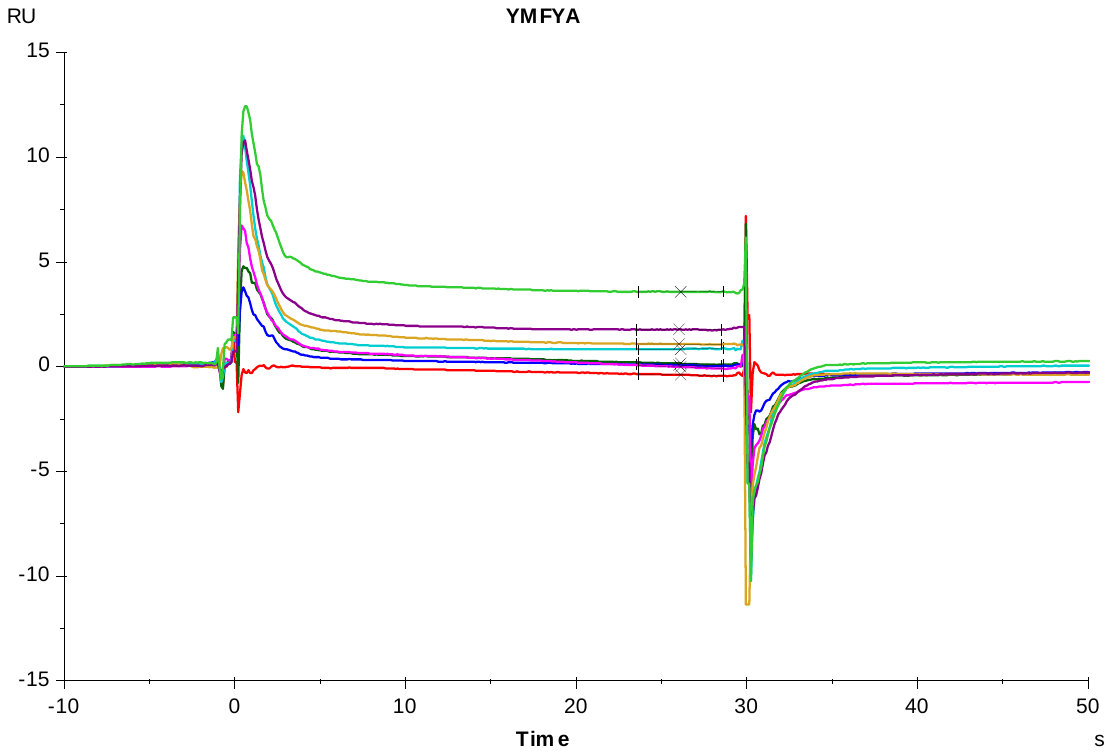

**S6 Fig. Representative sensorgrams (Top) and steady-state binding responses (Bottom) for CymR derived peptides.** (continued)

**CymR 5-6a**

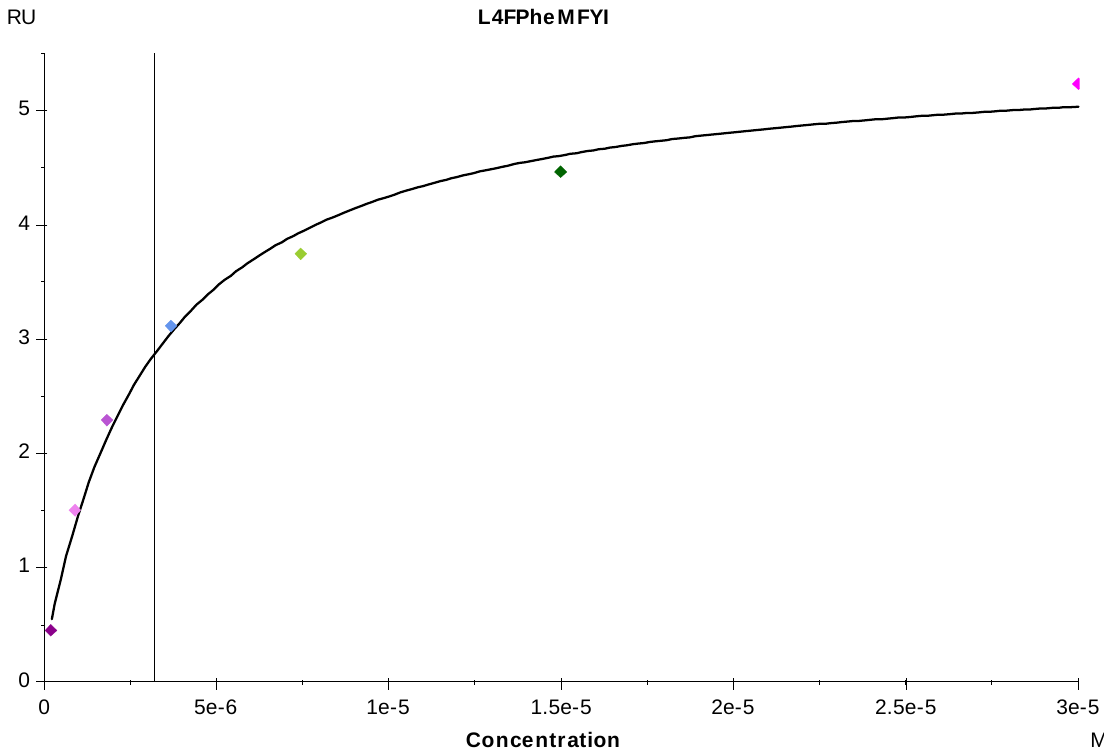

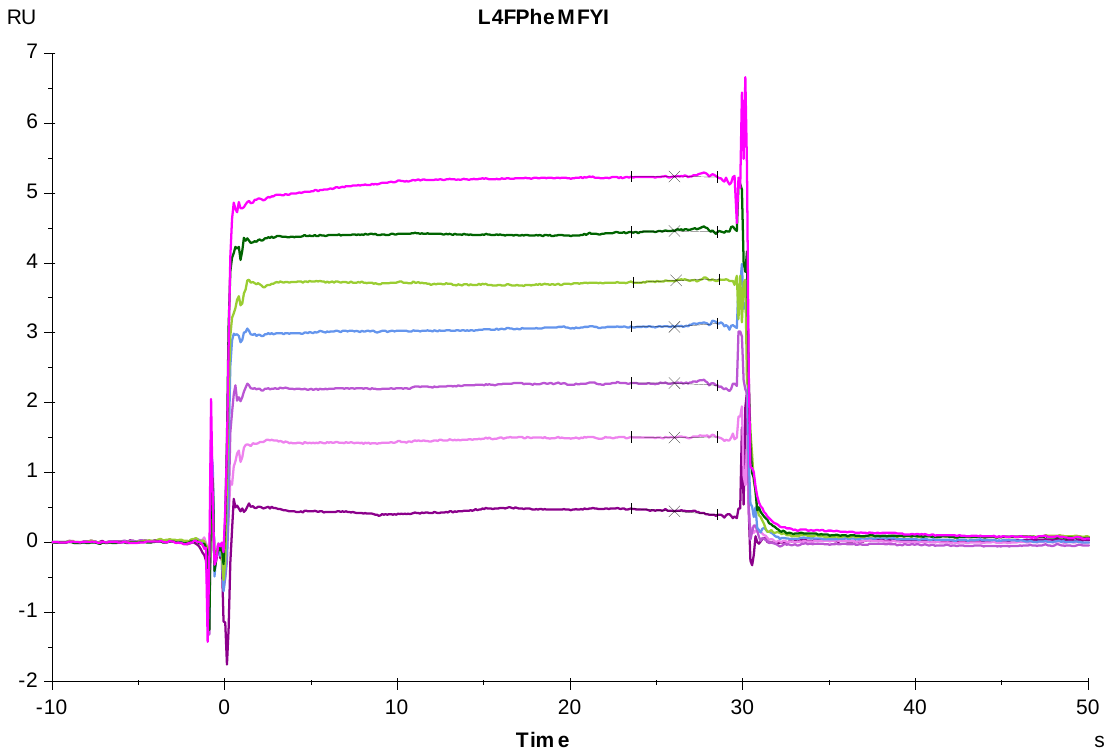

**S6 Fig. Representative sensorgrams (Top) and steady-state binding responses (Bottom) for CymR derived peptides.** (continued)

**CymR 5-6b**

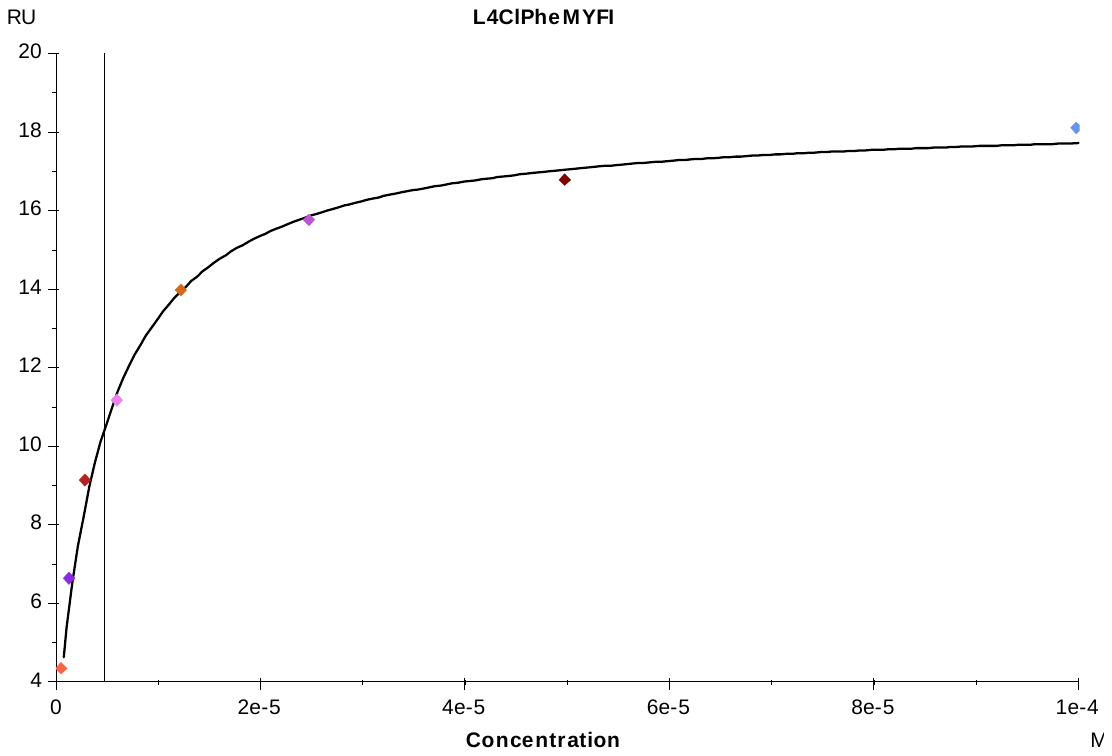

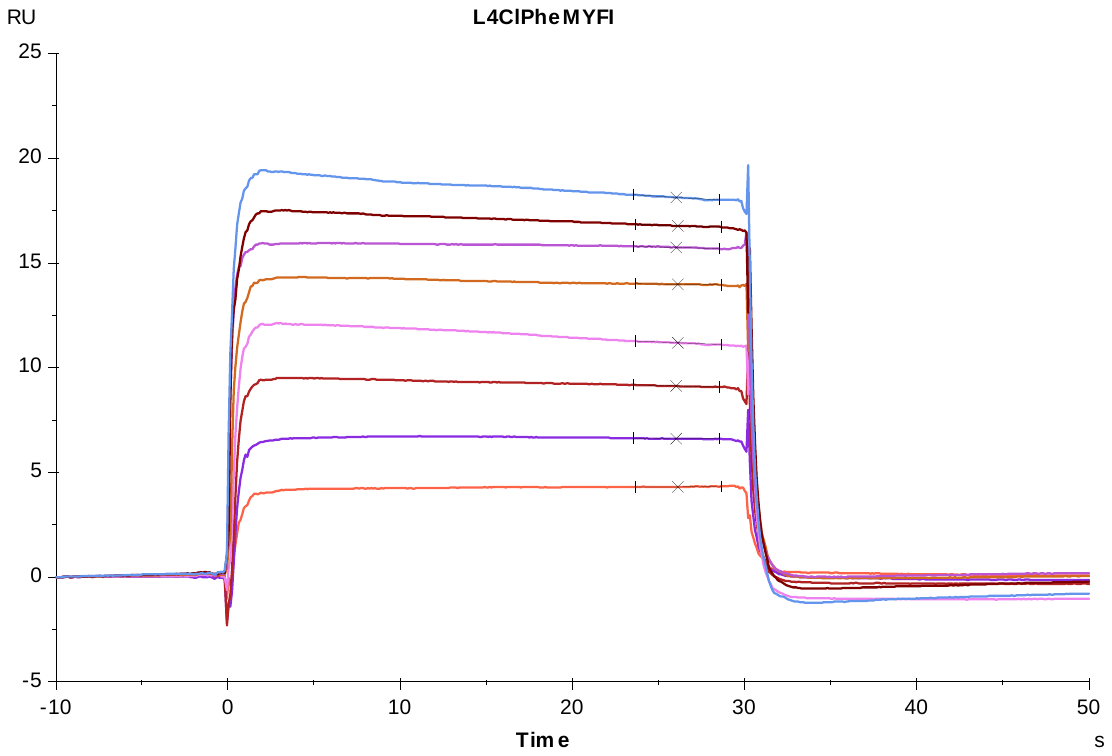

**S6 Fig. Representative sensorgrams (Top) and steady-state binding responses (Bottom) for CymR derived peptides.** (continued)

**CymR 5-7a**

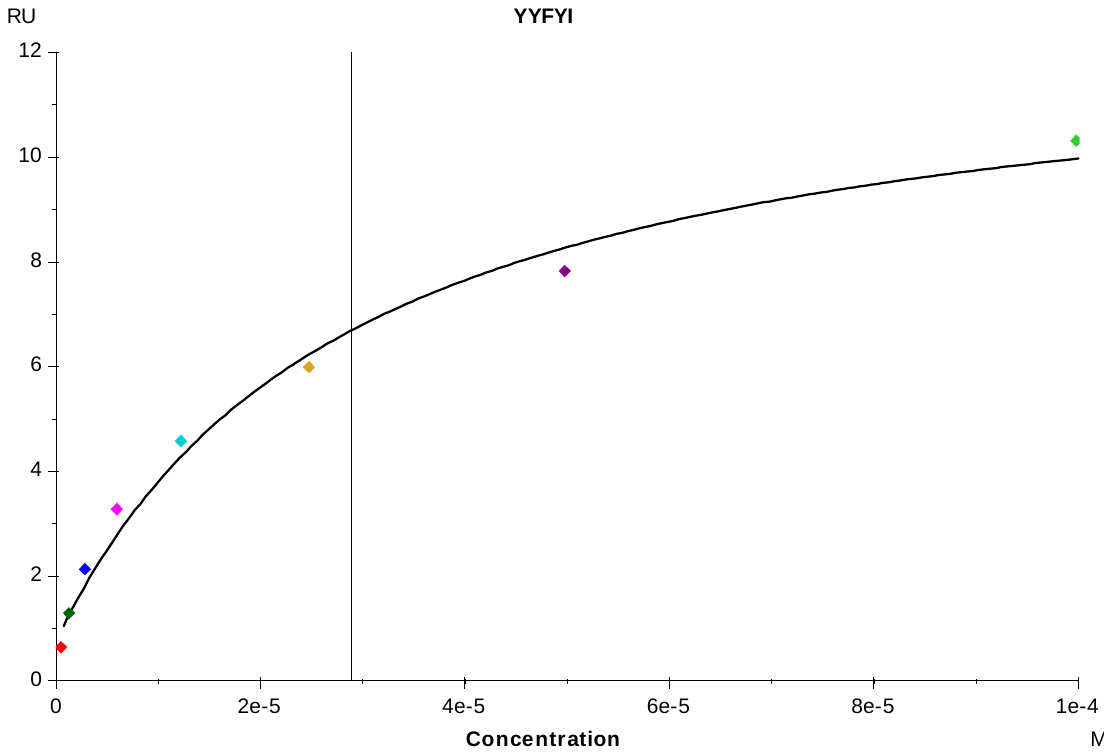

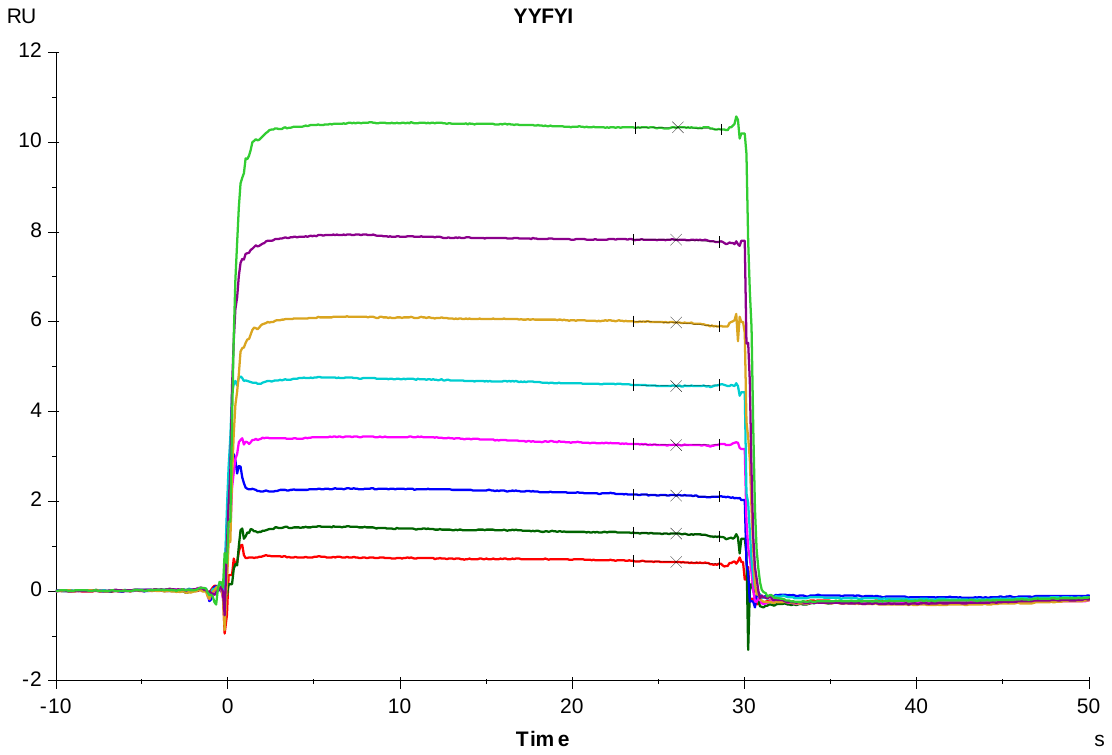

**S6 Fig. Representative sensorgrams (Top) and steady-state binding responses (Bottom) for CymR derived peptides.** (continued)

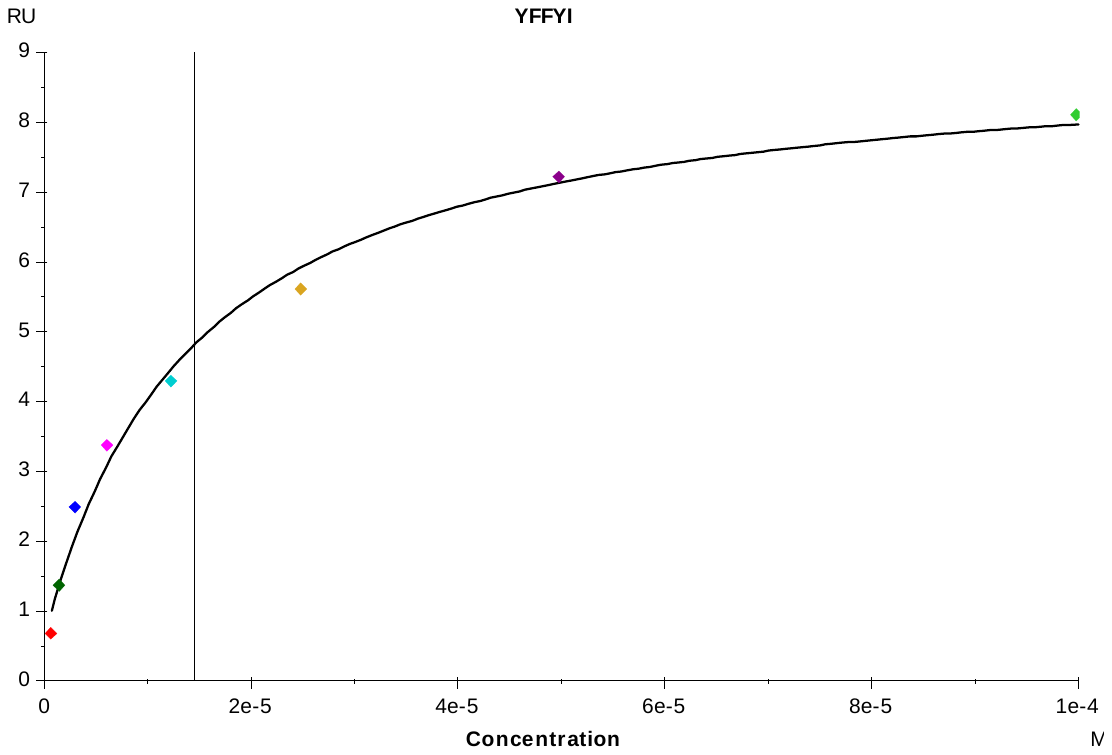

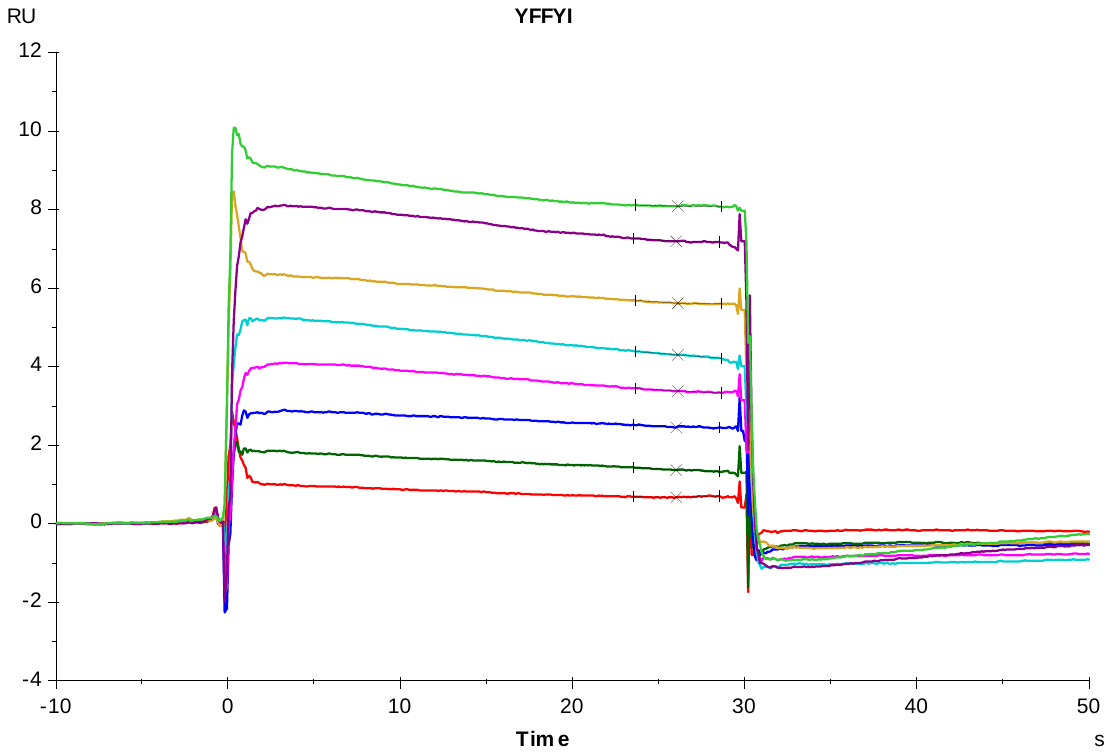

**CymR 5-7b**

**S6 Fig. Representative sensorgrams (Top) and steady-state binding responses (Bottom) for CymR derived peptides.** (continued)

**CymR 5-7c**

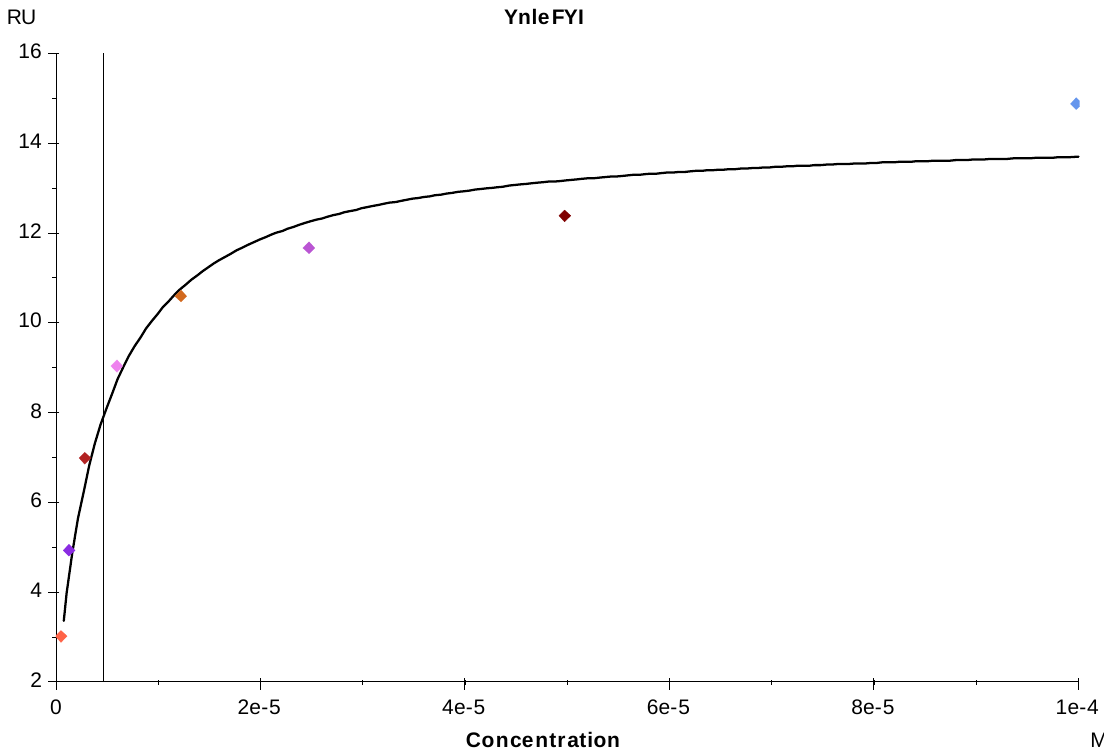

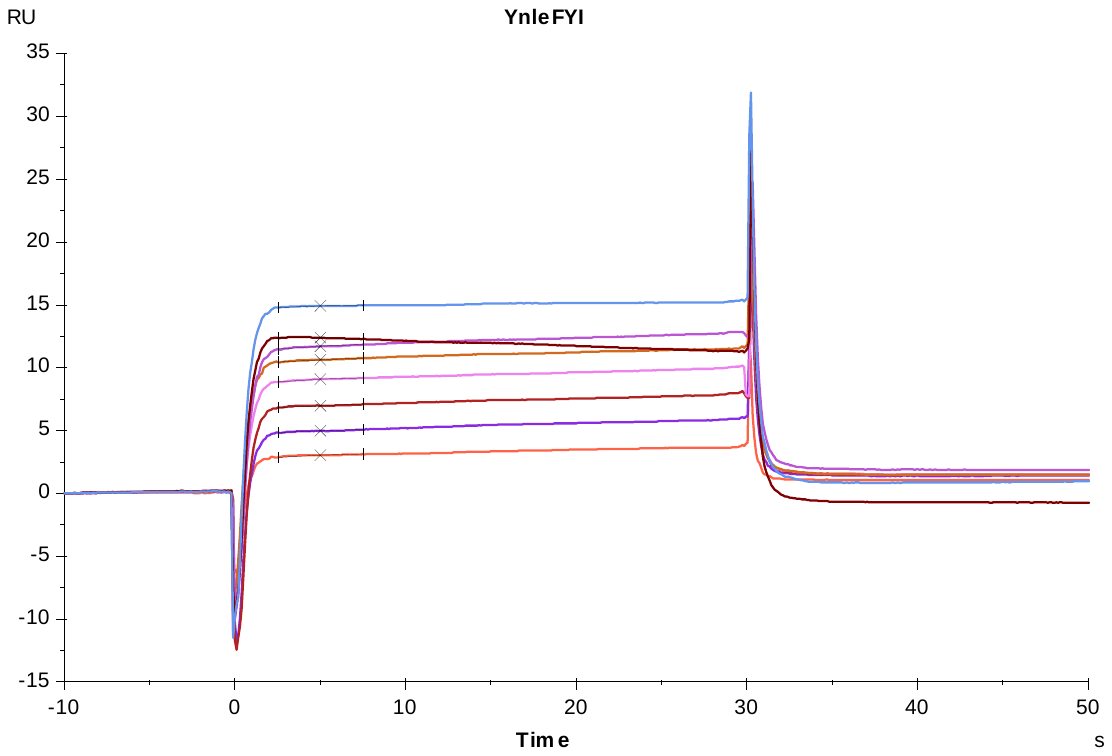

**S6 Fig. Representative sensorgrams (Top) and steady-state binding responses (Bottom) for CymR derived peptides.** (continued)

**CymR 5-7d**

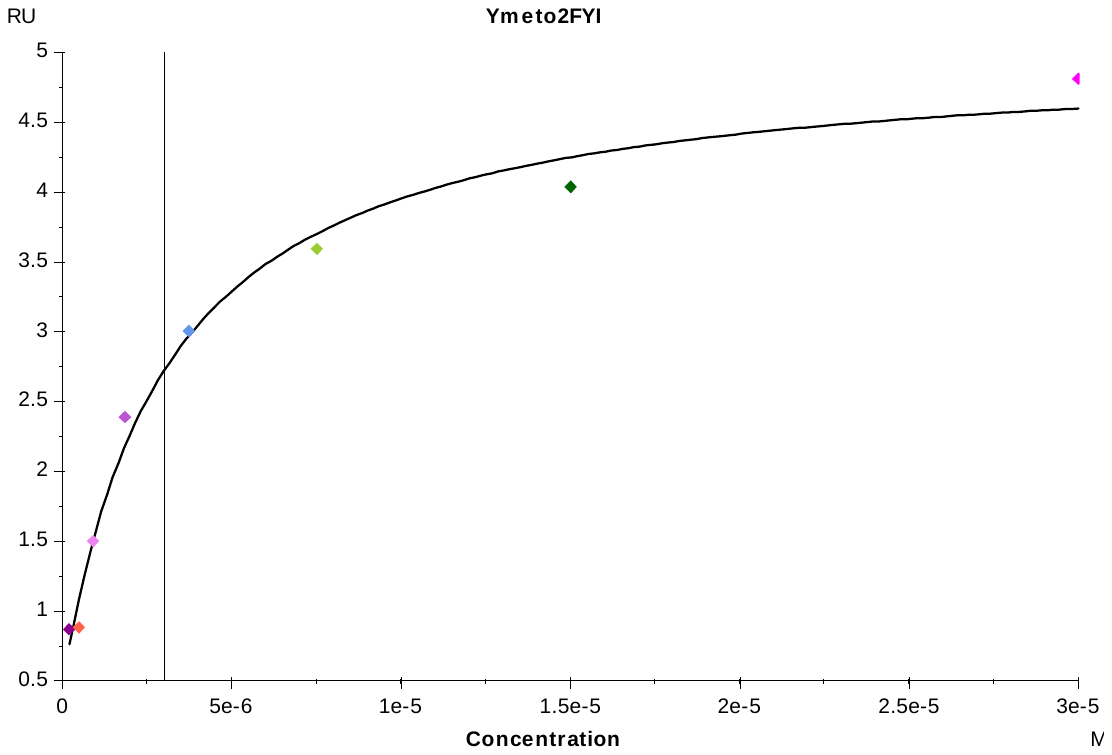

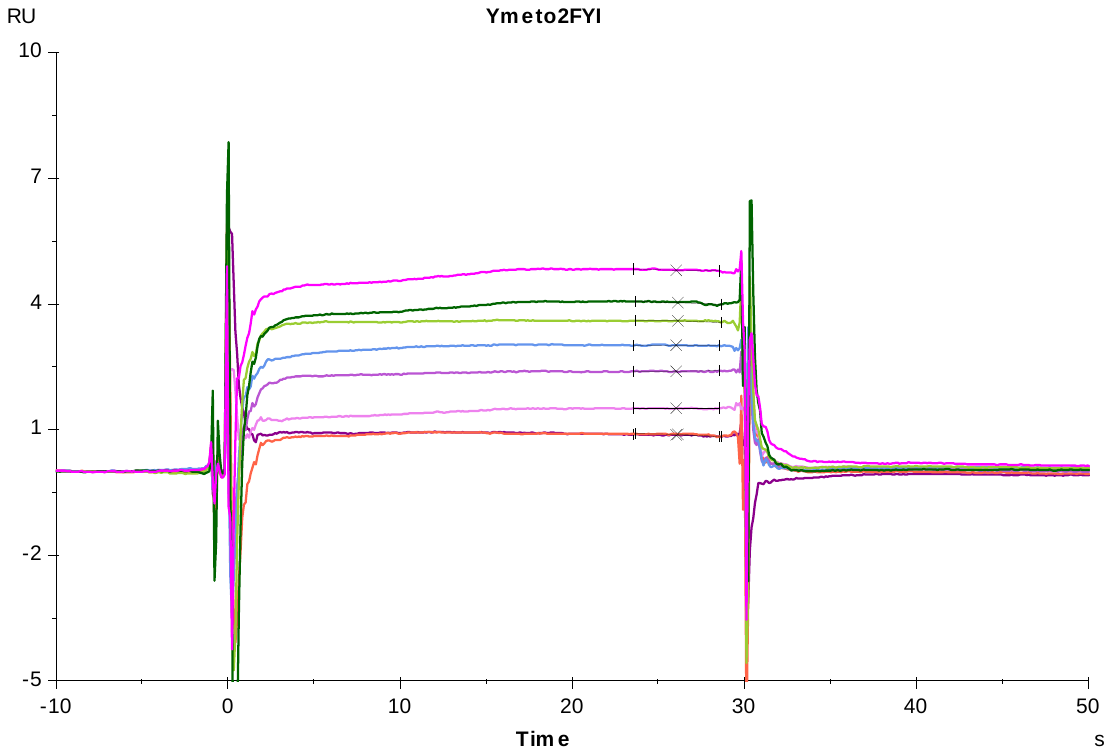

**S6 Fig. Representative sensorgrams (Top) and steady-state binding responses (Bottom) for CymR derived peptides.** (continued)

**CymR 5-8a**

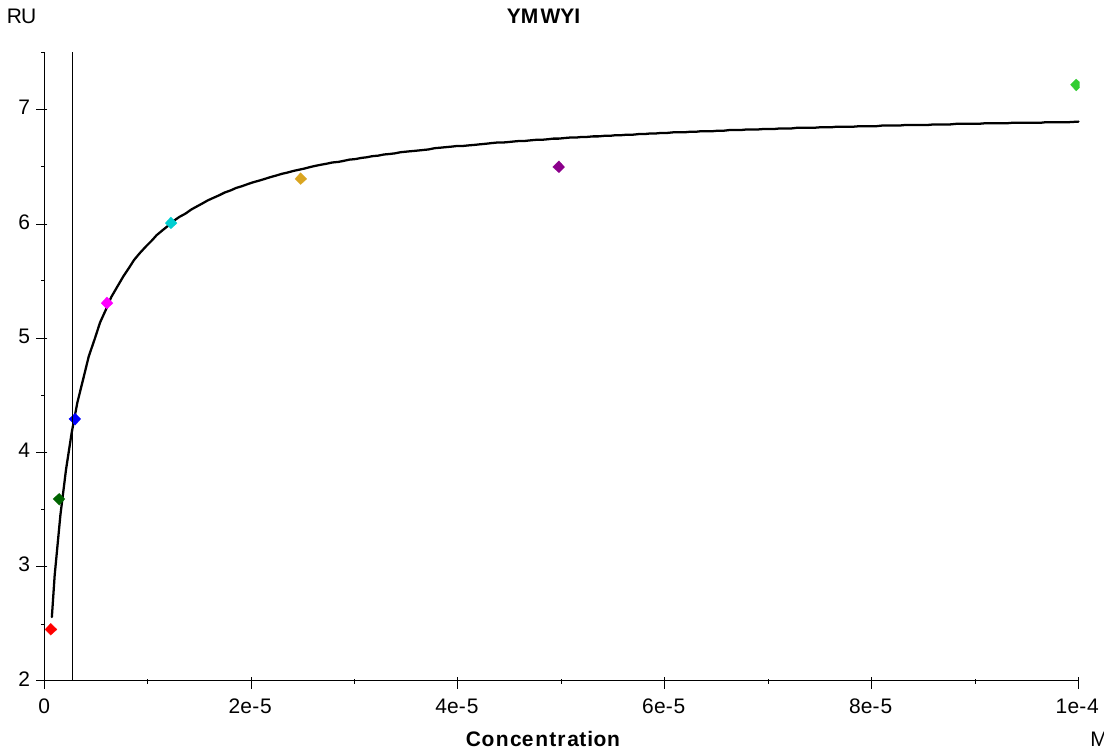

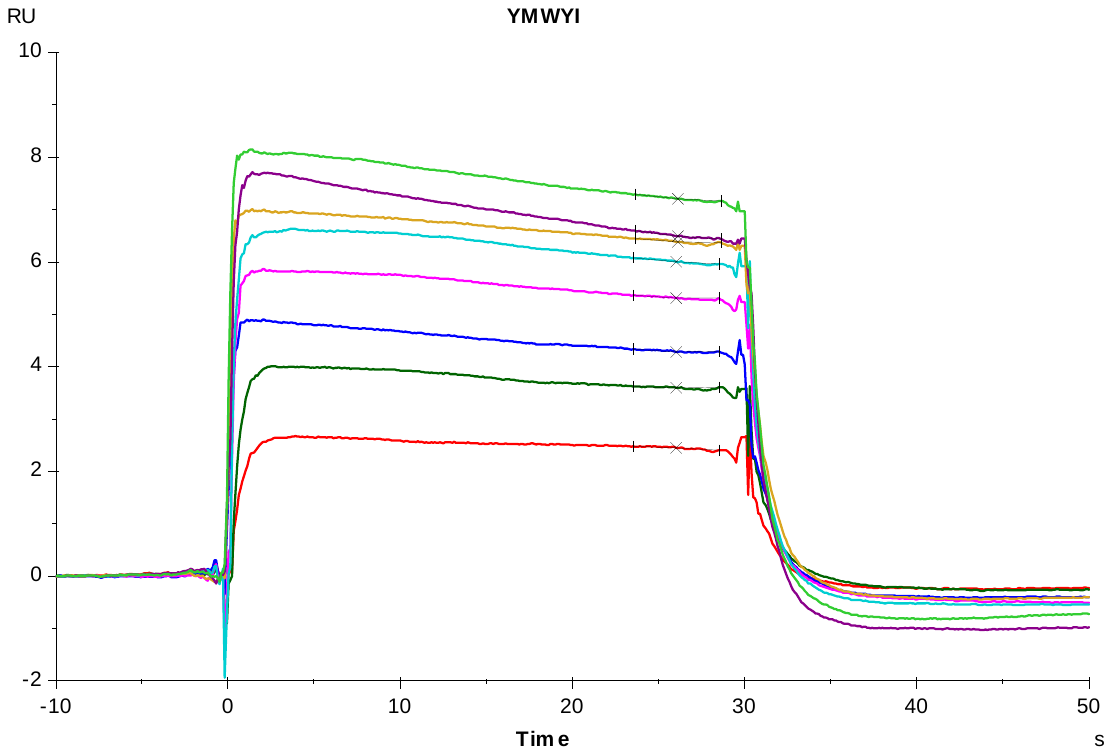

**S6 Fig. Representative sensorgrams (Top) and steady-state binding responses (Bottom) for CymR derived peptides.** (continued)

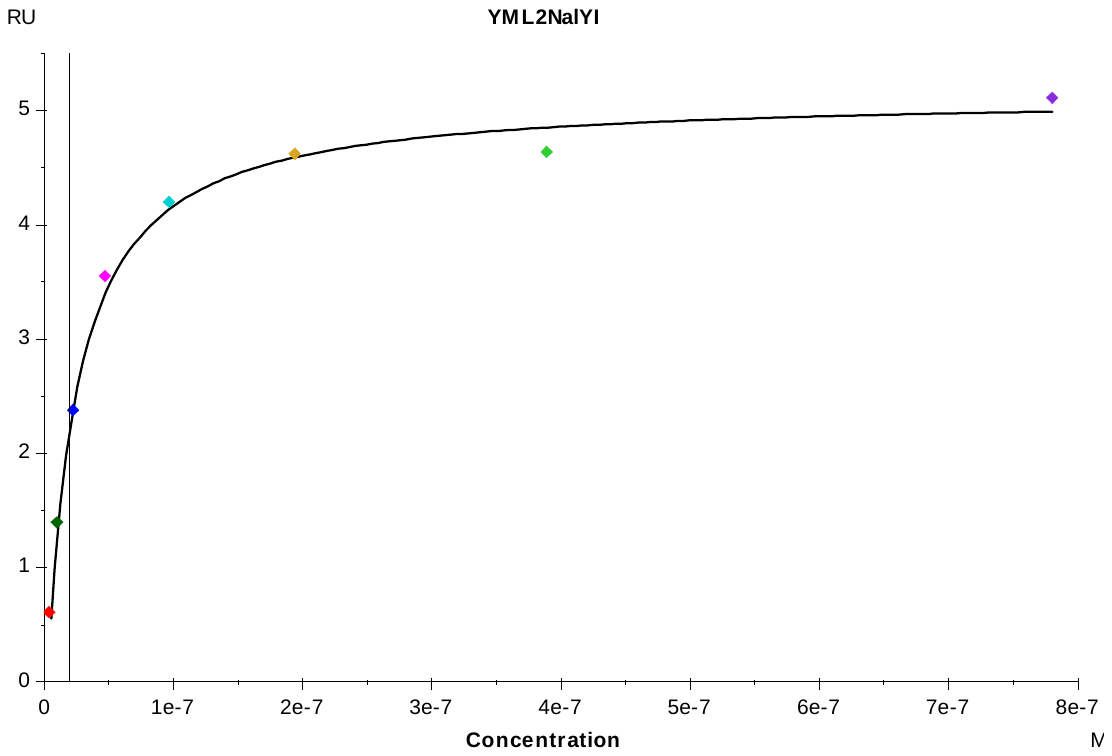

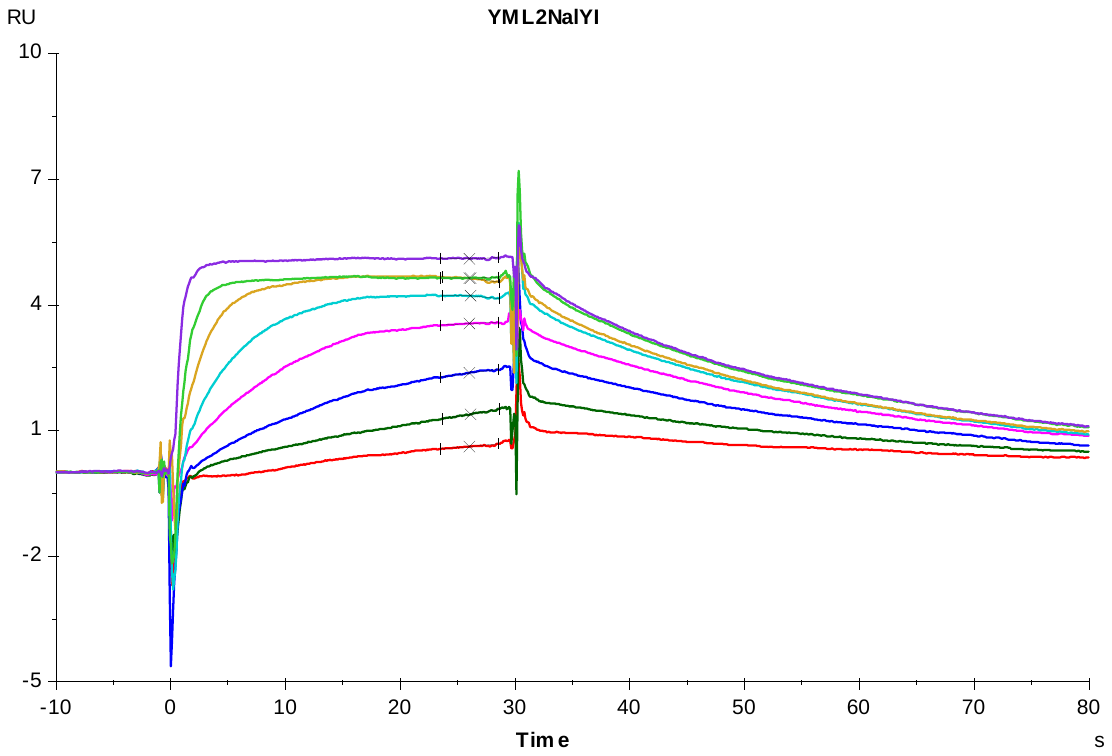

**CymR 5-8b**

**S6 Fig. Representative sensorgrams (Top) and steady-state binding responses (Bottom) for CymR derived peptides.** (continued)

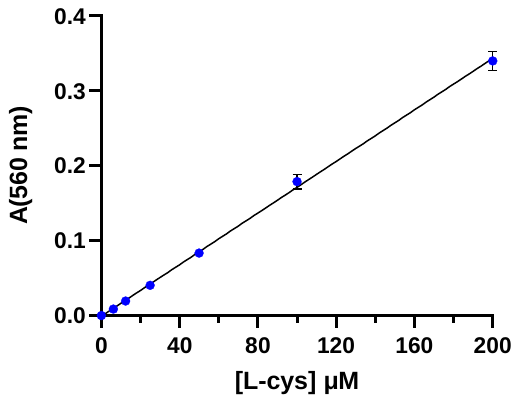

Slope = 0.001720 ± 0.00003

R^2^ = 0.997

**S7 Fig. Standard curve generated for 100 µl samples containing 5 – 200 µM** **ʟ-cys.**

Color development was performed exactly as described in section “Measurement of *Sa*CysK activity”. Samples contained 50 mM Tris-HCl pH 7.5 with the ʟ-cys concentration varied. The slope of the standard curve was used to estimate the *Sa*CysK reaction velocity. Data represents the mean ± standard deviation of two experiments with a single technical replicate for each concentration assayed.
